# Identification and functional characterization of CXCL17 in cartilaginous fishes reveals an ancient origin of the CXCL17–GPR25 signaling pathway

**DOI:** 10.64898/2026.03.04.709523

**Authors:** Jie Yu, Juan-Juan Wang, Hao-Zheng Li, Ya-Li Liu, Zhan-Yun Guo

## Abstract

The newly identified signaling system comprising C-X-C motif chemokine ligand 17 (CXCL17) and G protein-coupled receptor 25 (GPR25) is involved in immune regulation and tumor development. However, the evolutionary origin of this pair has remained unclear because CXCL17 orthologs in lower vertebrates exhibit extreme sequence variation and cannot be identified through conventional homology-based searches. In this study, we identified seven possible CXCL17 orthologs in primitive cartilaginous fishes, including sharks and rays, using an integrated approach based on key amino acid sequence features as well as gene synteny, architecture, and RNA sequencing data in the NCBI gene database. To validate these candidates, a representative ortholog from the cloudy catshark (*Scyliorhinus torazame*), termed St-CXCL17, was prepared via bacterial overexpression and *in vitro* refolding. In cell-based functional assays, St-CXCL17 demonstrated high binding affinity and activation potency toward its corresponding receptor, St-GPR25. Further analysis revealed that removing three conserved C-terminal residues almost completely abolished this activity. While these cartilaginous fish CXCL17s share considerable homology with one another, they lack significant overall similarity to orthologs in mammals, amphibians, or bony fishes. These findings identify functional CXCL17 orthologs in cartilaginous fishes for the first time, implying that the CXCL17–GPR25 signaling pair likely originated in ancient cartilaginous fish ancestors or earlier and has been conserved throughout the evolution of jawed vertebrates.

## 1. Introduction

C-X-C motif chemokine ligand 17 (CXCL17) is a small secreted protein that was first identified two decades ago [1,2]. It is predominantly expressed in human mucosal tissues, including the lung, stomach, salivary glands, and esophagus, where it functions as a chemoattractant that recruits immune cells such as T cells, monocytes, macrophages, and dendritic cells [3–10]. CXCL17 is also expressed in certain tumors and has been associated with tumor progression [11–16], likely through modulation of tumor-associated immune responses. Although designated as “CXCL17” because it contains a C-X-C motif, structural studies indicate that it does not conform to the classical chemokine family, as it adopts a highly flexible structure rather than the compact chemokine fold [17,18].

As a chemoattractant, CXCL17 is expected to interact with a specific plasma membrane receptor; however, the identity of this receptor has long been debated. The orphan G protein-coupled receptor 35 (GPR35) was initially proposed as its receptor [19], but subsequent studies failed to support this interaction [20–22]. A more recent report suggested that CXCL17 modulates the chemokine receptor CXCR4 [22], although this interaction has not yet been independently validated. Additional studies have shown that CXCL17 can act as an agonist for the MAS-related receptors MRGPRX2, MRGPRX1, and MAS1 [23,24]. However, activation of these receptors occurs independently of the conserved C-terminal region of CXCL17 [24], suggesting that they are unlikely to represent evolutionarily conserved receptors for this ligand.

Most recently, Ocón’s team and our group independently demonstrated that CXCL17 activates the rarely studied G protein-coupled receptor 25 (GPR25) through its conserved C-terminal fragment [25–27], suggesting that CXCL17 and GPR25 may constitute an evolutionarily conserved ligand–receptor pair. However, their phylogenetic distributions appear discordant: GPR25 orthologs are widely conserved from fishes to mammals, whereas CXCL17 orthologs initially seemed to be restricted to mammals. To resolve this discrepancy, we previously searched public databases for potential non-mammalian CXCL17 orthologs based on conserved amino acid sequence features as well as gene synteny and gene architecture characteristic of mammalian CXCL17s. Using this strategy, we identified CXCL17 orthologs in amphibians and several bony fishes, including the ray-finned zebrafish (*Danio rerio*) and the lobe-finned coelacanth (*Latimeria chalumnae*) [28–30]. These non-mammalian CXCL17s exhibit little overall amino acid similarity to their mammalian counterparts and therefore remain unrecognized and annotated as hypothetical proteins in public databases.

To further clarify the evolutionary origin of the CXCL17–GPR25 signaling system, we extended our search to more primitive cartilaginous fishes using the same criteria. Seven candidate CXCL17 orthologs were identified, most of which are currently unannotated and display no significant overall amino acid similarity to mammalian, amphibian, or bony fish CXCL17s. Following bacterial expression and purification, a representative CXCL17 from *Scyliorhinus torazame* (cloudy catshark), designated as St-CXCL17, robustly activated its cognate receptor St-GPR25 in cell-based assays, whereas deletion of three C-terminal residues nearly abolished its activity. These findings provide the first functional evidence of CXCL17 orthologs in cartilaginous fishes and suggest that the CXCL17–GPR25 signaling system likely originated in early cartilaginous fishes, or earlier, and was subsequently retained throughout vertebrate evolution.

## 2. Experimental methods

### 2.1. Bacterial overexpression of St-CXCL17

The coding region of the N-terminally 6×His-tagged mature peptide of St-CXCL17 was chemically synthesized at CWBio (Taizhou, Jiangsu province, China) and ligated into a pET vector, resulting in the expression construct pET/6×His-St-CXCL17 (Fig. S1). The expression construct of the C-terminally truncated mutant 6×His-St-[desC3]CXCL17 was generated via the QuikChange approach using pET/6×His-St-CXCL17 as the mutagenesis template. The coding region of the small NanoLuc fragment for NanoBiT (SmBiT)-fused St-CXCL17 was amplified by polymerase chain reaction (PCR) using pET/6×His-St-CXCL17 as the template and ligated into a pET vector via Gibson assembly, resulting in the expression construct pET/6×His-SmBiT-St-CXCL17 (Fig. S1). The St-CXCL17 coding region in these expression constructs was confirmed by DNA sequencing.

Thereafter, the recombinant St-CXCL17 forms were prepared according to our previous procedure developed for other CXCL17s [26–28], including overexpression in *E. coli*, inclusion bodies solubilization via an *S*-sulfonation approach, purification via a Ni^2+^ column, *in vitro* refolding, and further purification by high performance liquid chromatography (HPLC) via C_18_ reverse-phase columns. After lyophilization, the HPLC purified proteins were dissolved in 1.0 mM aqueous hydrochloride (pH 3.0) and quantified by ultra-violet absorbance at 280 nm according to their theoretical extinction coefficient (20970 M^-1^ cm^-1^ for the wild-type (WT) or C-terminally truncated forms; 22460 M^-1^ cm^-1^ for the SmBiT-fused form). Samples at different preparation steps were also analyzed by sodium dodecyl sulfate-polyacrylamide gel electrophoresis (SDS-PAGE). Molecular mass of the refolded WT form was measured by mass spectroscopy conducted on an Orbitrap Exploris 240 mass spectrometer (ThermoFisher Scientific, Waltham, MA, USA).

### 2.2. Generation of the expression constructs for St-GPR25

The coding region of St-GPR25 was chemically synthesized at CWBio (Taizhou) according to its reference cDNA sequence (XM_072480354) and ligated into a pcDNA3.1(+) vector (Table S1 and Fig. S2). Thereafter, the coding region of St-GPR25 was PCR amplified using appropriate primers and ligated into doxycycline (Dox)-inducible vectors for functional assays (Table S1 and Fig. S2). The construct pTRE3G-BI/St-GPR25-LgBiT:SmBiT-ARRB2 coexpresses a C-terminally large NanoLuc fragment (LgBiT)-fused St-GPR25 and an N-terminally SmBiT-fused human β-arrestin 2 (SmBiT-ARRB2) for NanoBiT-based β-arrestin recruitment assays; the construct PB-TRE/sLgBiT-St-GPR25 encodes an N-terminally secretory LgBiT (sLgBiT)-fused St-GPR25 for NanoBiT-based ligand-receptor binding assays; the construct PB-TRE/St-GPR25 encodes an untagged St-GPR25 for calcium mobilization assays and chemotaxis assays.

### 2.3. NanoBiT-based β-arrestin recruitment assays

The NanoBiT-based β-arrestin recruitment assays were conducted on transiently transfected human embryonic kidney (HEK) 293T cells according to our previous procedures for other receptors [26–31]. After cotransfection with pTRE3G-BI/St-GPR25-LgBiT:SmBiT-ARRB2 and expression control plasmid pCMV-Tet3G (Clontech, Mountain View, CA, USA), HEK293T cells were trypsinized, seeded into a white opaque 96-well plate, and cultured in induction medium (complete DMEM plus 1.0 ng/mL of Dox) for ∼24 h to ∼90% confluence. To conduct the β-arrestin recruitment assay, the induction medium was removed, pre-warmed activation solution (serum-free DMEM plus 1% bovine serum albumin) was added (40 μL/well, containing 0.5 μL of NanoLuc substrate stock from Promega, Madison, WI, USA), and bioluminescence data were immediately collected for ∼4 min on a SpectraMax iD3 plate reader (Molecular Devices, Sunnyvale, CA, USA). Thereafter, recombinant St-CXCL17 protein was added (10 μL/well), and bioluminescence data were continuously collected for ∼10 min. The absolute bioluminescence signals were corrected for inter well variability by forcing all curves before addition of agonist to same level and plotted using the SigmaPlot 10.0 software (SYSTAT software, Chicago, IL, USA). To obtain the dose-response curve, the measured bioluminescence data at highest point were plotted with the peptide concentration using the SigmaPlot 10.0 software (SYSTAT software).

### 2.4. NanoBiT-based calcium mobilization assays

A previously designed NanoBiT-based calcium sensor [32] was modified and used in this study for measurement of the cytosolic calcium concentration change in living HEK293T cells. The calcium sensor is composed of a C-terminally SmBiT-fused human calmodulin 1 (CALM1-SmBiT) and a C-terminally human myosin light chain kinase 2 (MYLK2) fragment (L559-K584)-fused LgBiT (LgBiT-MYLK2S) [32]. The DNA fragment encoding LgBiT-MYLK2S was PCR amplified using pNL1.2 (Promega) as template and ligated into the multiple cloning site 1 (MCS1, cleaved by NotI) of the pTRE3G-BI vector (Clontech) via Gibson assembly (Fig. S3). The DNA fragment encoding CALM1-SmBiT was chemically synthesized at CWBio (Taizhou) and ligated into multiple cloning site 2 (MCS2, cleaved by KpnI) of the construct pTRE3G-BI/LgBiT-MYLK2S via Gibson assembly (Fig. S3), resulting in the coexpression construct pTRE3G-BI/LgBiT-MYLK2S:CALM1-SmBiT. The coexpression construct was linearized at 1560 bp position of the pTRE3G-BI vector via PCR amplification, and ligated with the expression frame of the Tet-On 3G transcription factor (PCR amplified from pCMV-Tet3G, 1 bp-2004 bp) via Gibson assembly, resulting in a Dox-response one-vector calcium sensor pTRE-Tet3G/LgBiT-MYLK2S:CALM1-SmBiT.

To measure cytosolic calcium concentration change induced by St-GPR25 activation, the receptor expression construct PB-TRE/St-GPR25 and the one-vector calcium sensor pTRE-Tet3G/LgBiT-MYLK2S:CALM1-SmBiT were cotransfected into HEK293T cells. Next day, the transfected cells were trypsinized, seeded into a white opaque 96-well plate, and cultured in induction medium (complete DMEM plus 20 ng/mL of Dox) for ∼24 h to ∼90% confluence. To conduct the calcium mobilization assay, the induction medium was removed, and pre-warmed activation solution (phosphate-buffered saline plus 1% bovine serum albumin) was added (40 μL/well, containing 0.5 μL of NanoLuc substrate stock). After incubation at room temperature for 10 min, bioluminescence data were collected for ∼4 min on a SpectraMax iD3 plate reader (Molecular Devices). Thereafter, recombinant St-CXCL17 protein was added (10 μL/well, diluted in activation solution), and bioluminescence data were continuously collected for ∼10 min. The absolute bioluminescence signals were corrected for inter well variability by forcing all curves before addition of agonist to same level and plotted using the SigmaPlot 10.0 software (SYSTAT software). To obtain the dose-response curve, the measured bioluminescence data at highest point were plotted with the peptide concentration using the SigmaPlot 10.0 software (SYSTAT software).

### 2.5. Homogenous ligand-receptor binding assays

The NanoBiT-based homogenous ligand-receptor binding assays were conducted on transiently transfected HEK293T cells according to our previous procedures for other receptors [28,29,33,34]. HEK293T cells were transfected with PB-TRE/sLgBiT-St-GPR25 with or without cotransfection of the human tyrosylprotein sulfotransferases expression construct pTRE3G-BI/TPST1:TPST2. Next day, the transfected cells were trypsinized, seeded into white opaque 96-well plates, and cultured in induction medium (complete DMEM plus 20 ng/mL of Dox) for ∼24 h to ∼90% confluence. To conduct the binding assay, the induction medium was removed, and pre-warmed binding solution (serum-free DMEM plus 0.1% bovine serum albumin and 0.01% Tween-20) was added (50 μL/well). For saturation binding assays, the binding solution contains different concentrations of 6×His-SmBiT-St-CXCL17. For competition binding assays, the binding solution contains a constant concentration of 6×His-SmBiT-St-CXCL17 and different concentrations of competitors. After incubation at room temperature for ∼1 h, diluted NanoLuc substrate (30-fold dilution in the binding solution) was added (10 μL/well), and bioluminescence was immediately measured on a SpectraMax iD3 plate reader (Molecular Devices). The measured bioluminescence data were expressed as mean ± standard deviation (SD, *n* = 3) and fitted to one-site binging model using the SigmaPlot 10.0 software (SYSTAT software).

## 3. Results

### 3.1. Identification of possible CXCL17s from cartilaginous fishes

To identify potential CXCL17 orthologs in cartilaginous fishes, we retrieved secretory proteins shorter than 200 residues from the UniProt database. Among 2,670 candidates, we identified a hypothetical protein, scyTo_0013081 (UniProt ID: A0A401NNW6), from the cloudy catshark (*Scyliorhinus torazame*). This sequence was also detectable in the NCBI protein database (GCB62570) by BLAST analysis. Although derived from the early draft genome (Storazame_v1.0) of *S. torazame* [35], it was not annotated in the most recent reference genome (sScyTor2.1). Examination of the updated genome assembly revealed clear transcriptional evidence based on RNA sequencing data (Fig. S4A). The transcript spans approximately 40 kb on chromosome 12 and consists of four exons and three introns (Fig. S4A,B), a gene architecture conserved in mammalian, amphibian, and bony fish CXCL17 genes. Manual assembly of the four exons generated a full-length cDNA encoding scyTo_0013081 (Table S2 and Fig. S4C,D), confirming its expression despite the lack of annotation. For clarity, scyTo_0013081 has been designated as St-CXCL17 for this study.

St-CXCL17 possesses key structural features of CXCL17 orthologs, including an N-terminal signal peptide and two partially overlapping Xaa-Pro-Yaa motifs at the C-terminus (Fig. 1A and Table S2). However, it displays no significant overall amino acid sequence similarity to known CXCL17 orthologs from mammals, amphibians, or bony fishes (Fig. 1A), including human CXCL17 (Hs-CXCL17), zebrafish CXCL17s (Dr-CXCL17, Dr-CXCL17L), and coelacanth CXCL17 (Lc-CXCL17a), which explains why it cannot be identified through conventional homology-based approaches. A distinctive feature of St-CXCL17 is that its mature peptide contains eight cysteine residues, in contrast to the six or four cysteines typically found in mammalian, amphibian, or bony fish orthologs (Fig. 1A). This suggests that mature St-CXCL17 forms four intramolecular disulfide bonds. While CXCL17 sequences show high variation, GPR25 orthologs are much more evolutionarily conserved; for example, the GPR25 ortholog from *S. torazame* (St-GPR25) shares considerable sequence similarity with its human and bony fish counterparts (Fig. 1B).

**Fig. 1.**
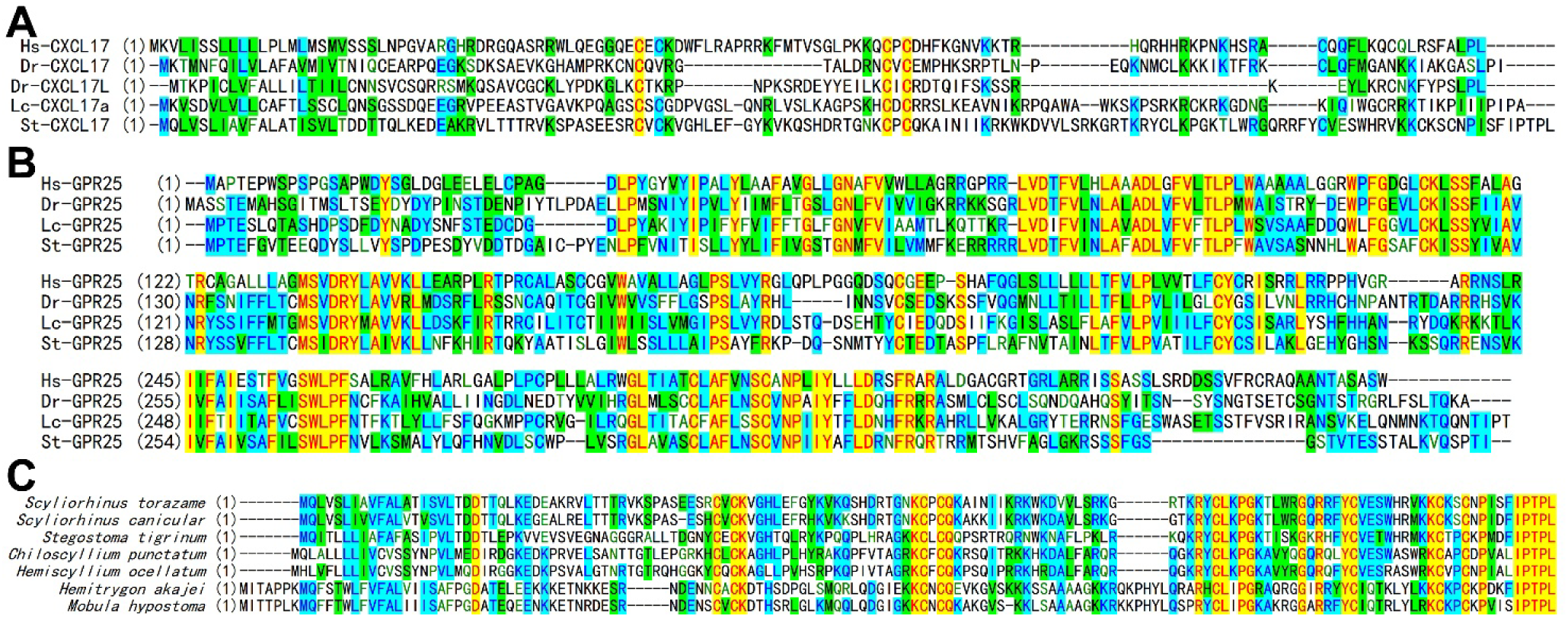
Amino acid sequence alignment of CXCL17 orthologs and GPR25 orthologs. (**A,B**) Amino acid sequence alignment of CXCL17s (A) or GPR25s (B) from cloudy catshark (*S. torazame*), zebrafish (*D. rerio*), coelacanth (*L. chalumnae*), and human (*H. sapiens*). (**C**) Amino acid sequence alignment of the newly identified cartilaginous fish CXCL17s. Information about these proteins is listed in the supplementary Table S2, S3 and Fig. S4-S10.

No homologs were retrieved from the NCBI database via BLAST search using St-CXCL17 as a query (https://www.ncbi.nlm.nih.gov/blast), suggesting that other cartilaginous fish CXCL17 orthologs likely remain unannotated. The *st-cxcl17* gene is flanked by *V-set and immunoglobulin domain-containing protein 10-like* (LOC140386591), *cell adhesion molecule CEACAM5-like* (LOC140386590), *lipeb*, and *tmem145*, and it consists of four exons and three introns (Fig. S4A).

Hypothesizing that *cxcl17* orthologs in other cartilaginous fishes would share this genomic architecture and proximity to these marker genes, we screened the NCBI gene database based on available RNA sequencing data. This approach identified unannotated CXCL17 candidates in six additional cartilaginous fish species: *Scyliorhinus canicular* (smaller spotted catshark), *Stegostoma tigrinum*, *Chiloscyllium punctatum* (brownbanded bambooshark), *Hemiscyllium ocellatum* (epaulette shark), *Hemitrygon akajei* (red stingray), and *Mobula hypostoma* (lesser devil ray) (Fig. S5–S10). Upon manual assembly of these transcript exons, the resulting cDNAs were found to encode putative CXCL17 proteins featuring an N-terminal signal peptide, eight cysteine residues (forming a CXC-CXC-C-C-C-C motif) within the mature peptide, and two partially overlapping Xaa-Pro-Yaa motifs at the C-terminus (Fig. 1C and Table S2). These orthologs exhibit considerable sequence similarity, particularly at their C-terminal residues (Fig. 1C), suggesting a common ancestry and a slow evolutionary rate.

The mature cartilaginous fish CXCL17s are all highly basic proteins, with calculated isoelectric points (pI) ranging from 10.2 to 10.9. This alkalinity likely facilitates their function as chemoattractant, as it allows them to bind negatively charged extracellular glycosaminoglycans upon secretion, thereby establishing the local concentration gradients necessary to recruit immune cells. Structural predictions using the AlphaFold3 algorithm yielded pTM values of approximately 0.2, indicating that the mature proteins possess highly flexible conformations. However, the algorithm predicted their binding to corresponding GPR25 receptors with ipTM values of approximately 0.5, which is slightly below the standard reliability threshold of 0.6. In these predicted models, the C-terminus inserts into the orthosteric ligand-binding pocket of the GPR25 receptor, consistent with the high degree of conservation observed in these C-terminal residues.

### 3.2. Preparation of recombinant St-CXCL17 for functional assays

To determine whether cartilaginous fish CXCL17s are functionally active, St-CXCL17 from *S. torazame* was selected as a representative and produced by bacterial overexpression. Following induction of transformed *E. coli* with isopropyl-β-D-thiogalactopyranoside (IPTG), a prominent band slightly below 20 kDa appeared on SDS-PAGE (Fig. 2A, asterisk), indicating successful expression of 6×His-St-CXCL17. After sonication, the recombinant protein was predominantly detected in the pellet fraction (Fig. 2A), consistent with its accumulation in inclusion bodies. The inclusion bodies were solubilized using an *S*-sulfonation strategy, and the protein was subsequently purified by Ni^2+^ affinity chromatography. SDS-PAGE analysis of the eluted fraction revealed the expected monomeric band (indicated by an asterisk) along with higher-molecular-weight species (Fig. 2A), suggesting a tendency toward intermolecular crosslinking, as previously reported for other CXCL17 orthologs [26,28–30].

**Fig. 2.**
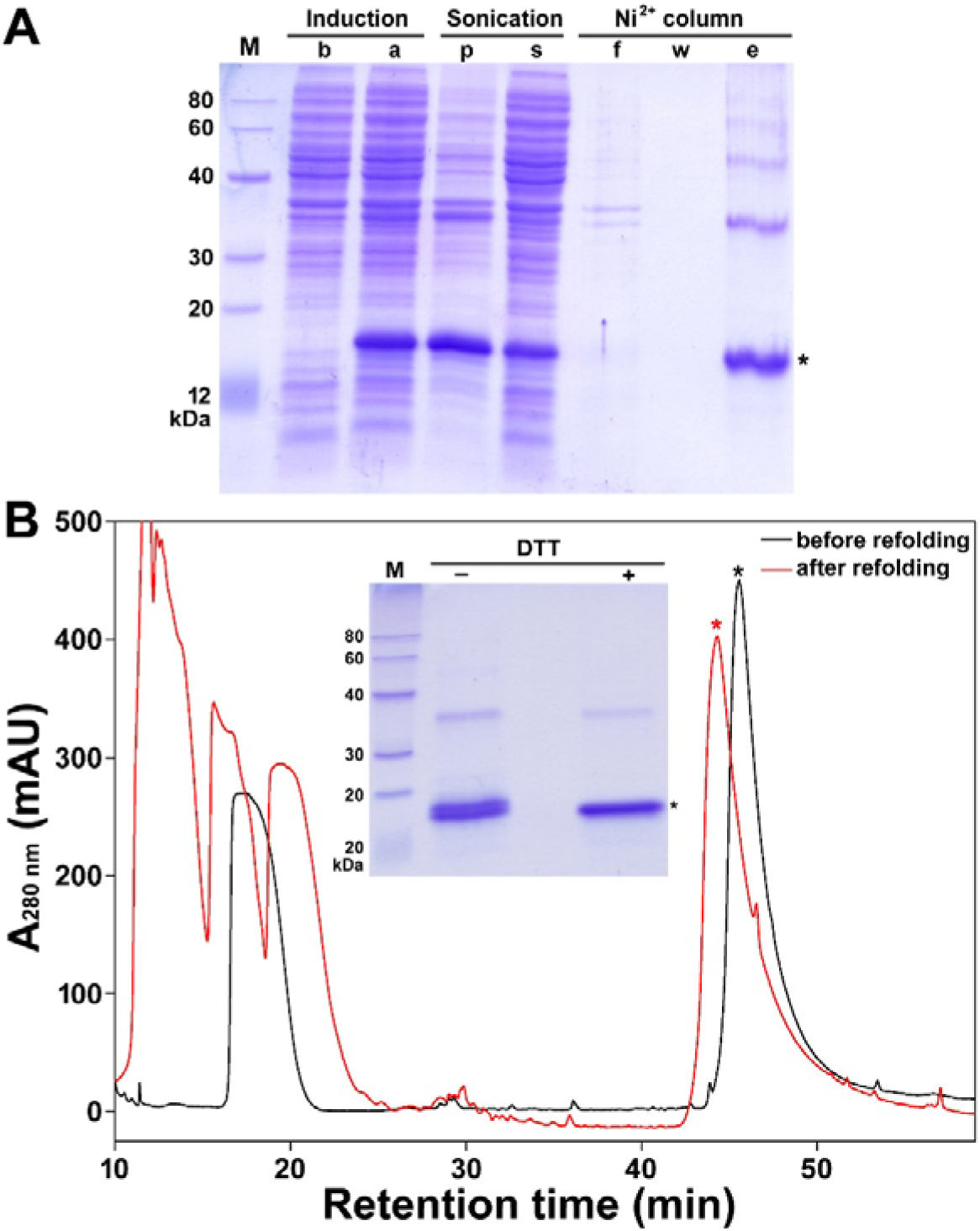
Recombinant preparation of St-CXCL17. (**A**) SDS-PAGE analysis of the recombinant 6×His-St-CXCL17 at different stages. Lane (M), protein ladder; lane (b), before IPTG induction; lane (a), after IPTG induction; lane (p), pellet after sonication; lane (s), supernatant after sonication; lane (f), flow-through fraction; lane (w), washing fraction; lane (e), eluted fraction. After electrophoresis, the SDS-gel was stained by Coomassie brilliant blue R250. Band of the monomeric 6×His-St-CXCL17 is indicated by an asterisk. (**B**) HPLC analysis of the recombinant 6×His-St-CXCL17 before or after *in vitro* refolding. **Inner panel**, SDS-PAGE analysis of the refolded 6×His-St-CXCL17. Lane (M), protein ladder; lane (□), without DTT treatment; lane (+), with DTT treatment. The SDS-gel was stained by Coomassie brilliant blue R250 after electrophoresis. Band of the monomeric form was indicated by an asterisk.

HPLC analysis showed a dominant peak corresponding to 6×His-St-CXCL17 both before and after *in vitro* refolding (Fig. 2B). SDS-PAGE of the refolded protein revealed two closely migrating monomeric bands and a faint dimer band (Fig. 2B, inner panel). The two monomer bands likely represent disulfide isomers, as they merged into a single band following dithiothreitol (DTT) treatment. In contrast, the dimer band was resistant to DTT, indicating possible intermolecular crosslinking through non-disulfide bonds. Mass spectrometry analysis confirmed an average molecular mass of 15,071.3 Da, in excellent agreement with the theoretical value (15,071.4 Da), verifying correct refolding and molecular integrity of the recombinant protein.

### 3.3. Activation of St-GPR25 by recombinant St-CXCL17 measured via β-arrestin recruitment

To evaluate whether recombinant St-CXCL17 activates its cognate receptor, we employed a NanoBiT-based β-arrestin recruitment assay in transfected HEK293T cells coexpressing C-terminally LgBiT-fused St-GPR25 (St-GPR25-LgBiT) and N-terminally SmBiT-fused human β-arrestin 2 (SmBiT-ARRB2). Upon ligand-induced receptor activation, recruitment of SmBiT-ARRB2 to St-GPR25-LgBiT restores NanoLuc activity through proximity-driven complementation of SmBiT and LgBiT, resulting in increased bioluminescence.

Following induction of receptor and β-arrestin expression, addition of NanoLuc substrate alone produced only a minimal increase in bioluminescence. In contrast, subsequent addition of 6×His-St-CXCL17 triggered a rapid and dose-dependent increase in signal, with a calculated EC_50_ of approximately 60 nM (Fig. 3A), demonstrating that St-CXCL17 effectively activates St-GPR25 and promotes β-arrestin recruitment. Specifically, the C-terminally truncated mutant 6×His-St-[desC3]CXCL17 exhibited a dramatic loss of activity (Fig. 3B), indicating that the conserved C-terminal residues are critical for receptor activation.

**Fig. 3.**
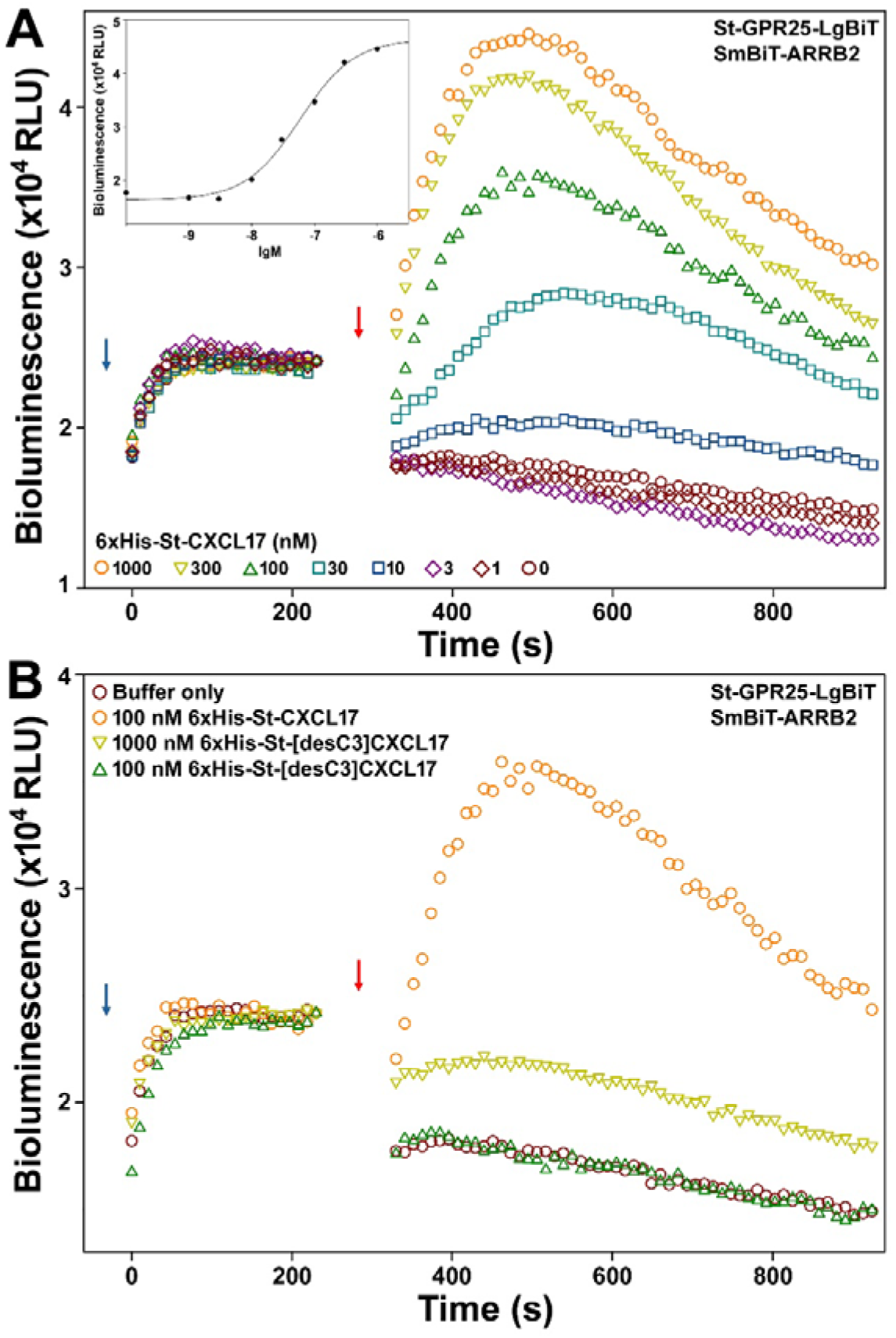
Activation of St-GPR25 by recombinant St-CXCL17 measured via NanoBiT-based β-arrestin recruitment assays. (**A**) Bioluminescence change induced by WT 6×His-St-CXCL17. **Inner panel**, Dose-response curve. (**B**) Bioluminescence change induced by the C-terminally truncated 6×His-St-[desC3]CXCL17. The β-arrestin recruitment assays were conducted on living HEK293T cells coexpressing St-GPR25-LgBiT and SmBiT-ARRB2 by sequential addition of NanoLuc substrate and different concentrations of indicated WT or mutant peptide. Blue arrows indicate the addition of NanoLuc substrate, and red arrows indicate the addition of peptide.

### 3.4. Activation of St-GPR25 by recombinant St-CXCL17 measured via calcium mobilization

To further assess downstream signaling, we monitored cytosolic calcium changes using a NanoBiT-based calcium sensor composed of a SmBiT-fused human calmodulin 1 (CALM1-SmBiT) and a human myosin light chain kinase 2 (MYLK2) fragment (L559-K584)-fused LgBiT (LgBiT-MYLK2S) [31]. Elevation of intracellular calcium induces conformational changes in CALM1-SmBiT, enabling its interaction with LgBiT-MYLK2S and restoring NanoLuc activity through fragment complementation.

In HEK293T cells coexpressing St-GPR25 and the calcium sensor, addition of NanoLuc substrate produced only background-level bioluminescence. Subsequent stimulation with 6×His-St-CXCL17 resulted in a rapid, dose-dependent increase in signal, with an EC_50_ of approximately 140 nM (Fig. 4A). These results indicate that St-GPR25 activation triggers intracellular calcium mobilization, consistent with downstream G protein signaling. In contrast, 6×His-St-[desC3]CXCL17 produced minimal response in this assay (Fig. 4B), in agreement with its markedly reduced activity in the β-arrestin recruitment assay.

**Fig. 4.**
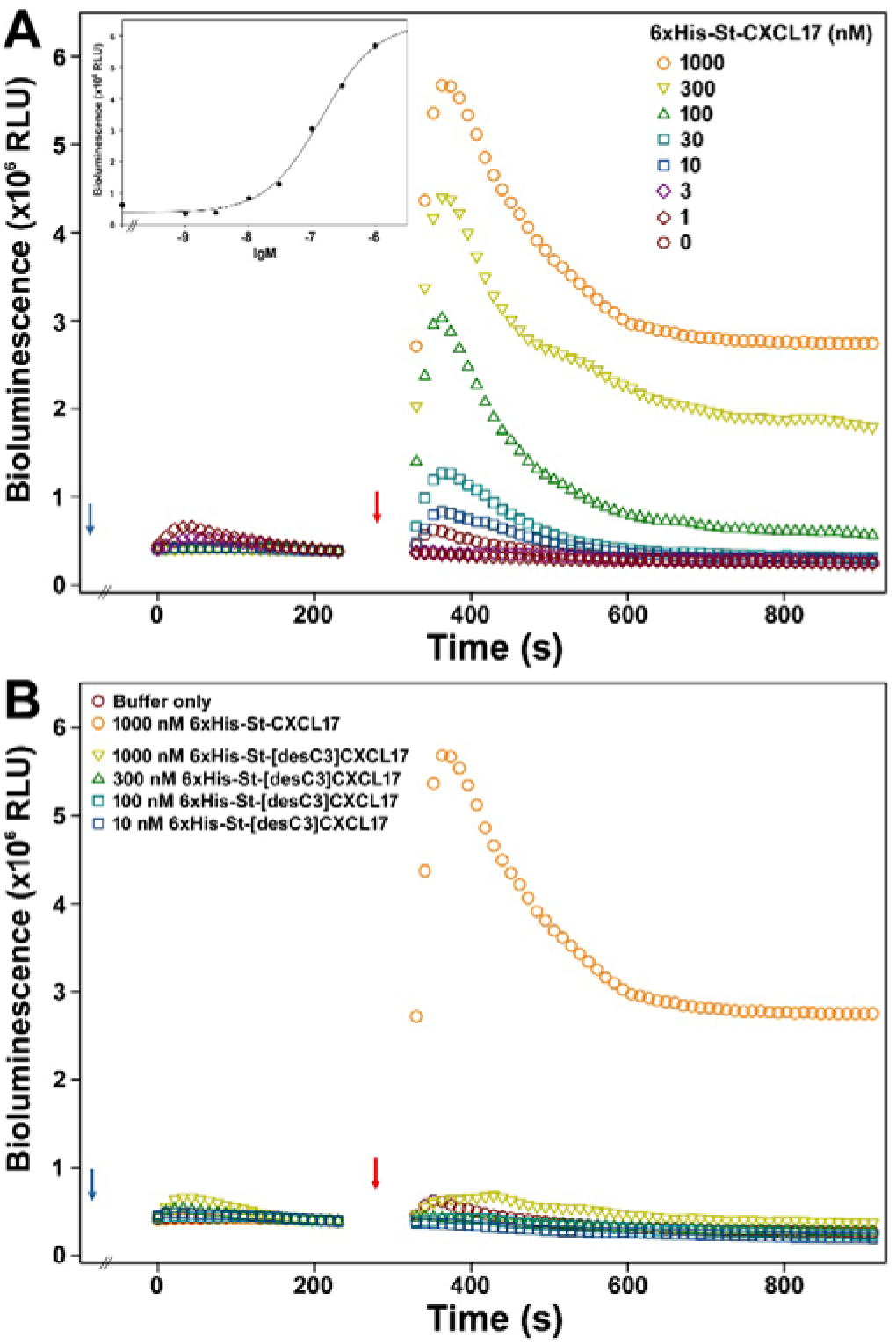
Activation of St-GPR25 by recombinant St-CXCL17 measured via NanoBiT-based calcium mobilization assays. (**A**) Bioluminescence change induced by WT 6×His-St-CXCL17. **Inner panel**, Dose-response curve. (**B**) Bioluminescence change induced by the C-terminally truncated 6×His-St-[desC3]CXCL17. The calcium mobilization assays were conducted on living HEK293T cells coexpressing St-GPR25 and the NanoBiT-based cytosolic calcium sensor by sequential addition of NanoLuc substrate and different concentrations of indicated WT or mutant peptide. Blue arrows indicate the addition of NanoLuc substrate, and red arrows indicate the addition of peptide.

### 3.5. Direct binding of recombinant St-CXCL17 to St-GPR25

To examine direct ligand–receptor interaction, we performed a NanoBiT-based homogenous binding assay using a SmBiT-tagged ligand and an N-terminally sLgBiT-fused receptor [28–32]. For tracer preparation, a 6×His-SmBiT tag was fused to the N-terminus of St-CXCL17 (Fig. S1). Following bacterial expression and purification, 6×His-SmBiT-St-CXCL17 retained the ability to activate St-GPR25 in the β-arrestin recruitment assay, with an EC_50_ of approximately 150 nM (Fig. 5A), indicating that the SmBiT tag did not substantially impair biological activity.

**Fig. 5.**
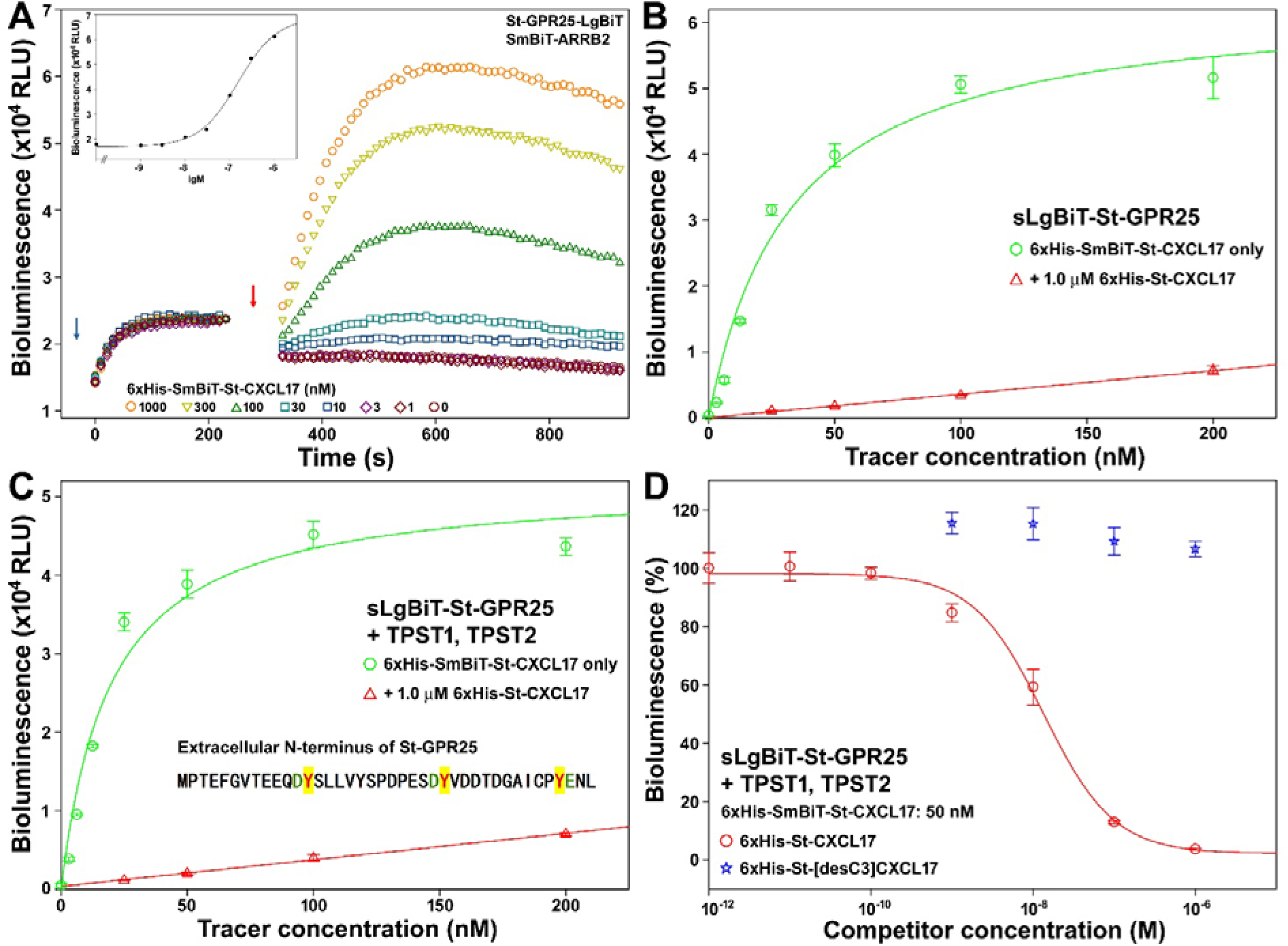
Direct binding of recombinant St-CXCL17 with St-GPR25 measured via NanoBiT-based homogenous binding assays. (**A**) Activity of 6×His-SmBiT-St-CXCL17 towards St-GPR25 measured via β-arrestin recruitment assay. **Inner panel**, dose-response curve. NanoLuc substrate and different concentrations of 6×His-SmBiT-St-CXCL17 were sequentially added to living HEK293T cells coexpressing St-GPR25-LgBiT and SmBiT-ARRB2, and bioluminescence was continuously measured on a plate reader. Blue arrow indicates the addition of NanoLuc substrate, and red arrow indicates the addition of peptide. (**B,C**) Saturation binding of 6×His-SmBiT-St-CXCL17 with sLgBiT-St-GPR25 without (B) or with (C) coexpression of human TPST1 and TPST2. The bioluminescence data are expressed as mean ± SD (*n* = 3) and plotted using the SigmaPlot10.0 software. Total binding data (green circles) were fitted with the function of Y = B_max_X/(K_d_+X), the non-specific binding data (red triangles) were fitted with linear curves. (**D**) The NanoBiT-based competition binding assays. The relative bioluminescence data are expressed as mean ± SD (*n* = 3) and fitted with sigmoidal curves using the SigmaPlot10.0 software.

When 6×His-SmBiT-St-CXCL17 was applied to HEK293T cells overexpressing sLgBiT-St-GPR25, bioluminescence increased in a saturable manner (Fig. 5B), yielding a dissociation constant (K_d_) of 33.0 ± 4.1 nM (*n* = 3). The signal was markedly reduced in the presence of excess unlabeled 6×His-St-CXCL17 (1.0 μM), confirming specific binding (Fig. 5B). The N-terminus of St-GPR25 contains three Tyr residues adjacent to negatively charged residues (Fig. 5C), suggesting potential tyrosine sulfation. Coexpression of human TPST1 and TPST2 modestly enhanced binding affinity, reducing the K_d_ to 19.5 ± 2.4 nM (*n* = 3) (Fig. 5C), indicating that N-terminal sulfation slightly strengthens ligand–receptor interaction.

In competition binding assays, 6×His-St-CXCL17 inhibited tracer binding in a sigmoidal manner, with an IC_50_ of 13.5 ± 1.8 nM (n = 3) (Fig. 5D). In contrast, 6×His-St-[desC3]CXCL17 failed to compete (Fig. 5D), demonstrating that the conserved C-terminal residues are critical for St-CXCL17 binding to St-GPR25.

## Discussion

In this study, we identified and functionally characterized CXCL17 orthologs in primitive cartilaginous fishes for the first time, thereby tracing the origin of the CXCL17–GPR25 signaling system to early cartilaginous fish ancestors. These CXCL17 sequences show no significant overall amino acid similarity to CXCL17 orthologs from mammals, amphibians, or bony fishes [26,28,29,35], and most were previously unannotated in public databases. Their identification was achieved through an integrated strategy that combined analysis of conserved amino acid features, gene synteny and genomic architecture, and RNA sequencing evidence from reference genomes. This approach proved essential for detecting highly divergent orthologs that cannot be recognized by conventional homology-based methods. With the continued improvement of genome assemblies and transcriptomic datasets, application of this strategy is likely to facilitate the discovery of additional CXCL17 orthologs across diverse vertebrate lineages.

These cartilaginous fish CXCL17 orthologs share considerable sequence homology with each other, indicating derivation from a common ancestor and relatively slow evolutionary divergence, consistent with the generally slow molecular evolution of cartilaginous fishes. When compared with known CXCL17s from mammals, amphibians, and bony fishes [26,28–30], several distinctive features become apparent. First, all cartilaginous fish CXCL17s contain eight cysteine residues in their mature peptides, suggesting the formation of four intramolecular disulfide bonds. In contrast, mammalian and bony fish CXCL17s typically possess six cysteine residues [26,28,29], whereas most amphibian CXCL17s contain only four [30]. Second, although all known CXCL17s harbor a conserved C-terminal Xaa-Pro-Yaa motif, in which Xaa and Yaa are usually large aliphatic residues such as leucine, isoleucine, methionine, or valine in mammals, amphibians, and bony fishes [26,28–30], the corresponding motif in cartilaginous fish CXCL17s uniquely includes a hydrophilic threonine residue (Fig. 1C). These structural distinctions expand our understanding of the core features and permissible sequence variability of CXCL17 orthologs and provide valuable criteria for identifying additional divergent members of this ligand family in future investigations.

In transfected HEK293T cells, St-CXCL17 directly bound to and efficiently activated St-GPR25, as demonstrated by NanoBiT-based ligand–receptor binding, β-arrestin recruitment, and calcium mobilization assays. However, St-CXCL17 did not induce chemotactic migration of HEK293T cells expressing St-GPR25 (data not shown), whereas other non-mammalian CXCL17–GPR25 pairs previously examined displayed this capability [28–30]. One possible explanation is that components of the intracellular signaling machinery in human-derived HEK293T cells are not fully compatible with St-GPR25, thereby limiting downstream pathways required for cell migration. In native cartilaginous fish cells, the CXCL17–GPR25 pair is likely to mediate chemotactic responses, although this remains to be experimentally verified. By establishing the presence and functional activity of CXCL17 orthologs in cartilaginous fishes, the present study lays a foundation for future investigations into the physiological roles of the CXCL17–GPR25 signaling system in this primitive vertebrate lineage.

## CRediT authorship contribution statement

Jie Yu: Investigation, Methodology, Visualization. Juan-Juan Wang: Investigation, Methodology. Hao-Zheng Li: Investigation, Methodology. Ya-Li Liu: Project administration. Zhan-Yun Guo: Supervision, Conceptualization, Writing - Original Draft, Writing - Review & Editing, Funding acquisition.

## Declaration of competing interest

The authors declare no competing interests.

## Acknowledgments

This work was supported by grant from the National Natural Science Foundation of China (31971193).

## Supplementary materials

Supplementary materials for this article can be found online (Table S1-S3 and Fig. S1-S10).

## Notes

### Competing Interest Statement

The authors have declared no competing interest.

