## Supplemental Table S1-S3 and Fig. S1-S10 for "Identification and functional characterization of CXCL17 in cartilaginous fishes reveals an ancient origin of the CXCL17–GPR25 signaling pathway"

**Contents:**

**Table S1.** Information for generation of the expression constructs for St-GPR25.

**Table S2.** Information about the cartilaginous fish CXCL17s and their corresponding GPR25s.

**Table S3.** Information about other CXCL17s and GPR25s used in this study.

**Fig. S1.** The nucleotide sequence and amino acid sequence of the St-CXCL17 constructs overexpressed in *E. coli*.

**Fig. S2.** The nucleotide sequence and amino acid sequence of the St-GPR25 constructs overexpressed HEK293T cells.

**Fig. S3.** The nucleotide sequence and amino acid sequence of NanoBiT-based calcium sensor.

**Fig. S4.** Gene position and architecture (**A**), genomic DNA sequence (**B**), cDNA sequence (**C**), and encoded protein sequence (**D**) of the unannotated *cxcl17* gene in the NCBI reference genome of *Scyliorhinus torazame* (cloudy catshark).

**Fig. S5.** Gene position and architecture (**A**), genomic DNA sequence (**B**), cDNA sequence (**C**), and encoded protein sequence (**D**) of the unannotated *cxcl17* gene in the NCBI reference genome of *Scyliorhinus canicular* (smaller spotted catshark).

**Fig. S6.** Gene position and architecture (**A**), genomic DNA sequence (**B**), cDNA sequence (**C**), and encoded protein sequence (**D**) of the unannotated *cxcl17* gene in the NCBI reference genome of *Stegostoma tigrinum*.

**Fig. S7.** Gene position and architecture (**A**), genomic DNA sequence (**B**), cDNA sequence (**C**), and encoded protein sequence (**D**) of the unannotated *cxcl17* gene in the NCBI reference genome of *Chiloscyllium punctatum* (brownbanded bambooshark).

**Fig. S8.** Gene position and architecture (**A**), genomic DNA sequence (**B**), cDNA sequence (**C**), and encoded protein sequence (**D**) of the unannotated *cxcl17* gene in the NCBI reference genome of *Hemiscyllium ocellatum* (epaulette shark).

**Fig. S9.** Gene position and architecture (**A**), genomic DNA sequence (**B**), cDNA sequence (**C**), and encoded protein sequence (**D**) of the unannotated *cxcl17* gene in the NCBI reference genome of *Hemitrygon akajei* (red stingray).

**Fig. S10.** Gene position and architecture (**A**), genomic DNA sequence (**B**), cDNA sequence (**C**), and encoded protein sequence (**D**) of the unannotated *cxcl17* gene in the NCBI reference genome of *Mobula hypostoma* (lesser devil ray).

**Table S1.** Information for generation of the expression constructs for St-GPR25.

| Expression constructs | Vectors for cloning | Restriction enzymes cleaving the vector | Primers for PCR amplification (5' to 3') | Template for PCR amplification | Approach for construct generation |
|---|---|---|---|---|---|
| pcDNA3.1/St-GPR25 | pcDNA3.1(+) | NheI, NotI | Chemical synthesis of the coding region of St-GPR25 | | Gibson assembly |
| pTRE3G-BI/St-GPR25-LgBiT:SmBiT-ARRB2 | pTRE3G-BI/GPR182-LgBiT:SmBiT-ARRB2 | NheI; AgeI (removing GPR182) | Forward: CCG TCA GAT CGC CTG GAG AAT TCG GGG AGA CCC AAG CTG GCT AGC<br>Reverse: GCT AGA CCC TCC GCC GGT ACC GAC CGG TGG GAT TGT TGG GCT CTG CAC CT | pcDNA3.1/St-GPR25 | Gibson assembly |
| PB-TRE/sLgBiT-St-GPR25 | PB-TRE/sLgBiT-GPR182 | KpnI; PmeI (removing GPR182) | Forward: GGT GGC AGC GGC GGT GGT ACC CCC ACC GAA TTT GGA GTC A<br>Reverse: GAG GCT GAT CAG CGG GT TTA GAT TGT TGG GCT CTG CAC CT | pcDNA3.1/St-GPR25 | Gibson assembly |
| PB-TRE/St-GPR25 | PB-TRE/dCas9-VPR | NheI; PmeI (removing dCas9-VPR) | Forward: TTC CTA CCC TCG TAA AGG TCT AGA G CTC ACT ATA GGG AGA CCC AAG CT<br>Reverse: GTT TCA GTT AGC CTC CCC CGT TT C ACA GTC GAG GCT GAT CAG CGG | pcDNA3.1/St-GPR25 | Gibson assembly |

PB-TRE/sLgBiT-GPR182 was generated in our laboratory based on PB-TRE/dCas9-VPR (Addgene cat#: 63800) via removal of the dCas9-VPR fragment by NheI and PmeI cleavage and then ligation of sLgBiT-GPR182 fragment, unpublished data.

pTRE3G-BI/GPR182-LgBiT:SmBiT-ARRB2 was generated in our laboratory, unpublished data.

For oligo primers, the sequence pairing with vector in Gibson assembly is highlighted in yellow, the sequence pairing with PCR template is underlined.

**Table S2.** Information about the cartilaginous fish CXCL17s and their corresponding GPR25s. The cartilaginous fish CXCL17s were identified according to RNA sequencing data in the NCBI reference genome. Their signal peptide was predicted by the SignalP6.0 algorithm and shaded.

| Species | Protein | Gene ID | mRNA ID | Protein ID | Amino acid sequence |
|---|---|---|---|---|---|
| *Scyliorhinus torazame* (cloudy catshark) | CXCL17 | UniProt ID of A0A401NNW6 and NCBI protein ID of GCB62570 derived from early draft genome (Storazame_v1.0);<br><br>Unannotated in latest reference genome (Fig. S4)<br>Reference genome: sScyTor2.1;<br>Chromosome: 12;<br>Position: 99,176,700–99,214,700 bp | | | MQLVSLIAVFALATISVLTDDTTQLKEDEAKRVLTTTRVKSPASEESRCVCKVGHLEFGYKVKQSHDRTGNKCPCQKAINIIKRKWKDVVLSRKGRTKRYCLKPGKTLWRGQRRFYCVESWHRVKKCKSCNPISFIPTPL |
| | GPR25 | 140393839 | XM_072480354 | XP_072336455 | MPTEFGVTEEQDYSLLVYSPDPESDYVDDTDGAICPYENLPFVNITISLLYYLIFIVGSTGNMFVILVMMFKERRRRRLVDTFVINLAFADLVFVFTLPFWAVSASNNHLWAFGSAFCKISSYIVAVNRYSSVFFLTCMSIDRYLAIVKLLNFKHIRTQKYAATISLGIWLSSLLLAIPSAYFRKPDQSNMTYYCTEDTASPFLRAFNVTAINLTFVLPVATILFCYCSILAKLGEHYGHSNKSSQRRENSVKIVFAIVSAFILSWLPFNVLKSMALYLQFHNVDLSCWPLVSRGLAVASCLAFLNSCVNPIIYAFLDRNFRQRTRRMTSHVFAGLGKRSSSFGSGSTVTESSTALKVQSPTI |
| *Scyliorhinus canicular* (smaller spotted catshark) | CXCL17 | Unannotated (Fig. S5)<br>Reference genome: sScyCan1.1;<br>Chromosome: 12;<br>Position: 71,349,500–71,369,700 bp | | | MQLVSLIVVFALVTVSVLTDDTTQLKEGEALRELTTTRVKSPASESHCVCKVGHLEFRHKVKKSHDRTGNKCPCQKAKKIIKRKWKDAVLSRKGGTKRYCLKPGKTLWRGQRRFYCVESWHRMKKCKSCNPIDFIPTPL |
| | GPR25 | 119978173 | XM_038819606 | XP_038675534 | MATEFGVTEQQYYPFLDYSPDSEYDYVDDTDGGICSYENLPFVNIAISLLYYLIFIVGSTGNMFVILVMMFKERKRRRLVDTFVINLAFADLVFVFTLPFWAVSASNNHLWAFGSAFCKISSYIVAVNRYSSVFFLTCMSIDRYLAIVKLLHFKHIRTQKYAATISLGIWLSSLLLAIPSAYFRKPDQSNMTYYCTEDTASPFLRAFNVTAINLTFVLPVAIILFCYCSILVKLGQHYGHSSKSSQRRENSVKIVFAIVSAFILSWLPFNVFKSVALYLQFHNVELSCWPMVSRGLAVASCLAFLNSCVNPIIYAFLDRNFRQRTRRMTSHVFAGLGKRGSSFGSGSTVTESSTALKVQSPTI |
| *Stegostoma tigrinum* | CXCL17 | Unannotated (Fig. S6)<br>Reference genome: sSteTig4.hap1;<br>Chromosome: 41;<br>Position: 10,443,000–10,459,100 bp, complementary strand | | | MQITLLLIAFAFASIPVLTDDTLEPKVVEVSVEGNAGGGRALLTDGNYCECKVGHTQLRYKPQQPLHRAGKKCLCQQPSRTRQRNWKNAFLPKLRKQKRYCLKPGKTISKGKRHFYCVETWHRMKKCTPCKPMDFIPTPL |
| | GPR25 | 125462684 | XM_048552921 | XP_048408878 | MSATVGVTEQHGYTVSEYAPYSEYNDYVDDTDGRICPPQNLHFLNIAISILYYLIFIVGSLGNIFVILVMICKERRRKRLVDTFVINLAFADLIFVFTLPFWAASAISDHHWNFGSAFCKISSYIVAVNRYSSIFFLTCMSIDRYLAIVKLRDFKHLRTQKYAVTISVMIWVSSLLLAVPSAYFRKPDQVNVTQCREDTDSLFLRVFNLTAVSLTFVLPVIIILFCYCSILVKLRHHYGHSTKAIQRRENSLKIVFAIVSAFVLSWLPFNVFKTIALYLQFRDTDLSCWTVVSRGLAIASCIAFINSCVNPIIYAFLDRNFRQRTGWLTSYVFAGLGKRSGSFGSGSTPSESSTMLRIQSLLSQQ |
| *Chiloscyllium punctatum* (brownbanded bambooshark) | CXCL17 | Unannotated (Fig. S7)<br>Reference genome: sChiPun1.3;<br>Chromosome: 34;<br>Position: 38,085,100–38,096,800 bp | | | MQLALLLLIVCVSSYNPVLMEDIRDGKEDKPRVELSANTTGTLEPGRKHCLCKAGHLPLHYRAKQPFVTAGRKCFCQKRSQITRKKHKDALFARQRQGKRYCLKPGKAVYQGQRQLYCVESWASWRKCAPCDPVALIPTPL |
| | GPR25 | 140467290 | XM_072563551 | XP_072419652 | MSTEAGVTEQDGHTFSEDPFYLVYEDYVDDTDGRICPSQNLHFLNIAISMLYYLIFIVGSLGNIFVILVMICKERRRKRLVDTFVINLAFADLIFVFTLPFWAASASNNHQWNFGSVFCKISSYIVAVNRYSSIFFLTCMSIDRYLAIVKLRDFKHIRTQKYAVTISVMIWLSSLLLAIPSAYFRKPDQLNVTQCREDTDSPFLQVFNLTAVSLTFVLPVIIILFCYCSILARLRNHYGHSMKAIQRRENSLKIVFAIVSAFVLSWLPFNVFKTIALYLQFRSANLSCWTVVSRGLAIASCIAFVNSCVNPIIYAFLDRNFRRRTGWLTSYVFAGLGKRNSSFGSGSTPSESSTVVRIQSLLSYQ |
| *Hemiscyllium ocellatum* (epaulette shark) | CXCL17 | Unannotated (Fig. S8)<br>Reference genome: sHemOce1.pat.X.cur;<br>Chromosome: 38;<br>Position: 21,046,400–21,059,800 bp | | | MHLVFLLLIVCVSSYNPVLMQDIRGGKEDKPSVALGTNRTGTRQHGGKYCQCKAGLLPVHSRPKQPIVTAGRKCFCQKPSQIPRRKHRDALFARQRQGKRYCLKPGKAVYRGQRQFYCVESRASWRKCVPCNPIALIPTPL |
| | GPR25 | 132828234 | XM_060845195 | XP_060701178 | MSTEAGVTEQDGHTFSEDPFYLGYEDYVDDTDGRICPSQNLHFLNIAISILYYLIFIVGSLGNIFVILVMICKERRRKRLVDTFVINLAFADLIFVFTLPFWAASASSNHQWNFGSVFCKISSYIVAVNRYSSIFFLTCMSIDRYLAIVKLRDFKHIRTQKYAVTISVMIWLSSLLLAIPSAYFRKPDQLNVTQCREDTDSPFLQVFNLTAVSLTFVLPVIIILFCYCSILAKLRNHYGHSMKAIQRRENSLKIVFAIVSAFVLSWLPFNVFKTIALSLQFRNVNLSCWTGVSRGLAIASCIAFVNSCVNPIIYAFLDRNFRQRTGWLTSYVFAGLGKRNSSFGSGSTPSESSTVVRIQSLLSYQ |
| *Hemitrygon akajei* (red stingray) | CXCL17 | Unannotated (Fig. S9)<br>Reference genome: sHemAka1.3;<br>Chromosome: 1;<br>Position: 184,415,600–184,427,300 bp | | | MITAPPKMQFSTWLFVFALVIISAFPGDATELEEKKKETNKKESRNDENNCACKDTHSDPGLSMQRLQDGIEKKCNCQEVKGVSKKKSSAAAAGKKRQKPHYLQRARHCLIPGRAQRGGIRRYYCIQTKLYLKKCKPCKPDKFIPTPL |
| | GPR25 | 140717019 | XM_073030211 | XP_072886312 | MKMSSEEYFNMTDNYPTLELMTNYYVDDINGDICYHENLPWANISISVLYYLIFLIGSLGNIFVIINMAFKEKKRRRLVDTFVINLAVADLIFVFSLPLWASSAGNNHHWTFGNELCKISSYIIAVNKYSSIFFLTCMSIDRYLAIVKMLDFKHLRTQNYAMTISFVIWFASMLLAIPSAYFRKLIPAQDDEIRCTEDSDSYFYRGFYLFAICLTFILPVIVILFCYCSILNRLRVHYEYNNKVLQRRENSLKIIFAIVSGFVLSWLPFNIFKTISLCLEFSNVNMSCWAIVNHCLAIASCLAFVNSCMNPIIYAFLDRNFRLRAHRFLGCIFGNFRRRRNSIGSVSMATDSSMFAEPSKLNHL |
| *Mobula hypostoma* | CXCL17 | Unannotated (Fig. S10)<br>Reference genome: sMobHyp1.1;<br>Chromosome: 8;<br>Position: 124,882,300–124,895,000 bp | | | MITTPLKMQFFTWLFVFALIIISAFPGDATEQEENKKETNRDESRNDENSCVCKDTHSRLGLKMQQLQDGIGKKCNCQKAKGVSKKLSAAAAGKKRKKPHYLQSPRYCLIPGKAKRGGARRFYCIQTRLYLRKCKPCKPVISIPTPL |

| | | | | | |
|---|---|---|---|---|---|
| （lesser devil ray） | GPR25 | 134355934 | XM_063066470 | XP_062922540 | MSSEEEYFNVTDNYPASEFMTDYFVDDTNGDICYYENLPWANISVSVLYYLIFLIGSLGNILVIINMVFKEKRRRRLVDTFVINLAIADLIFVFSLPLWASTAGNNHRWVFGNKLCKISSYIIAVNRYSSIFFLTCMSIDRYLAIVKLLDFKHIRTQKYAVTISFVVWFVSMLLAIPSAYFKKLVQHDVIQCRDDPDSYFYRGFNLFAICLTFVLPVIVISFCYCSILNRLRVHYEYNRKLLHRRENSVKIIFAIVSGFVLSWLPFNIFRTIILCLEFNNVNLSCWSTVNHSFAIATCLAFVNSCMNPIIYVFLDRNFRLRAHRFLCCTFGNFRRRRNSVGSASMATESSLFAELSKSNL |

**Table S3.** Information about other CXCL17s and GPR25s used in this study. The information was downloaded from the NCBI gene database. The predicted signal peptide of CXCL17s is shaded.

| Class | Name in this study | Species | Gene ID | mRNA ID | Protein ID | Amino acid sequence |
|---|---|---|---|---|---|---|
| CXCL17 orthologs | Hs-CXCL17 | *Homo sapiens* (human) | 284340 | NM_198477 | NP_940879 | MKVLISSLLLLLPLMLMSMVSSSLNPGVARGHRDRGQASRRWLQEGGGQECECKDWFLRAPRRKFMTVSGLPKKQCPCDHFKGNVKKTRHQRHHRKPNKHSRACQQFLKQCQLRSFALPL |
| | Dr-CXCL17 | *Danio rerio* (zebrafish) | 100151367 | NM_001144821 | NP_001138293 | MKTMNFQILVLAFAVMIVTNIQCEARPQEGKSDKSAEVKGHAMPRKCNCQVRGTALDRNCVCEMPHKSRPTLNPEQKNMCLKKKIKTFRKCLQFMGANKKIAKGASLPI |
| | Dr-CXCL17L | *Danio rerio* (zebrafish) | 100536854 | NM_001386806<br>XM_073906074 | NP_001373735<br>XP_073762175 | MTKPICLVFALLILTIILCNNSVCSQRRSMKQSAVCGCKLYPDKGLKCTKRPNPKSRDEYYEILKCICRDTQIFSKSSRKEYLKRCNKFYPSLPL |
| | Lc-CXCL17a | *Latimeria chalumnae* (coelacanth) | 106704074 | XM_014490303 | XP_014345789 | MKVSDVLVLLCAFTLSSCLQNSGSSDQEEGRVPEEASTVGAVKPQAGSCSCGDPVGSLQNRLVSLKAGPSKHCDCRRSLKEAVNIKRPQAWAWKSKPSRKRCKRKGDNGKIQIWGCRRKTIKPIIIPIPA |
| | Lc-CXCL17b | *Latimeria chalumnae* (coelacanth) | 106704074 | XM_014490304 | XP_014345790 | MKVSDVLVLLCAFTLSSCLQNSGSSDQEEGRVPEEASTVGAVKPQAGSCSCGDPVGSLQNRLVSLKAGPSKHCDCRRSLKEVNIKRPQAWAWKSKPSRKRCKRKGDNGKIQIWGCRRKTIKPIIIPIPA |
| GPR25 orthologs | Hs-GPR25 | *Homo sapiens* (human) | 2848 | NM_005298 | NP_005289 | MAPTEPWSPSPGSAPWDYSGLDGLEELELCPAGDLPYGYVYIPALYLAAFAVGLLGNAFVVWLLAGRRGPRRLVDTFVLHLAAADLGFVLTLPLWAAAAALGGRWPFGDGLCKLSSFALAGTRCAGALLLAGMSVDRYLAVVKLLEARPLRTPRCALASCCGVWAVALLAGLPSLVYRGLQPLPGGQDSQCGEEPSHAFQGLSLLLLLLTFVLPLVVTLFCYCRISRRLRRPPHVGRARRNSLRIIFAIESTFVGSWLPFSALRAVFHLARLGALPLPCPLLLALRWGLTIATCLAFVNSCANPLIYLLLDRSFRARALDGACGRTGRLARRISSASSLSRDDSSVFRCRAQAANTASASW |
| | Dr-GPR25 | *Danio rerio* (zebrafish) | 795188 | XM_073916757 | XP_073772858 | MASSTEMAHSGITMSLTSEYDYDYPINSTDENPIYTLPDAELLPMSNIYIPVLYIIMFLTGSLGNLFVIVVIGKRRKKSGRLVDTFVLNLALADLVFVLTLPMWAISTRYDEWPFGEVLCKISSFIIAVNRFSNIFFLTCMSVDRYLAVVRLMDSRFLRSSNCAQITCGIVWVVSFFLGSPSLAYRHLINNSVCSEDSKSSFVQGMNLLTILLTFLLPVLILGLCYGSILVNLRRHCHNPANTRTDARRRHSVKIVFAIISAFLISWLPFNCFKAIHVALLIINGDLNEDTYVVIHRGLMLSCCLAFLNSCVNPAIYFFLDQHFRRRASMLCLSCLSQNDQAHQSYITSNSYSNGTSETCSGNTSTRGRLFSLTQKA |
| | Lc-GPR25 | *Latimeria chalumnae* (coelacanth) | 102365624 | XM_005988473 | XP_005988535 | MPTESLQTASHDPSDFDYNADYSNFSTEDCDGDLPYAKIYIPIFYFVIFFTGLFGNVFVIAAMTLKQTTKRLVDIFVINLAVADLVFVFTLPLWSVSAAFDDQWLFGGVLCKLSSYVIAVNRYSSIFFMTGMSVDRYMAVVKLLDSKFIRTRRCILITCTIIWIISLVMGIPSLVYRDLSTQDSEHTYCIEDQDSIIFKGISLASLFLAFVLPVIIILFCYCSISARLYSHFHHANRYDQKRKKTLKIIFTIITAFVCSWLPFNTFKTLYLLFSFQGKMPPCRVGILRQGLTITACFAFLSSCVNPIIYTFLDNHFRKRAHRLLVKALGRYTERRNSFGESWASETSSTFVSRIRANSVKELQNMNKTQQNTIPT |

**6×His-St-CXCL17**

```
      NdeI
1     CAT ATG CAT CAC CAC CAC CAC CAT GAT GAC ACA ACG CAG CTG AAA GAG GAC GAA GCT AAA CGC GTG TTG ACC ACC ACT CGC GTC AAG TCG
      GTA TAC GTA GTG GTG GTG GTG GTA CTA CTG TGT TGC GTC GAC TTT CTC CTG CTT CGA TTT GCG CAC AAC TGG TGG TGA GCG CAG TTC AGC
           M   H   H   H   H   H   D   D   T   T   Q   L   K   E   D   E   A   K   R   V   L   T   T   T   R   V   K   S

91    CCG GCA AGC GAG GAA TCC CGT TGT GTT TGC AAA GTT GGT CAC CTT GAG TTC GGC TAT AAA GTG AAG CAA AGC CAT GAT CGT ACC GGC AAC
      GGC CGT TCG CTC CTT AGG GCA ACA CAA ACG TTT CAA CCA GTG GAA CTC AAG CCG ATA TTT CAC TTC GTT TCG GTA CTA GCA TGG CCG TTG
       P   A   S   E   E   S   R   C   V   C   K   V   G   H   L   E   F   G   Y   K   V   K   Q   S   H   D   R   T   G   N

181   AAA TGC CCG TGT CAG AAA GCG ATT AAC ATC ATC AAA CGC AAG TGG AAG GAC GTT GTT CTG AGC CGT AAG GGT CGT ACG AAA CGC TAC TGC
      TTT ACG GGC ACA GTC TTT CGC TAA TTG TAG TAG TTT GCG TTC ACC TTC CTG CAA CAA GAC TCG GCA TTC CCA GCA TGC TTT GCG ATG ACG
       K   C   P   C   Q   K   A   I   N   I   I   K   R   K   W   K   D   V   V   L   S   R   K   G   R   T   K   R   Y   C

271   CTG AAG CCG GGT AAA ACC CTG TGG AGA GGC CAA CGT CGT TTC TAC TGC GTG GAA TCC TGG CAC CGT GTG AAG AAG TGC AAG AGC TGT AAT
      GAC TTC GGC CCA TTT TGG GAC ACC TCT CCG GTT GCA GCA AAG ATG ACG CAC CTT AGG ACC GTG GCA CAC TTC TTC ACG TTC TCG ACA TTA
       L   K   P   G   K   T   L   W   R   G   Q   R   R   F   Y   C   V   E   S   W   H   R   V   K   K   C   K   S   C   N

                                              EcoRI
361   CCG ATT TCT TTT ATC CCG ACC CCA CTG TAA GAA TTC
      GGC TAA AGA AAA TAG GGC TGG GGT GAC ATT CTT AAG
       P   I   S   F   I   P   T   P   L   *
```

**6×His-SmBiT-St-CXCL17**

```
      NsiI
1     ATG CAT CAT CAC CAT CAC CAT GGT GTG ACC GGC TAC CGT CTG TTT GAA GAA ATT CTG GGC GGC GAT GAC ACA ACG CAG CTG AAA GAG GAC
      TAC GTA GTA GTG GTA GTG GTA CCA CAC TGG CCG ATG GCA GAC AAA CTT CTT TAA GAC CCG CCG CTA CTG TGT TGC GTC GAC TTT CTC CTG
       M   H   H   H   H   H   H   G   V   T   G   Y   R   L   F   E   E   I   L   G   G   D   D   T   T   Q   L   K   E   D

91    GAA GCT AAA CGC GTG TTG ACC ACC ACT CGC GTC AAG TCG CCG GCA AGC GAG GAA TCC CGT TGT GTT TGC AAA GTT GGT CAC CTT GAG TTC
      CTT CGA TTT GCG CAC AAC TGG TGG TGA GCG CAG TTC AGC GGC CGT TCG CTC CTT AGG GCA ACA CAA ACG TTT CAA CCA GTG GAA CTC AAG
       E   A   K   R   V   L   T   T   T   R   V   K   S   P   A   S   E   E   S   R   C   V   C   K   V   G   H   L   E   F

181   GGC TAT AAA GTG AAG CAA AGC CAT GAT CGT ACC GGC AAC AAA TGC CCG TGT CAG AAA GCG ATT AAC ATC ATC AAA CGC AAG TGG AAG GAC
      CCG ATA TTT CAC TTC GTT TCG GTA CTA GCA TGG CCG TTG TTT ACG GGC ACA GTC TTT CGC TAA TTG TAG TAG TTT GCG TTC ACC TTC CTG
       G   Y   K   V   K   Q   S   H   D   R   T   G   N   K   C   P   C   Q   K   A   I   N   I   I   K   R   K   W   K   D

271   GTT GTT CTG AGC CGT AAG GGT CGT ACG AAA CGC TAC TGC CTG AAG CCG GGT AAA ACC CTG TGG AGA GGC CAA CGT CGT TTC TAC TGC GTG
      CAA CAA GAC TCG GCA TTC CCA GCA TGC TTT GCG ATG ACG GAC TTC GGC CCA TTT TGG GAC ACC TCT CCG GTT GCA GCA AAG ATG ACG CAC
       V   V   L   S   R   K   G   R   T   K   R   Y   C   L   K   P   G   K   T   L   W   R   G   Q   R   R   F   Y   C   V

                                                                                                      EcoRI
361   GAA TCC TGG CAC CGT GTG AAG AAG TGC AAG AGC TGT AAT CCG ATT TCT TTT ATC CCG ACC CCA CTG TAA GAA TTC
      CTT AGG ACC GTG GCA CAC TTC TTC ACG TTC TCG ACA TTA GGC TAA AGA AAA TAG GGC TGG GGT GAC ATT CTT AAG
       E   S   W   H   R   V   K   K   C   K   S   C   N   P   I   S   F   I   P   T   P   L   *
```

**Fig. S1.** The nucleotide sequence and amino acid sequence of the St-CXCL17 constructs overexpressed in *E. coli*. The amino acid sequence of the mature St-CXCL17 is shown in red, that of SmBiT tag in blue.

#### Untagged St-GPR25 in pcDNA3.1 or PB-TRE

```
1     ATG CCC ACC GAA TTT GGA GTC ACC GAG GAG CAG GAC TAT TCA CTT CTT GTT TAT TCT CCA GAT CCT GAG TCT GAC TAT GTG GAT GAC ACA GAT GGA GCA ATC TGT
      TAC GGG TGG CTT AAA CCT CAG TGG CTC CTC GTC CTG ATA AGT GAA GAA CAA ATA AGA GGT CTA GGA CTC AGA CTG ATA CAC CTA CTG TGT CTA CCT CGT TAG ACA
       M   P   T   E   F   G   V   T   E   E   Q   D   Y   S   L   L   V   Y   S   P   D   P   E   S   D   Y   V   D   D   T   D   G   A   I   C

106   CCT TAT GAG AAT CTA CCT TTT GTA AAC ATC ACT ATC TCA CTT CTC TAC TAC CTG ATC TTC ATC GTT GGG TCT ACG GGG AAC ATG TTT GTC ATC TTG GTG ATG ATG
      GGA ATA CTC TTA GAT GGA AAA CAT TTG TAG TGA TAG AGT GAA GAG ATG ATG GAC TAG AAG TAG CAA CCC AGA TGC CCC TTG TAC AAA CAG TAG AAC CAC TAC TAC
       P   Y   E   N   L   P   F   V   N   I   T   I   S   L   L   Y   Y   L   I   F   I   V   G   S   T   G   N   M   F   V   I   L   V   M   M

211   TTC AAA GAG AGG AGG AGG AGG AGA CTA GTG GAC ACC TTT GTG ATC AAC CTA GCG TTT GCA GAC CTA GTC TTT GTC TTC ACC CTG CCT TTC TGG GCG GTG TCG GCC
      AAG TTT CTC TCC TCC TCC TCC TCT GAT CAC CTG TGG AAA CAC TAG TTG GAT CGC AAA CGT CTG GAT CAG AAA CAG AAG TGG GAC GGA AAG ACC CGC CAC AGC CGG
       F   K   E   R   R   R   R   R   L   V   D   T   F   V   I   N   L   A   F   A   D   L   V   F   V   F   T   L   P   F   W   A   V   S   A

316   AGC AAC AAC CAT CTG TGG GCC TTC GGC AGT GCC TTC TGC AAA ATC AGC AGC TAC ATC GTT GCT GTC AAC AGA TAC TCC AGC GTC TTC TTC CTG ACC TGC ATG AGC
      TCG TTG TTG GTA GAC ACC CGG AAG CCG TCA CGG AAG ACG TTT TAG TCG TCG ATG TAG CAA CGA CAG TTG TCT ATG AGG TCG CAG AAG AAG GAC TGG ACG TAC TCG
       S   N   N   H   L   W   A   F   G   S   A   F   C   K   I   S   S   Y   I   V   A   V   N   R   Y   S   S   V   F   F   L   T   C   M   S

421   ATT GAC CGG TAT CTG GCC ATC GTC AAG CTG CTG AAC TTC AAG CAC ATC CGG ACC CAG AAG TAC GCC GCC ACG ATC AGC TTG GGG ATC TGG CTG TCA TCC CTG CTG
      TAA CTG GCC ATA GAC CGG TAG CAG TTC GAC GAC TTG AAG TTC GTG TAG GCC TGG GTC TTC ATG CGG CGG TGC TAG TCG AAC CCC TAG ACC GAC AGT AGG GAC GAC
       I   D   R   Y   L   A   I   V   K   L   L   N   F   K   H   I   R   T   Q   K   Y   A   A   T   I   S   L   G   I   W   L   S   S   L   L

526   CTG GCC ATC CCC TCG GCC TAT TTC CGG AAA CCT GAC CAG TCC AAC ATG ACC TAC TAC TGC ACA GAG GAC ACG GCA TCT CCC TTC CTG CGG GCT TTC AAC GTG ACT
      GAC CGG TAG GGG AGC CGG ATA AAG GCC TTT GGA CTG GTC AGG TTG TAC TGG ATG ATG ACG TGT CTC CTG TGC CGT AGA GGG AAG GAC GCC CGA AAG TTG CAC TGA
       L   A   I   P   S   A   Y   F   R   K   P   D   Q   S   N   M   T   Y   Y   C   T   E   D   T   A   S   P   F   L   R   A   F   N   V   T

631   GCC ATC AAC CTG ACA TTT GTC CTG CCT GTT GCC ACC ATC CTG TTC TGC TAC TGC TCC ATC CTG GCC AAG CTC GGG GAG CAT TAC GGC CAC AGC AAC AAA AGC AGC
      CGG TAG TTG GAC TGT AAA CAG GAC GGA CAA CGG TGG TAG GAC AAG ACG ATG ACG AGG TAG GAC CGG TTC GAG CCC CTC GTA ATG CCG GTG TCG TTG TTT TCG TCG
       A   I   N   L   T   F   V   L   P   V   A   T   I   L   F   C   Y   C   S   I   L   A   K   L   G   E   H   Y   G   H   S   N   K   S   S

736   CAG AGG AGG GAA AAC TCC GTA AAA ATC GTC TTC GCC ATC GTG TCG GCT TTT ATT TTA TCC TGG CTG CCC TTT AAC GTC TTA AAA TCG ATG GCC CTG TAC CTG CAG
      GTC TCC TCC CTT TTG AGG CAT TTT TAG CAG AAG CGG TAG CAC AGC CGA AAA TAA AAT AGG ACC GAC GGG AAA TTG CAG AAT TTT AGC TAC CGG GAC ATG GAC GTC
       Q   R   R   E   N   S   V   K   I   V   F   A   I   V   S   A   F   I   L   S   W   L   P   F   N   V   L   K   S   M   A   L   Y   L   Q

841   TTC CAC AAT GTG GAC CTC TCC TGC TGG CCG CTG GTC AGC CGG GGT CTG GCC GTG GCT TCC TGC TTG GCT TTC CTC AAT AGC TGC GTC AAC CCC ATC ATT TAT GCC
      AAG GTG TTA CAC CTG GAG AGG ACG ACC GGC GAC CAG TCG GCC CCA GAC CGG CAC CGA AGG ACG AAC CGA AAG GAG TTA TCG ACG CAG TTG GGG TAG TAA ATA CGG
       F   H   N   V   D   L   S   C   W   P   L   V   S   R   G   L   A   V   A   S   C   L   A   F   L   N   S   C   V   N   P   I   I   Y   A

946   TTC CTG GAC CGC AAT TTC AGG CAG AGG ACC CGC CGG ATG ACC TCG CAC GTC TTC GCC GGC CTG GGG AAG AGG AGC AGC AGC TTC GGC TCC GGC TCC ACG GTC ACA
      AAG GAC CTG GCG TTA AAG TCC GTC TCC TGG GCG GCC TAC TGG AGC GTG CAG AAG CGG CCG GAC CCC TTC TCC TCG TCG TCG AAG CCG AGG CCG AGG TGC CAG TGT
       F   L   D   R   N   F   R   Q   R   T   R   R   M   T   S   H   V   F   A   G   L   G   K   R   S   S   S   F   G   S   G   S   T   V   T

1051  GAG AGC AGC ACC GCG CTG AAG GTG CAG AGC CCA ACA ATC TGA
      CTC TCG TCG TGG CGC GAC TTC CAC GTC TCG GGT TGT TAG ACT
       E   S   S   T   A   L   K   V   Q   S   P   T   I   *
```

#### St-GPR25-LgBiT in MCS1 of pTRE3G-BI vector (SmBiT-ARRB2 in MCS2)

```
1     ATG CCC ACC GAA TTT GGA GTC ACC GAG GAG CAG GAC TAT TCA CTT CTT GTT TAT TCT CCA GAT CCT GAG TCT GAC TAT GTG GAT GAC ACA GAT GGA GCA ATC TGT
      TAC GGG TGG CTT AAA CCT CAG TGG CTC CTC GTC CTG ATA AGT GAA GAA CAA ATA AGA GGT CTA GGA CTC AGA CTG ATA CAC CTA CTG TGT CTA CCT CGT TAG ACA
       M   P   T   E   F   G   V   T   E   E   Q   D   Y   S   L   L   V   Y   S   P   D   P   E   S   D   Y   V   D   D   T   D   G   A   I   C

106   CCT TAT GAG AAT CTA CCT TTT GTA AAC ATC ACT ATC TCA CTT CTC TAC TAC CTG ATC TTC ATC GTT GGG TCT ACG GGG AAC ATG TTT GTC ATC TTG GTG ATG ATG
      GGA ATA CTC TTA GAT GGA AAA CAT TTG TAG TGA TAG AGT GAA GAG ATG ATG GAC TAG AAG TAG CAA CCC AGA TGC CCC TTG TAC AAA CAG TAG AAC CAC TAC TAC
       P   Y   E   N   L   P   F   V   N   I   T   I   S   L   L   Y   Y   L   I   F   I   V   G   S   T   G   N   M   F   V   I   L   V   M   M

211   TTC AAA GAG AGG AGG AGG AGG AGA CTA GTG GAC ACC TTT GTG ATC AAC CTA GCG TTT GCA GAC CTA GTC TTT GTC TTC ACC CTG CCT TTC TGG GCG GTG TCG GCC
      AAG TTT CTC TCC TCC TCC TCC TCT GAT CAC CTG TGG AAA CAC TAG TTG GAT CGC AAA CGT CTG GAT CAG AAA CAG AAG TGG GAC GGA AAG ACC CGC CAC AGC CGG
       F   K   E   R   R   R   R   R   L   V   D   T   F   V   I   N   L   A   F   A   D   L   V   F   V   F   T   L   P   F   W   A   V   S   A

316   AGC AAC AAC CAT CTG TGG GCC TTC GGC AGT GCC TTC TGC AAA ATC AGC AGC TAC ATC GTT GCT GTC AAC AGA TAC TCC AGC GTC TTC TTC CTG ACC TGC ATG AGC
      TCG TTG TTG GTA GAC ACC CGG AAG CCG TCA CGG AAG ACG TTT TAG TCG TCG ATG TAG CAA CGA CAG TTG TCT ATG AGG TCG CAG AAG AAG GAC TGG ACG TAC TCG
       S   N   N   H   L   W   A   F   G   S   A   F   C   K   I   S   S   Y   I   V   A   V   N   R   Y   S   S   V   F   F   L   T   C   M   S

421   ATT GAC CGG TAT CTG GCC ATC GTC AAG CTG CTG AAC TTC AAG CAC ATC CGG ACC CAG AAG TAC GCC GCC ACG ATC AGC TTG GGG ATC TGG CTG TCA TCC CTG CTG
      TAA CTG GCC ATA GAC CGG TAG CAG TTC GAC GAC TTG AAG TTC GTG TAG GCC TGG GTC TTC ATG CGG CGG TGC TAG TCG AAC CCC TAG ACC GAC AGT AGG GAC GAC
       I   D   R   Y   L   A   I   V   K   L   L   N   F   K   H   I   R   T   Q   K   Y   A   A   T   I   S   L   G   I   W   L   S   S   L   L

526   CTG GCC ATC CCC TCG GCC TAT TTC CGG AAA CCT GAC CAG TCC AAC ATG ACC TAC TAC TGC ACA GAG GAC ACG GCA TCT CCC TTC CTG CGG GCT TTC AAC GTG ACT
      GAC CGG TAG GGG AGC CGG ATA AAG GCC TTT GGA CTG GTC AGG TTG TAC TGG ATG ATG ACG TGT CTC CTG TGC CGT AGA GGG AAG GAC GCC CGA AAG TTG CAC TGA
       L   A   I   P   S   A   Y   F   R   K   P   D   Q   S   N   M   T   Y   Y   C   T   E   D   T   A   S   P   F   L   R   A   F   N   V   T

631   GCC ATC AAC CTG ACA TTT GTC CTG CCT GTT GCC ACC ATC CTG TTC TGC TAC TGC TCC ATC CTG GCC AAG CTC GGG GAG CAT TAC GGC CAC AGC AAC AAA AGC AGC
      CGG TAG TTG GAC TGT AAA CAG GAC GGA CAA CGG TGG TAG GAC AAG ACG ATG ACG AGG TAG GAC CGG TTC GAG CCC CTC GTA ATG CCG GTG TCG TTG TTT TCG TCG
       A   I   N   L   T   F   V   L   P   V   A   T   I   L   F   C   Y   C   S   I   L   A   K   L   G   E   H   Y   G   H   S   N   K   S   S

736   CAG AGG AGG GAA AAC TCC GTA AAA ATC GTC TTC GCC ATC GTG TCG GCT TTT ATT TTA TCC TGG CTG CCC TTT AAC GTC TTA AAA TCG ATG GCC CTG TAC CTG CAG
      GTC TCC TCC CTT TTG AGG CAT TTT TAG CAG AAG CGG TAG CAC AGC CGA AAA TAA AAT AGG ACC GAC GGG AAA TTG CAG AAT TTT AGC TAC CGG GAC ATG GAC GTC
       Q   R   R   E   N   S   V   K   I   V   F   A   I   V   S   A   F   I   L   S   W   L   P   F   N   V   L   K   S   M   A   L   Y   L   Q

841   TTC CAC AAT GTG GAC CTC TCC TGC TGG CCG CTG GTC AGC CGG GGT CTG GCC GTG GCT TCC TGC TTG GCT TTC CTC AAT AGC TGC GTC AAC CCC ATC ATT TAT GCC
      AAG GTG TTA CAC CTG GAG AGG ACG ACC GGC GAC CAG TCG GCC CCA GAC CGG CAC CGA AGG ACG AAC CGA AAG GAG TTA TCG ACG CAG TTG GGG TAG TAA ATA CGG
       F   H   N   V   D   L   S   C   W   P   L   V   S   R   G   L   A   V   A   S   C   L   A   F   L   N   S   C   V   N   P   I   I   Y   A

946   TTC CTG GAC CGC AAT TTC AGG CAG AGG ACC CGC CGG ATG ACC TCG CAC GTC TTC GCC GGC CTG GGG AAG AGG AGC AGC AGC TTC GGC TCC GGC TCC ACG GTC ACA
      AAG GAC CTG GCG TTA AAG TCC GTC TCC TGG GCG GCC TAC TGG AGC GTG CAG AAG CGG CCG GAC CCC TTC TCC TCG TCG TCG AAG CCG AGG CCG AGG TGC CAG TGT
       F   L   D   R   N   F   R   Q   R   T   R   R   M   T   S   H   V   F   A   G   L   G   K   R   S   S   S   F   G   S   G   S   T   V   T

1051  GAG AGC AGC ACC GCG CTG AAG GTG CAG AGC CCA ACA ATC CCA CCG GTC GGT ACC GGC GGA GGG TCT AGC AGT GGC GGT GGG ATG GTC TTC ACA CTC GAA GAT TTC
      CTC TCG TCG TGG CGC GAC TTC CAC GTC TCG GGT TGT TAG GGT GGC CAG CCA TGG CCG CCT CCC AGA TCG TCA CCG CCA CCC TAC CAG AAG TGT GAG CTT CTA AAG
       E   S   S   T   A   L   K   V   Q   S   P   T   I   P   P   V   G   T   G   G   G   S   S   S   G   G   G   M   V   F   T   L   E   D   F

1156  GTT GGG GAC TGG GAA CAG ACA GCC GCC TAC AAC CTG GAC CAA GTC CTT GAA CAG GGA GGT GTG TCC AGT TTG CTG CAG AAT CTC GCC GTG TCC GTA ACT CCG ATC
      CAA CCC CTG ACC CTT GTC TGT CGG CGG ATG TTG GAC CTG GTT CAG GAA CTT GTC CCT CCA CAC AGG TCA AAC GAC GTC TTA GAG CGG CAC AGG CAT TGA GGC TAG
       V   G   D   W   E   Q   T   A   A   Y   N   L   D   Q   V   L   E   Q   G   G   V   S   S   L   L   Q   N   L   A   V   S   V   T   P   I

1261  CAA AGG ATT GTC CGG AGC GGT GAA AAT GCC CTG AAG ATC GAC ATC CAT GTC ATC ATC CCG TAT GAA GGT CTG AGC GCC GAC CAA ATG GCC CAG ATC GAA GAG GTG
      GTT TCC TAA CAG GCC TCG CCA CTT TTA CGG GAC TTC TAG CTG TAG GTA CAG TAG TAG GGC ATA CTT CCA GAC TCG CGG CTG GTT TAC CGG GTC TAG CTT CTC CAC
       Q   R   I   V   R   S   G   E   N   A   L   K   I   D   I   H   V   I   I   P   Y   E   G   L   S   A   D   Q   M   A   Q   I   E   E   V

1366  TTT AAG GTG GTG TAC CCT GTG GAT GAT CAT CAC TTT AAG GTG ATC CTG CCC TAT GGC ACA CTG GTA ATC GAC GGG GTT ACG CCG AAC ATG CTG AAC TAT TTC GGA
```

```
     AAA TTC CAC CAC ATG GGA CAC CTA CTA GTA GTG AAA TTC CAC TAG GAC GGG ATA CCG TGT GAC CAT TAG CTG CCC CAA TGC GGC TTG TAC GAC TTG ATA AAG CCT
      F   K   V   V   Y   P   V   D   D   H   H   F   K   V   I   L   P   Y   G   T   L   V   I   D   G   V   T   P   N   M   L   N   Y   F   G

1471 CGG CCG TAT GAA GGC ATC GCC GTG TTC GAC GGC AAA AAG ATC ACT GTA ACA GGG ACC CTG TGG AAC GGC AAC AAA ATT ATC GAC GAG CGC CTG ATC ACC CCC GAC
     GCC GGC ATA CTT CCG TAG CGG CAC AAG CTG CCG TTT TTC TAG TGA CAT TGT CCC TGG GAC ACC TTG CCG TTG TTT TAA TAG CTG CTC GCG GAC TAG TGG GGG CTG
      R   P   Y   E   G   I   A   V   F   D   G   K   K   I   T   V   T   G   T   L   W   N   G   N   K   I   I   D   E   R   L   I   T   P   D

1576 GGC TCC ATG CTG TTC CGA GTA ACC ATC AAC AGT TAA
     CCG AGG TAC GAC AAG GCT CAT TGG TAG TTG TCA ATT
      G   S   M   L   F   R   V   T   I   N   S   *
```

**sLgBiT-St-GPR25 in PB-TRE vector**

```
1    ATG AAC TCC TTC TCC ACA AGC GCC TTC GGT CCA GTT GCC TTC TCC CTG GGC CTG CTC CTG GTG TTG CCT GCT GCC TTC CCT GCC CCA GTC TTC ACA CTC GAA GAT
     TAC TTG AGG AAG AGG TGT TCG CGG AAG CCA GGT CAA CGG AAG AGG GAC CCG GAC GAG GAC CAC AAC GGA CGA CGG AAG GGA CGG GGT CAG AAG TGT GAG CTT CTA
      M   N   S   F   S   T   S   A   F   G   P   V   A   F   S   L   G   L   L   L   V   L   P   A   A   F   P   A   P   V   F   T   L   E   D

106  TTC GTT GGG GAC TGG GAA CAG ACA GCC GCC TAC AAC CTG GAC CAA GTC CTT GAA CAG GGA GGT GTG TCC AGT TTG CTG CAG AAT CTC GCC GTG TCC GTA ACT CCG
     AAG CAA CCC CTG ACC CTT GTC TGT CGG CGG ATG TTG GAC CTG GTT CAG GAA CTT GTC CCT CCA CAC AGG TCA AAC GAC GTC TTA GAG CGG CAC AGG CAT TGA GGC
      F   V   G   D   W   E   Q   T   A   A   Y   N   L   D   Q   V   L   E   Q   G   G   V   S   S   L   L   Q   N   L   A   V   S   V   T   P

211  ATC CAA AGG ATT GTC CGG AGC GGT GAA AAT GCC CTG AAG ATC GAC ATC CAT GTC ATC ATC CCG TAT GAA GGT CTG AGC GCC GAC CAA ATG GCC CAG ATC GAA GAG
     TAG GTT TCC TAA CAG GCC TCG CCA CTT TTA CGG GAC TTC TAG CTG TAG GTA CAG TAG TAG GGC ATA CTT CCA GAC TCG CGG CTG GTT TAC CGG GTC TAG CTT CTC
      I   Q   R   I   V   R   S   G   E   N   A   L   K   I   D   I   H   V   I   I   P   Y   E   G   L   S   A   D   Q   M   A   Q   I   E   E

316  GTG TTT AAG GTG GTG TAC CCT GTG GAT GAT CAT CAC TTT AAG GTG ATC CTG CCC TAT GGC ACA CTG GTA ATC GAC GGG GTT ACG CCG AAC ATG CTG AAC TAT TTC
     CAC AAA TTC CAC CAC ATG GGA CAC CTA CTA GTA GTG AAA TTC CAC TAG GAC GGG ATA CCG TGT GAC CAT TAG CTG CCC CAA TGC GGC TTG TAC GAC TTG ATA AAG
      V   F   K   V   V   Y   P   V   D   D   H   H   F   K   V   I   L   P   Y   G   T   L   V   I   D   G   V   T   P   N   M   L   N   Y   F

421  GGA CGG CCG TAT GAA GGC ATC GCC GTG TTC GAC GGC AAA AAG ATC ACT GTA ACA GGG ACC CTG TGG AAC GGC AAC AAA ATT ATC GAC GAG CGC CTG ATC ACC CCC
     CCT GCC GGC ATA CTT CCG TAG CGG CAC AAG CTG CCG TTT TTC TAG TGA CAT TGT CCC TGG GAC ACC TTG CCG TTG TTT TAA TAG CTG CTC GCG GAC TAG TGG GGG
      G   R   P   Y   E   G   I   A   V   F   D   G   K   K   I   T   V   T   G   T   L   W   N   G   N   K   I   I   D   E   R   L   I   T   P

526  GAC GGC TCC ATG CTG TTC CGA GTA ACC ATC AAC AGT GGT GGC GGC TCT GGT GGT GGC AGC GGC GGT GGT ACC CCC ACC GAA TTT GGA GTC ACC GAG GAG CAG GAC
     CTG CCG AGG TAC GAC AAG GCT CAT TGG TAG TTG TCA CCA CCG CCG AGA CCA CCA CCG TCG CCG CCA CCA TGG GGG TGG CTT AAA CCT CAG TGG CTC CTC GTC CTG
      D   G   S   M   L   F   R   V   T   I   N   S   G   G   G   S   G   G   G   S   G   G   G   T   P   T   E   F   G   V   T   E   E   Q   D

631  TAT TCA CTT CTT GTT TAT TCT CCA GAT CCT GAG TCT GAC TAT GTG GAT GAC ACA GAT GGA GCA ATC TGT CCT TAT GAG AAT CTA CCT TTT GTA AAC ATC ACT ATC
     ATA AGT GAA GAA CAA ATA AGA GGT CTA GGA CTC AGA CTG ATA CAC CTA CTG TGT CTA CCT CGT TAG ACA GGA ATA CTC TTA GAT GGA AAA CAT TTG TAG TGA TAG
      Y   S   L   L   V   Y   S   P   D   P   E   S   D   Y   V   D   D   T   D   G   A   I   C   P   Y   E   N   L   P   F   V   N   I   T   I

736  TCA CTT CTC TAC TAC CTG ATC TTC ATC GTT GGG TCT ACG GGG AAC ATG TTT GTC ATC TTG GTG ATG ATG TTC AAA GAG AGG AGG AGG AGG AGA CTA GTG GAC ACC
     AGT GAA GAG ATG ATG GAC TAG AAG TAG CAA CCC AGA TGC CCC TTG TAC AAA CAG TAG AAC CAC TAC TAC AAG TTT CTC TCC TCC TCC TCC TCT GAT CAC CTG TGG
      S   L   L   Y   Y   L   I   F   I   V   G   S   T   G   N   M   F   V   I   L   V   M   M   F   K   E   R   R   R   R   R   L   V   D   T

841  TTT GTG ATC AAC CTA GCG TTT GCA GAC CTA GTC TTT GTC TTC ACC CTG CCT TTC TGG GCG GTG TCG GCC AGC AAC AAC CAT CTG TGG GCC TTC GGC AGT GCC TTC
     AAA CAC TAG TTG GAT CGC AAA CGT CTG GAT CAG AAA CAG AAG TGG GAC GGA AAG ACC CGC CAC AGC CGG TCG TTG TTG GTA GAC ACC CGG AAG CCG TCA CGG AAG
      F   V   I   N   L   A   F   A   D   L   V   F   V   F   T   L   P   F   W   A   V   S   A   S   N   N   H   L   W   A   F   G   S   A   F

946  TGC AAA ATC AGC AGC TAC ATC GTT GCT GTC AAC AGA TAC TCC AGC GTC TTC TTC CTG ACC TGC ATG AGC ATT GAC CGG TAT CTG GCC ATC GTC AAG CTG CTG AAC
     ACG TTT TAG TCG TCG ATG TAG CAA CGA CAG TTG TCT ATG AGG TCG CAG AAG AAG GAC TGG ACG TAC TCG TAA CTG GCC ATA GAC CGG TAG CAG TTC GAC GAC TTG
      C   K   I   S   S   Y   I   V   A   V   N   R   Y   S   S   V   F   F   L   T   C   M   S   I   D   R   Y   L   A   I   V   K   L   L   N

1051 TTC AAG CAC ATC CGG ACC CAG AAG TAC GCC GCC ACG ATC AGC TTG GGG ATC TGG CTG TCA TCC CTG CTG CTG GCC ATC CCC TCG GCC TAT TTC CGG AAA CCT GAC
     AAG TTC GTG TAG GCC TGG GTC TTC ATG CGG CGG TGC TAG TCG AAC CCC TAG ACC GAC AGT AGG GAC GAC GAC CGG TAG GGG AGC CGG ATA AAG GCC TTT GGA CTG
      F   K   H   I   R   T   Q   K   Y   A   A   T   I   S   L   G   I   W   L   S   S   L   L   L   A   I   P   S   A   Y   F   R   K   P   D

1156 CAG TCC AAC ATG ACC TAC TAC TGC ACA GAG GAC ACG GCA TCT CCC TTC CTG CGG GCT TTC AAC GTG ACT GCC ATC AAC CTG ACA TTT GTC CTG CCT GTT GCC ACC
     GTC AGG TTG TAC TGG ATG ATG ACG TGT CTC CTG TGC CGT AGA GGG AAG GAC GCC CGA AAG TTG CAC TGA CGG TAG TTG GAC TGT AAA CAG GAC GGA CAA CGG TGG
      Q   S   N   M   T   Y   Y   C   T   E   D   T   A   S   P   F   L   R   A   F   N   V   T   A   I   N   L   T   F   V   L   P   V   A   T

1261 ATC CTG TTC TGC TAC TGC TCC ATC CTG GCC AAG CTC GGG GAG CAT TAC GGC CAC AGC AAC AAA AGC AGC CAG AGG AGG GAA AAC TCC GTA AAA ATC GTC TTC GCC
     TAG GAC AAG ACG ATG ACG AGG TAG GAC CGG TTC GAG CCC CTC GTA ATG CCG GTG TCG TTG TTT TCG TCG GTC TCC TCC CTT TTG AGG CAT TTT TAG CAG AAG CGG
      I   L   F   C   Y   C   S   I   L   A   K   L   G   E   H   Y   G   H   S   N   K   S   S   Q   R   R   E   N   S   V   K   I   V   F   A

1366 ATC GTG TCG GCT TTT ATT TTA TCC TGG CTG CCC TTT AAC GTC TTA AAA TCG ATG GCC CTG TAC CTG CAG TTC CAC AAT GTG GAC CTC TCC TGC TGG CCG CTG GTC
     TAG CAC AGC CGA AAA TAA AAT AGG ACC GAC GGG AAA TTG CAG AAT TTT AGC TAC CGG GAC ATG GAC GTC AAG GTG TTA CAC CTG GAG AGG ACG ACC GGC GAC CAG
      I   V   S   A   F   I   L   S   W   L   P   F   N   V   L   K   S   M   A   L   Y   L   Q   F   H   N   V   D   L   S   C   W   P   L   V

1471 AGC CGG GGT CTG GCC GTG GCT TCC TGC TTG GCT TTC CTC AAT AGC TGC GTC AAC CCC ATC ATT TAT GCC TTC CTG GAC CGC AAT TTC AGG CAG AGG ACC CGC CGG
     TCG GCC CCA GAC CGG CAC CGA AGG ACG AAC CGA AAG GAG TTA TCG ACG CAG TTG GGG TAG TAA ATA CGG AAG GAC CTG GCG TTA AAG TCC GTC TCC TGG GCG GCC
      S   R   G   L   A   V   A   S   C   L   A   F   L   N   S   C   V   N   P   I   I   Y   A   F   L   D   R   N   F   R   Q   R   T   R   R

1576 ATG ACC TCG CAC GTC TTC GCC GGC CTG GGG AAG AGG AGC AGC AGC TTC GGC TCC GGC TCC ACG GTC ACA GAG AGC AGC ACC GCG CTG AAG GTG CAG AGC CCA ACA
     TAC TGG AGC GTG CAG AAG CGG CCG GAC CCC TTC TCC TCG TCG TCG AAG CCG AGG CCG AGG TGC CAG TGT CTC TCG TCG TGG CGC GAC TTC CAC GTC TCG GGT TGT
      M   T   S   H   V   F   A   G   L   G   K   R   S   S   S   F   G   S   G   S   T   V   T   E   S   S   T   A   L   K   V   Q   S   P   T

1681 ATC TGA
     TAG ACT
      I   *
```

**Fig. S2.** The nucleotide sequence and amino acid sequence of the St-GPR25 constructs overexpressed HEK293T cells. The amino acid sequence of St-GPR25 is shown in red, that of LgBiT is in blue, the signal peptide of sLgBiT is shaded.

**LgBiT-MYLK2S in MCS1 of pTRE3G-BI**

```
1       CGC GGG GAT CCA TCG ATC CGC ATG GTC TTC ACA CTC GAA GAT TTC GTT GGG GAC TGG GAA CAG ACA GCC GCC TAC AAC CTG GAC CAA GTC CTT GAA CAG GGA GGT
        GCG CCC CTA GGT AGC TAG GCG TAC CAG AAG TGT GAG CTT CTA AAG CAA CCC CTG ACC CTT GTC TGT CGG CGG ATG TTG GAC CTG GTT CAG GAA CTT GTC CCT CCA
                                     M   V   F   T   L   E   D   F   V   G   D   W   E   Q   T   A   A   Y   N   L   D   Q   V   L   E   Q   G   G

106     GTG TCC AGT TTG CTG CAG AAT CTC GCC GTG TCC GTA ACT CCG ATC CAA AGG ATT GTC CGG AGC GGT GAA AAT GCC CTG AAG ATC GAC ATC CAT GTC ATC ATC CCG
        CAC AGG TCA AAC GAC GTC TTA GAG CGG CAC AGG CAT TGA GGC TAG GTT TCC TAA CAG GCC TCG CCA CTT TTA CGG GAC TTC TAG CTG TAG GTA CAG TAG TAG GGC
         V   S   S   L   L   Q   N   L   A   V   S   V   T   P   I   Q   R   I   V   R   S   G   E   N   A   L   K   I   D   I   H   V   I   I   P

211     TAT GAA GGT CTG AGC GCC GAC CAA ATG GCC CAG ATC GAA GAG GTG TTT AAG GTG GTG TAC CCT GTG GAT GAT CAT CAC TTT AAG GTG ATC CTG CCC TAT GGC ACA
        ATA CTT CCA GAC TCG CGG CTG GTT TAC CGG GTC TAG CTT CTC CAC AAA TTC CAC CAC ATG GGA CAC CTA CTA GTA GTG AAA TTC CAC TAG GAC GGG ATA CCG TGT
         Y   E   G   L   S   A   D   Q   M   A   Q   I   E   E   V   F   K   V   V   Y   P   V   D   D   H   H   F   K   V   I   L   P   Y   G   T

316     CTG GTA ATC GAC GGG GTT ACG CCG AAC ATG CTG AAC TAT TTC GGA CGG CCG TAT GAA GGC ATC GCC GTG TTC GAC GGC AAA AAG ATC ACT GTA ACA GGG ACC CTG
        GAC CAT TAG CTG CCC CAA TGC GGC TTG TAC GAC TTG ATA AAG CCT GCC GGC ATA CTT CCG TAG CGG CAC AAG CTG CCG TTT TTC TAG TGA CAT TGT CCC TGG GAC
         L   V   I   D   G   V   T   P   N   M   L   N   Y   F   G   R   P   Y   E   G   I   A   V   F   D   G   K   K   I   T   V   T   G   T   L

421     TGG AAC GGC AAC AAA ATT ATC GAC GAG CGC CTG ATC ACC CCC GAC GGC TCC ATG CTG TTC CGA GTA ACC ATC AAC AGT GGT GGC GGC TCT GGT GGT GGC AGC GGC
        ACC TTG CCG TTG TTT TAA TAG CTG CTC GCG GAC TAG TGG GGG CTG CCG AGG TAC GAC AAG GCT CAT TGG TAG TTG TCA CCA CCG CCG AGA CCA CCA CCG TCG CCG
         W   N   G   N   K   I   I   D   E   R   L   I   T   P   D   G   S   M   L   F   R   V   T   I   N   S   G   G   G   S   G   G   G   S   G

526     GGT GGT TTG CTT AAG AAA TAC CTC ATG AAG AGG CGC TGG AAG AAA AAC TTC ATT GCT GTC AGC GCT GCC AAC CGC TTC AAG AAG TGA GCC GGC GAT ATC TCC AGA
        CCA CCA AAC GAA TTC TTT ATG GAG TAC TTC TCC GCG ACC TTC TTT TTG AAG TAA CGA CAG TCG CGA CGG TTG GCG AAG TTC TTC ACT CGG CCG CTA TAG AGG TCT
         G   G   L   L   K   K   Y   L   M   K   R   R   W   K   K   N   F   I   A   V   S   A   A   N   R   F   K   K   *
```

**CALM1-SmBiT in MCS2 of pTRE3G-BI**

```
1       TGG AGA ATT CCC CGG GGG TAC ATG GCT GAT CAG CTG ACC GAA GAA CAG ATT GCT GAA TTC AAG GAA GCC TTC TCC CTA TTT GAT AAA GAT GGC GAT GGC ACC ATC
        ACC TCT TAA GGG GCC CCC ATG TAC CGA CTA GTC GAC TGG CTT CTT GTC TAA CGA CTT AAG TTC CTT CGG AAG AGG GAT AAA CTA TTT CTA CCG CTA CCG TGG TAG
                                     M   A   D   Q   L   T   E   E   Q   I   A   E   F   K   E   A   F   S   L   F   D   K   D   G   D   G   T   I

106     ACA ACA AAG GAA CTT GGA ACT GTC ATG AGG TCA CTG GGT CAG AAC CCA ACA GAA GCT GAA TTG CAG GAT ATG ATC AAT GAA GTG GAT GCT GAT GGT AAT GGC ACC
        TGT TGT TTC CTT GAA CCT TGA CAG TAC TCC AGT GAC CCA GTC TTG GGT TGT CTT CGA CTT AAC GTC CTA TAC TAG TTA CTT CAC CTA CGA CTA CCA TTA CCG TGG
         T   T   K   E   L   G   T   V   M   R   S   L   G   Q   N   P   T   E   A   E   L   Q   D   M   I   N   E   V   D   A   D   G   N   G   T

211     ATT GAC TTC CCC GAA TTC CTG ACT ATG ATG GCT AGA AAA ATG AAA GAT ACA GAT AGT GAA GAA GAA ATC CGT GAG GCA TTC CGA GTC TTT GAC AAG GAT GGC AAT
        TAA CTG AAG GGG CTT AAG GAC TGA TAC TAC CGA TCT TTT TAC TTT CTA TGT CTA TCA CTT CTT CTT TAG GCA CTC CGT AAG GCT CAG AAA CTG TTC CTA CCG TTA
         I   D   F   P   E   F   L   T   M   M   A   R   K   M   K   D   T   D   S   E   E   E   I   R   E   A   F   R   V   F   D   K   D   G   N

316     GGT TAT ATC AGT GCA GCA GAA CTA CGT CAC GTC ATG ACA AAC TTA GGA GAA AAA CTA ACA GAT GAA GAA GTA GAT GAA ATG ATC AGA GAA GCA GAT ATT GAT GGA
        CCA ATA TAG TCA CGT CGT CTT GAT GCA GTG CAG TAC TGT TTG AAT CCT CTT TTT GAT TGT CTA CTT CTT CAT CTA CTT TAC TAG TCT CTT CGT CTA TAA CTA CCT
         G   Y   I   S   A   A   E   L   R   H   V   M   T   N   L   G   E   K   L   T   D   E   E   V   D   E   M   I   R   E   A   D   I   D   G

421     GAC GGA CAA GTC AAC TAT GAA GAA TTC GTA CAG ATG ATG ACT GCA AAA GGT GGT GGC GGG AGC GGA GGT GGA GGC TCG AGC GGT GGA GTG ACC GGC TAC CGG CTG
        CTG CCT GTT CAG TTG ATA CTT CTT AAG CAT GTC TAC TAC TGA CGT TTT CCA CCA CCG CCC TCG CCT CCA CCT CCG AGC TCG CCA CCT CAC TGG CCG ATG GCC GAC
         D   G   Q   V   N   Y   E   E   F   V   Q   M   M   T   A   K   G   G   G   G   S   G   G   G   G   S   S   G   G   V   T   G   Y   R   L

526     TTC GAG GAG ATT CTG TAA GTA CCA CGC GTC TGC AGC ATA
        AAG CTC CTC TAA GAC ATT CAT GGT GCG CAG ACG TCG TAT
         F   E   E   I   L   *
```

**Fig. S3.** The nucleotide sequence and amino acid sequence of NanoBiT-based calcium sensor. LgBiT and SmBiT are shown in blue, CALM1 and MYLK2S are shown in red.

### A: Gene position and architecture

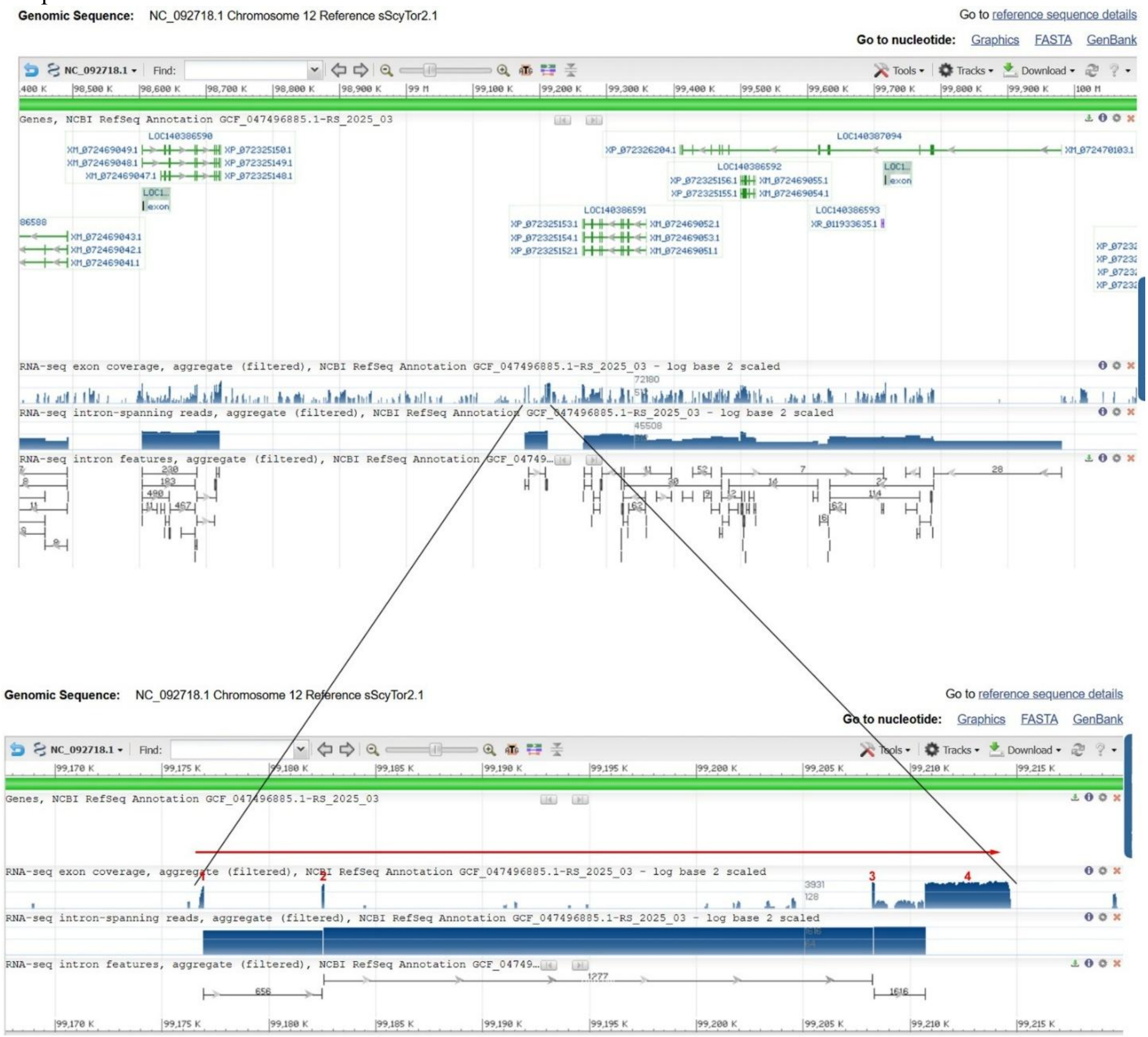


### B: Sequence of genomic DNA (exons in red and highlighted in yellow, coding region underlined)

gcagatttacaggttcccagcaatgccgtgacttgctttcggttggccgggggaggggaaacttgcccctcctcttctctggggagggagcaggtgcagggctgtcggggatctgcatgactgataatgggagagatataagtcttccaacattcagaagcactgattaaactt
tcgatcgtgccacttctggatcagtccttaaatctgaagatgcagctggtttccctgattgctgtgtttgccttggccacaatatctgtcctcacagatggtaagaatttaaatctgtgcaatgatttgctttatggtttacatttagagaaaataatatcacaatgtttattc
agataggtaatttaaaaaatatattatttatatttgcaacgagggtgttgctggttaaaccagggctgaaattctccagttgttgcgattcaattttcgcgccagcaacgtacacccaccaacggtttcacggcagctcgggctagttacgatgggaaatcccattgacaatcg
gtgggaattggggaagctgacatagaggcgcgcagcgacaaactggagaaaccagccccaaattatttgcaaatttttagttacaatagagaggctttctcagtcacttctgtttcagtatctaacagcatatattccaccagcaattctgaatgtagttaacaatgcgcagtt
aaatgaggaataaattccctggctgttcctcttcaaagtaatgtaatgggatacttcatatcaatttgatgaggcagatggtttaaggtctcacccgaaagattgcacctctgacagtgcagtgccccctctgtactgcactggagtgtcagccttaatttccatgcgctcaag
ccttggagtgggatttgaaatcagaatctcggtgttcagaggtgagggtgttaccatctgagctaaggtgtcatgcttgcctctatactaagaatcaatcgcatagttgtggaactggagtctcatttcagccagtccgtttaaggatggtagattttgttaactgaaggacat
ccgtgatcaagttgggtttttacgacaatctggcaatgtcactgtcactttgactcgtaccagttttttctttttatcttcccagacatatctttaaactttatttaaatgctgaacttacaaggatgacatttgaacccaagttacctggggtttaatgtgaacgtgaataat
tgcattcatatagcgcctttcatgaggtttgcatgtcctaaagtgctttctaatttattaattgcattcagagctggagtcactgttgtcctgtaagaaatccaaagctagtttgtacacaacaagctcccagaaacaacaatgtttaagagcagaattgagcggtaaggattg
accacaaaatgggggacaccactgttcttctctgaagagtgaccgtgacatttttttttgtttgttgcaaaaatatactttatttgtaaaatatctgatagaacattacaacatttcaaagtggccatcacagaaagtgcaataatattcaaattttcgacataagtcatgggct
ggattctctggacccccagctgcgtgtttctcgatggtgtgctgttcgctgtcggtggaattttcccttcccgctgcttgtcaatgggatttccaattgaagccagcccgtgctgccgggaatcccattggtgggggtgggttgccagcaggaaaagggaatcccaccaactgg
agtattctggcccaagttgcactctggggcacttcaatacagtcaaagaaacattacagacatttcaaaatggtcataacagcgcaatggtattttagttttctacacaaatcaagttgcattatgaggtgcttcaatacaattctgatacatttaatattcactgtatacatt
cactgtgagtcatacggcccgaggggggttctgtacaattcccttcccttcagttcacttgggctgaaaggtcttaggcaatgaccgtcccccattgcgcctctacggcaggttccctgagcttcagtgcctccctcagcacgtagtcctgggttttggttgtccaactgagagg
ttcaaagtctcatctgaacgatattacctcagccagtgcagctcccccctcagtgctgccctgcagcgctggcctagattatgcgctcaaatctctggagtggatttcatttagggaagagttaagaagcaactgttttgttctggaatgcactgcatgagaggacaatggaagc
agattcaatcgcaacttcagggaattgggtaaatatttaaaggggggaaattcttcttctgactgcccctcacagcaaggagaaacttgccacagagtctagcccatagcaagaacctgcgcaaagagtcatacagttttacaggaagaggcctttggcctatcgtgtctgtgtc
ggccatcgagcaccaatctattctaaacccatttttctgcacttggtctgtagtcttgtatgctatggcatttcaagtgctcatttaagtgtttcttaaatgttgtgagggttcatggctctacaacactttcagtctgtgagttccaggttcccgccatcctctgggtgaaaa
ggactttcctcaaatcccctcaaaatctcctgccttacattaaatctgtgcccctggttatcgacctctctactgaggggaaatgtttgttcctgtctatcctctcgatgtctctcataattgtatacacctcaattggttcccctctgcagccttctctgctctaaggagaac
aacccccagtctgaccagcctctcttctcagctgaaactctccagcccaggcaacatcctggtgaatctcctttgcatcctttccagtgcaatcacatccgtcctgaagtgtggcaaccagaactgtacacagtatctagctgtggccgaatgagcattttatacagatccacc
ataacctgtctgcacttatattctatgcctcggctaattatggcaagtatatcatgtgcctttttaaccaccttatctacctctcctgctgccttcaggggtctttggacactcaagtccctctcgccctctgtacatcctagtgtccgaccattcattgcgtattctcttgct
ttgttagtcctaccaaaaagcatcacctcaccctttttcagggttaaattccatttgctaggcttggtttaatccaattcgaaatagtttatagggcgcacctgagtaagtccagaatgagtcgcttcattgcatgggtggaagacacgtgcgcgctctgctcaaccaggttctg
ctcgagccccaagcttgtgggtttctgggtctccttcttcagcactatgtcggcgattctgacaattcaactggagccctgtcccttggtagccaaattgtgagtgtcagtcttgcccgagctgcagacaggtgtgggggctgatgccttggccttggcctcgctgattgcctg
gaggtgggtactgctggggtggaggtctgcttctccacccagtacctcagtgtggctgggagatctgatggagtttctatacctcgagaaagtcaagtacaccatgcggggagcaattgagggggttttactgaagatgacagccgtctattgtttatttcaaggaattggccat
ggttcattgctcagaaggggcgaggggaggaagaagcaaagggggggaggggttaggttttgtgaggggtgggctatgttttggaggctttatgattgtattagtgttttattgtgttgtgttgttttgtattatgcataaaatgaaaaaataaaaaaaataaaatttttaaattcc
atttgctattgttctgcacatctgcccaacctgtctatattatcctgtaatcacagagtctgttgtgcggcagctaccacaatgttcagtccaatcaggagactatcccactcttcctctctgccttagacttattggacagattttcaggttgacctccttctctccttcgtg
atagtcaaagacttggttagttatagcaatgagcagttggatccatatgaacaagcacctcttccctggagccttatctgaactgaaattgttctgttccgttggcagtgcaccacaatcagatagattacaagtatgagttgtgggtactgttggactcggccattaagtgga
tgtgatctatgaatcaccgctggggttctcaaattctcaaaacatttcagaaactcaaacatggccccccttacaatttgttgtttggtctgtcactactgtatagcaaacctctcctataaatagtattgaatctccaactccaaggtgcatcaagggaccaagcctagaatct
tgccacctttgtggtttaattgcatcttagcttgcaacaagaaatccccagggtcctcctaggggcagggcaagacgacactccagaaaaccccagaaattacgagtcagaaaataagaattcaggaatccatcaaaataaaaggtccatggagcccatttctaccaccttcca
catgtaaatgttgcaatcaatctctgtagatcaatctttaactggaggtgaggcaaagcccaagtggtgcagcgttttgagaactgcagctcctgcccctccgtgttccactcaccacagttttttttactcaaatgggggtgtagctgactgggccaacatttacttctcattc
cttgaattgcccttgaactgagtgacttacgaggccatttcagaggatagttacgagtcacatgtaggccagaccagctagtgatgacaggtttgcttccctgaagtgaacgagatgggtttttacaacaatcatttcatgttattttattcattccagattgttttccactga
attaaattttgactggctgctaaactggaatttggacctgggtccctaatttcctaggtcactggattgcaggttcagtgacaatatcactgcaccaatgcctccttggttaatgttagggtagtcttggccagtgaccgatttccctaagtagtcctgggttaaggggcagga
atcagtcagtgttcttttttcatgttctctgaacaaccctaacaacaacttccaattattttgtgccttttaacagaagaaaaggtcccaccaaactacttcagaggagtgttatatcaataaacaccgtgtcagacaggatgatattggacgcatgatcaaatgtttagtgaa
agtggtagattttcagggaggaggagaggaaagtagagaggaagataggtttacagggagggaacaacaattagtatttaaaatcaaatggtttcgagactaagaatcagtgtaaatctgcaatgactggaacattacaatcaagaaaaaaaattgtcagggcggcacagtggta
aatggttagcactgttgcctcatagcaccaaggacctgggtttgagtcccggctccaggtcactgtccgtgtggcgaatgcacattctctccatgtctgcgtcgttctcacccccacaacccaaagacatgcagggtaggtggattggacacgttaaattgctccttaattaga
aataaaaagaattgggcactttaatcttttttctaaattatcagtatcattgaacaaagtgaaggtctgtcacagcctctattcagtcagtgacatcacttcaaatggactgggcaggtctgacaagcatgtttttttaggggtcaccacaatgacataaaaggtcacaaaaatt
gctttgattatgaatataaagtgccagggtatatattttttagaggcatgtttcttatttccagatacaactcagctgaaagaggatgaggcaaagagggtactgacaacaaccagggtgaaatctcctgccagtgaggagagccgctgtgtgtgcaaaggtaggaaggcacatt

cagacctcgattactcacaaacctgaccaacaagataatgaaatcagagactgggacgaacaaaggcaaaactgagttaaccatgatgcagataaaccagggacagggtgaatagggaaacagttcgaataaaaaggactggataaagatgggaatagttaatccagagattgg
ataaccacgggaacagttaatccagggactggataaacacagaaatagttaatcaatggcttttataaacatgggaatatttaatctgtcgactggataaagacaggaatagtgaatctgtagactggataaagacaggaatagttaatccatagactggagaaacatgggaat
agttaatctgtagactggataaagacaggaatagttaatctgtagactggataaagacaggaatagttaatccagggactggataaacacgggtatagttaatccagggactggataaacatgggaatagttaatctggagactggataaagacaggaatagttaatccagggt
ctgtataaacatgggaatagttaatccaaagactggataaagacaggaatagttaatccaaagactgggtaagcatgggaatagttaatccagggactggataaagacaggaatagttaatccagggtctggatgaacaggggaatagttaatccaatacctagataaagacag
gaatagttaatccaaagacctgataaatatgggaatagttaatccaaagacctgataaatatgggaatagttaatccagggactggataaacacgggaatagttcatctaggggctggatgaacacaggaatagttaatccaaaaacaggataaacacgggaatacttaatcca
aagactggataaacatgggaatagttaatccagggattagataaacacaggaatagttaatccaaaaactagataaacatgggtatagttaatccagggactggataaacacgggaggagttaatccagggactagataaacacgggaatagttaatccaggagttggataaac
acaggaatagttaatgcaaaaactggataaacacgggtatagttaatccagggactggataaacacgggaatagttaatccagggactagataaacacgggaatagttaatccaggagttggataaacacaggaatagttaatccaaaaactggataaacatgggaatagttaa
tccaggtactgaataaacacaggaataattaatccagggattggataaacacgggaatagttaatccaggggctggataaacacaggaatagttaatccaaaaactggataaacatgggtatagttaatccagggactggataaacacgggtatagttaatccatagactggat
aaagacaggaatagttaatccagggattggataaacacgggaatagttaatccagggactggataaacacgggaatattttatccaggtaaaggataaacacaggaatagttaatccagggactggataaacatgggaatagttaatccatggactggataaacacattaatcg
gggcgaaattctcgcccaacggcgcgatgtccgccgactggcgccaaaaatggcaccaattagacggacatcgcgccggcccaaaggtgcggaatgctccgcatctttgggggccgagccccaacattgaggggctagtccggcgccggagggacttccgccccgccagctggc
ggaaatggcgtttgttgccccgccagctggcggaaatggtgtttggtgccccgccagctggcgcacaaatgcggcgcatgcgcgggagcgtcagcggctgccgacagtttcccgcgcatgcgcagtggggagagtctcttccgcctccgccatggtggaggccgtggcggaagg
gaaagaatgcccccacggcacaggcccgcccacggatcggtgggccccgatcgtgggccaggccaccgtgggggcagcccccggggtcagatcaccccacgccctccccaggaccctggagcctgcccacgccgcctggtcccgccggtaaataccagctttgatttacgccgg
cgggacaggcaatttctgggcgggacttcggcccatccgggccggagaattgagcggggtgtctcgccaactggcacggcccgattcacaccccgcccactctccggtaccggagacttcggcgggggcgggggcgggattcacggcggccaacggccattctccgacccggc
ggggggggtcagagtatgacgtctaatttatccaaagactggataaacacaggaatagttaacccaaagacaggataaacatggaaatagttaatccagggactggataaacatgggaatagttaatccatagacaggataaacacgggaacagttaatccagcaactggataa
gcacgggtatagttaatacaggaactggataatcacgggaatagttaatccagggactggataaacaggggaatagttaatccaggggctggataaacacaggaatagttagtccaaaaactggataagcattggtatagttaatccagggcctggatttacacgggtctagat
aatccatagactggataaagacaggaatagttaatccagggactggataaacatgggaattgttaatccaggcactggataaacaaggaatagttaatccaggggctggataagcacaggaatagttaatccaaaaactggataaacacaggtatagataatccatagactgga
taaacacgggtatagttaatccagggactggataaacatgggaatagttaatccagagactggataaacacgggtatagttaatccagggactggataaacaagggaatagttaatccagggactggataaacatgggaatagttaatccatagactggataaacacgggtata
gttaatccagggactggataaacatgggaatagttaatccagggactggataaacatgggaatagttaatccagggactggataaacaagggaatagttaatccagaggctgcataaacacaggaatagttaatccaaaaactggataaacacgggtatagttaatccatagac
tggataaacacgggtatagttaatccagggactggataaacacgggaatagttaaaacatagactggataaacacgagaatagttaatccacggactggataaacacgggaatatttaatacaggggctggataaacacgggtatagttaatccatagactggataaagacagg
aatagttaatccagggactggaaaaacataggaatagttaatccagggactggataaacacgggaatagttaatccaggggctggataaacacgggtatagttaatccatagactggataaacatgggtatagttaatccagggactggataaacacgggtatagttaatccag
ggactggataaacacaggtatagttaatccagggactggataaacacgggaatagttaatcgagggactggataaacatgggaatagtaaatccatagactgtatcgtgacaggaatagttaatccaaagcctggataaacatgggaatagttaatccagggactggataaaga
caggaatagttaatgcaaaaactggataaacacgggtatagttaatccagggactggataaacacgggaatagttaatccaaggactggataaacatggaaatagttaatccagggacctgataaacacgggaatagttaatccagggagtggataaacatgggaatggttaat
ccccgtaccagcctccccgaacaggcgccggaatgtggcgactaggggcttttcacagtaacttcattgaagcctacatgtgacaataatcgattttcattttcattaatccatagactggataaacacaggtatagtttatccatagagtggataaacatgggaatagttagt
ctagaatctggataaacacgggaatagttcatccagagtcaggaaaacacgggtatagttaatccagggactggatgaacacgggaatagttaatctgtagactggataaacacaggaatagctaatccataggctggataaacaggggaatagttaatctatagactggataa
acatgggaatagttaatctgtagactggataaacacgggaacagttaatccagcaattggaaaaacacgggaatagttactccggagtctggatcaacacggatatagttaatccatagactggataaacatgggaatagttaattcagcaactggataaacacgggaatagtt
aatccagagtctggatcaacacgggtatagttaatccatagactggataatcatgggaatagttaatccagcaactggataaacgcgggaatagttaatccagagtctggatcaacacgggaatagttaatccagagtctggatcaacacgggaatagttaatccagggactgg
ttaaacacgggtatagttaatccaggagctggataaacacgggtatagttaatccatagactggataaacatgggaatagttaatccagggactggatacacacgggaatagttaatccaggagctggataaacacgggaaaagttattccaaagactggatcaacacggttat
agttaaggcagggactggataaacacgggaatagtaaatctgtggactggataaacacaggaatagtcaatccgtaggctggataaacacgggaatagttaatctatagactggataaacatgggaatagttaatccagagtcaggatcaacaccggtatagttaatccaggga
cttgagaaacacgggaacagttcatctgttgactggataaacacgggaatagttaatccatagactggataaacacgggtatagttaatccacggactggataaacacgggaatagttaatccagggactggataaacacaggtatagttaatccatagactggataaacacgg
gtatagttaatccacggactggataaacatgggaagagttaatccagaatctgaatcaacacgggaatagttaatccagggactggataaacacgggtatagttaatccaggtgctggataaacacgggtatagttaatccatagactggataagcatgggaatagttaatcca
gggactggataaacacgggaatagttaatccaggggctggataaacacgggtatatttaatccatagactggaaaaagacgggaatagttaatccagggactggataaacacgggaatagttaatccaggggctggataaacacgggaatagttattccaaagactggatcaac
acggatatagttaaggcagggactggataaacacgggaatagtaaatccagagactggataaacatgggaatagtaaatccagggacacgataaacacgggaatagttaatccagggactggataaacacgggaatagtaaatctgtggactggataaacacaggaatagttaa
tccgtaggctggataaacacgggaatagttaatctatagactggataaacatgggaatagttaatccagagtctggatcaacacgggtatagttaatccagggacttgagaaacacgggaacagttcatctgttgactggataacacgggaatagttaatccatagactggata
aacacgggaatatttaatccagggactggataaacatgggaagagttaatccacagactggataaacacgggaatagttcatacatagactggataaacacgggaatagctaatccatagactggataaacactggaatagttaacccatagactggatgaacacgggtatagt
taatccagggactggataaacacaggaatagttaatccatagacttgataaacacgggtatagttaatccagggactggataaacatgggaagagttaatccacagactggataaacacgggaatagttcatacatagactggataaacacgggaatagctaatccatagactg
gataaacactggaatagttaacccatagactggatgaacacgggtatagttaatccttagactggataaacacggaaatagttaatccataaactgtttaaacacgggaatatttaatccagggactgtctaaacacgggaatagttaatccatagactggataaacacaggaa
tagttaatccatagattgaataaacatgggaatagttagcccagggactggataaacacggatacagttaatcctgggactggataaacatgggaatagttaatccagggactggataaacacggatacagttaatccatagactggataaacatgggaatagttaagccaggg
actgtataaacatgggaatagttaattcagagactggataaacacgggaacagttaatccgtagactggataaacatgggtatagttaattcagagactgtataaacacggatacagttaatccagagactggataaacatgggaatagttaatccagggactggataaacacg
gatacagttaatccattgactggataaacatgggaatagtcaatccagggactggataaacatgggaatagttaattcagagactggataaacacgggaatagttaatccatagactggataaacatgggaatagttaattcagagactgtacaaacatggatacagttattcc
atagactggataaacatgggaatagttaatctagggactggataaacatgggaatagttaatccagggactggataaacatgggaatagttaatccagagactggataaacaagggaatagttaatccaggggctgcataaacacaggaatagttaatccaaaaactggataaa
cacgggtatagttaatccatagactggataaacacgggtttagttaatccagggactggataagcacgggaatagttaaaacatagactggataaacacgagaatagttaatccacggactggataaacacgggaatatttaatacaggggctggataaacacaggtatagtta
atccatagactggataaagacaggaataattaatccagggactggaaaaacataggaatagttaatccagggactggataaacacgggaatagttaatccaggggctggataaacacgggtatagttaatccatagactggataaacacgggtatggttaatccaggaactgga
taaacacgggtatagttaatccagggactggataaacacgggtatagttaatccagggactgaataaacacgggtatagttaatacatagactggataaacacgggtatagttaatccattgactggataaacacgggtatagttcatccatagactggataaacatgggaata
gttaatccagggactagataaacacgggaatagttaatccagggactggataaacatgggaatatttaatccagggactggagaaagacaggaatagttaatcgagggactggataaacatgggaatagtaaatccatagactgtataatgacaggaatagttaatccaaagcc
tggataaacatgggaatagttaatccagggagtggataaagacaggaatagttaatgcaaaaactggataaacacgggtatagttaatccagggactggataaacacgggaatagttaatccaaggactggataaacatggaaatagttaatccagggacctaataaacacggg
aatagttaatccagggagtggataaacatgggaatagttaatacccgtaccagcctccccgaacaggcgccggaatgtggcgactaggggcttttcacagtaacttcattgaagcctccttgtgacaataatcaattttcattttcattaatccatagactggataaacacagg
tatagtttatccatagactggataaacatgggaatagttagtccagaatctggataaacacgggaatagttaatccagagtcaggataaacacgggtatagttaatccagggactggatgaacacgggaatagttaatctgtagactggataaacacaggaatagttaatccat
agactggataaacacgggaatagttaatctatagactggataaacatgggaatagttaatctgtagactggataaacacgggaacattgtggcgcacaaagactccgtgagacaaatagggtgaagtcgatgaggctttattaagcgtgtctgttccccagcagcccgatagta
aactggcctgcgggggaaggcaccggcttcttatacttcgccttcagggcggagcatgaggtcaacggccaaccaggacccgggatctgtcagccaatgacattagggcttccagtcccacatgacccccaatacatactaccacattcaccccttgtcaaaatgaacccggcg
gggtgatgcttcgtatggtggtaagggtttacagtactggtcctgggaggatagaacagttacatggtagtaccgtattggacagtatcgtcctgttacaactatttacaggaggatataggaaaaacaaaatgttcattggacagtccatctttgttttacatcgacgccacg
agtcggtcgggcggtctggtcgtccgtgtcgatcgcctcggaccggcggtggtggtggtgtttgtaccagcgttgtcgcctccgggagccgcacggtttcagctgtcagtgcaaaggggaggggaactgatcctcctgggaaggggcggtcgcggggtgcggcggtggcagga
aggtgggcggttgggttgatggtgccgggggggtgtgcgtggtgccggcgggcgccagatcccgcagggagactgtgtcctgtcggccgtcggggtactccacgtaggcgtactgggggtttgcgtggaggaggtgaaccctttcgaccaacgggtccgccttatgtgcccgca
cgtgctttcggagcaggatgggtcctctggccgccagccaggtcggcagcgacgttccagaggaggacctcctagggaagacaaggagacactcgtgaggcgtttgattagtgcttgtacataataacgaccggatggaatggagagcgtccgggaggacctcctgccaccgtg
aaactgggagatccctggaccgtagggccagtaggacggccttccagaccgtgccgttctccctttctacttgcccgttcccccgggggttgtagctggtcgtcctgcttgaagctataccctttgctgagcaggaactggcgcagctcgtcactcatgaaagaggaccccctgt
cgctgtggacgtatgcggggtaaccgaacagtgtgaatatgctgttcagggctttaatgactgtggccgcggtcatgttggagcagggaatggcgaaagggaagcgggagtattcgtccactacattaaggaaatacgcgttgcggtcggtggagggaggggcccctttgaaat
cgagactgaggcgttcaaagggacgggaagccttaatcaggtgcgctccatctggcctgaaaaaatgcggtttgcattctgcgcagatgtggcagtctcttgtgactgtacggacctcctctaaggagtatgggagattgcgggacttgatgaagtggtaaaaccgagtgaccc
ccgggtggcagaggtcctcgtggagggcttggaggcggtcaatttgtgcgttggcacatgtcccgcgggatagggcatcagacggctcgttcagctttccgggccggtacaagatctcatagttgaaggtggagagttcgatcctccaccgcaagatcttgtcgttcttgatct
tgccccgctgtgcattatcgaacatgaaggctaccgaccattggtcagtgaggagagtgaatctcctgccggccaggtaatgcctccaatgtcgcacagcttccactatggcttgggcttccttttccactgaggagtggcggatttctgaggcgtggagggttcgggagaaga
aggccacgggtctgcccgcttggttaagggtagccgcgagagctacgtcagaggcgtcgctctcgacctggaaggggagggactcgtcgatggcgcgcatcgtggcctttgcgatgtccgctttgatgcggctgaaagcctggcaagcctctgtcgacagcgggaaagtcgtgg
tctgtattagggggcgggccttgtctgcgtactggggacccactgggcgtagtatgaaaagaaccccaggcagcgtttcagggcttttgggcagtgcgggaggggaaattccatgagtgcgcgcatacgttcggggtcggggcctattatcccattgcgcactatgtagccca
gaatggctagccgatcggtgctaaaaacgcacttgtcctcgttgtacgtgaggttcaaggctttggcagtctggaggaatttttggaggttggcgtcgtggtcctgctgatcgtggccgcagatggttacattgtcgaggtacgggaacgtggcccgtaacccatgttgatcaa
ccattcggtccatttcccgttggaagaccgagacgccgtttgtgacgccaaaagggacccttaggaagtggtataatcgcccgtctgcctcgaaggctgtgtacttgcggtcacttgggcggatggggagctgatggtaggcggacttgaggtccacggtggagaagactttat
actgggcaatccgattgaccatgtcggatatgcgggggagagggtacgcgtctagttgtgtgtacctgttgatggtctggctatagtcaatgaccatcctttctttctcccctgtcttcactactaccacctgcgctctccaggactattgctggcctggattatgccctcct
ttagtagccgctggacttcggagcgaatgaaggtccggtcctgggcgctgtaccgtctgctcctagtggcgacgggtttgcaatccggggtgaggttcgcaaacaaggacgggggttgcaccttgagggtggcgaggccgcagatagtgagtggggtatggggccgccgaatt
taaacgtaaggctctgtaggttacattgaaaatctaagcccagcaaggtgggggcacagagttggggaaggacgttaagtttgtagtttttgaattccctcccctgtaccgttagggtaactatgcagaatccctggatctgtacggagtgggatcctgcagctaggcaaatct
tttgtgcgctggggtaggtagttaaggaacagcgtcttaccgtgtcgggatgtataaaactttccgtgctcccggagtcgactaggcatggcgtctcgtggccgttaattaggaccgttgtcgtcgtcgtctggagagtccggggccgagcctgatcgagtgtaaccaaagcga
gccgtggttgtagtagtggggtggggtccgttgttgccgtccaagatggcgtcggggatggacaaaatggccaaccccatacctcgcccgtagctgaggggtcacaagatggcgtcgggggtggacaaaatggccgcccccatgcgtcgtacaggtcggggcggtccaagatg
gcggggcccctcctcccctcgtggtggtcggaccaaaatgacggcgtctgcgggtcgcacattgagtgctgggggtctgggggcgctgggaccgcgcggagttcctgcattgcgagcgcccgggccgcgggcgctaggaccgcgcggagttccttggctgcgagcgccagg
gccgcgggcgctaggaccgcacggagctccctcgctggggaccgcggtggtcggggccaaggccggcgttcagtactgcgagcgctgggaccgtgaggaggtccctcgccggggacggcgatcggggtccaggtaagttgtattgccggcgggtcaggcgtggggtgcgggacg
ggggtgagggcccaggtcggcggcgttggcgggtcggcgtggtgcgcgggacgggggcgagggcccaggttggcggctccggcgggtcaggcgtggggcgcgggacgggggcgagggcccaggtcggtggcgctggcgggtcaggcgtggggtgcgggacgggggcgagggcc
caggacggcggcgctggcgggtcaggcgtggggcgcgggacgggggcgagggcccaggtcggcggctccggcgggtcagtcgtggggccgcgagcgctggcaccgcgcgcagctccctcgccggggacagcggtggtcggggtcaaggtaggtggcgccgacgggtcaggcgtg
gggcgcgggacgggggtgagggcccaggtcggcggcgctggcgggtcaggcgtggggcgcgggacgggggtgagggtggcgttcgggatgcgaaaaacccccctcctctccgtggagagtgttggccggcacccaaatcgggagcgccggcggcgggtcgtacattgtctgcggg
gaggggggttggggagcgtaggcggcgtgcagggctcccagttctcccggggccgcggtggtcgggacccaaaatggcggcgcctgggggtcgcacatggcgtgctggggggggttgggaggcatagactgcctgtagggctccctgttctcccgggacggcggcgaccacgcgg
gaccggcacaccaccgcatagtggccctttttcccgcagcttttgcagatggctgcgcgggccggacagcgttgtcgggggtgcttcgcctggccgcagaaataacagcgggcgcccccggtgcgactcggcgtttggactgcgcaagcctgtggggtgtccgggggggggggg
gggggtagggggtttgccgcggcgtgtacgtacggggcccaatggcctgccgcgcggtcggggccgtcggcgcgggtattacgcgcggctacgtccagcgaggctgccagggcccgtgcctctgagagtcctcgtgactctttttccagaagtctttggcggatttgggaggat
tgtatacctgcaacaaaagcatcgcgcattaacatgtccgtgtgttcgtttgcgttcaccgacgggcagctgcaggctcgtcccaaactcaacagtgcggcgtagaactcgtccatcgattctccgggagcttgccgtctcgtcgcgagctggtagcgagcgtagatttggtta
actgggcgaacgtagagacttctcagtgctgtgaaggccatctggaaatcgcgggtgtcttcaatgagggtgaaaatctccgtgctcaccctcgagtgcaggacctgcagtttttgttcatctgagactcggcctgtggtcgatgtgatgtaggcctcaaagcaagtctgcctg
tgtttgaaggctgctgccgcattcactgcgtgggggctgatcctcaggcattctggaatgatcctgagctccatagtcctttttaggcacgcttaataaattgtggcgcacaaagactccgtgagacaaatagagtgaagttgatgaggctttattaagcgtgtctgttcccca
gcagcccgatagtaaactggcctgcgggggaaggcaccggcttcttatacttcgccttcagggcggagtatgaagtcaacggccaaccaggacccgggatctgtcagccaatgacattagggcttccagtcccacatgacccccaatacatactaccacaaacagttaatccag
caattggaaaaacacgggaatagttaatccagggtctggatcaacacggatatagttaatccatagactggaaaaacatgggaatagttaattcagcaactggataaacacgggaatagttaatccagagtctggatcaacacgggaatagttaatccagggactgaataaaca
cgggtatagttaatccaggggctggataaacacgggtatagttaatccatagactggataaacatgggaatagttaatccagggactggataaacacgggaatagttaatccaggggctggataaacacaggtatatttaatctatagactggataaagacaggaatagttaat
ccagggactggataaacacgggaatagttaatccaggggctggataaacacaggtatatttaatctatagactggataaagacaggaatagttaatccagggactggataaacacgggaatagtaaatccagagactggataaacacgggaatagtaaatccagggacaggata
aacacgtgaatagttattccagagtctggatcaacacgggtatagttaatccatagactggataaacatgggaatagttaatccagcaactggataaacacgggaatagttaatccagagtctggatcaacacggcaatagttaatccagggactggataaacatgggaatagt
taatccatagactggataaacacgggtatagttaatccaggggctggataaacacgggtatagttaatccatagactggataaacatgggaatagttaatccagggactggataaacacgggaatagttaatccaggggctggttaaacacgggtatatttaatcatagactgg
ataatcatgggaatagttaattcagcaactggataaacatgggaatagttaatccagggactggataaacacgggaatagttaatccagggactggataaacacgggaatagttaatccaggggctggataaacacgggaatagttattccaaagactggatcaacacgtatat
agttaaggcagggactggataaacacgggaatagtaaatccagagactggataaacacaggaatagttaatccatagacttgataaacacgggtatagttaatccagggactggataaacatgggaagagttaatccacagactggataaacacgggaatagttcatacataga
ctggataaacacgggaatagttaatctatagactgggtaaacatgggaatagttaatccagagtctggatcaacacgggtatagttaacccagggacttgataaacacgggaatagttcatctgttgactggataaacacgggaatagttattccagagtctgaatcaacacgg
atgtagttaatccgtagtctggataaacatgggaatagttaatccagcaactggataaacatgggaatagttaatccagagtctggatcaacacgggtatagttaatccatagactggataaacatgggaatagttaatccagcaactggataaacacgggaatagttaatcca
gagtctggatcaacacggcaatagttaatccagggactggataaacatgggaatagttaatccatagactggataaacacgggtatagttaatccaggggctggataaacacgggtatagttaatccatagactggataaacatgggaatagataatccagggactggataaac
acggcaatagttaacccaggggctggttaaacacgggtatatttaatccatagactggataatcatgggaatagttaattcagcaactggataaacatgggaatagttaatccagggactggataaacacgggaatagttaatccagggactggataaatacgggaatagttaa
tccagtggctggataaacacgggaatagttattccaaagactggatcaacacggatatagttaaggcagggactgtataaacacgggaatagtaaatccagagactggataaacacaggaatagttaatccatagactggataaacacgggtatagttaatccagggactggat
aaacatgggaagagttaatccacagactggataaacacgggaatagttaatccatagactgtttaaacacgggaatatttaatccagggactggctaaacacgggaatagttaatccatagactggataaacacaggaatagttaatccatagactgaataaacatgggaatag
ttagtccagggactggataaacacggatacagttaatccatagactggataaacatgggaatagttaatccagggattggataaacacggatacagttaatccatagactggataaacatgggaatagttaatccagggactggataaacatgggaatagttaattcagagact
ggataaacacgggaatagttaatccatagagtggataaacatgggaatagttaattcagagactgtatatacacggatacagttaatccagagactggataaacatgggaatagttaatccagggactggataaacacggatacagttaatccatagactggataaacatagga

atagttaacccagggactggaaaaacatgggattagttaattcagagactggataaacacgggaatagttaatccatagactggataaacatgggaatagttaattcagtgactgtataaacacggatacagttaatccatagactggataaacatgggaatagttaatccagg
gactggataaacatgggaatagttaatccagagactggataaacacgggaatagttaatccatagactggataaacacaggtatagttactccatagactggataaacatgggaatagttaatccagggactggataaacgcgggaatagttaatccagggactggataaacat
gggaatagttattccatcgactggataaacatgggaatagttaatccatagactggattaacatgggaatagttaatccagagactggataaacacgggaatagttaatccatagactggataaacacaggtatagttactccatagactggataaacatgggaatagttaatc
cagggactggataaatgcgggaatagttaatccagggactggataaacatgggaatagttattccatcgactggataaacatgggaatagttaatccatagactggattaacatgggaatagttaatccagggactggataaacatgggaatagttattccatcgactggataa
acgcgggaatagttaatccagggactggataaacatgggaatagttattccatcgactggataaacatgggaatagttattccatcgactggataaacatgggaatagttaacccagagactggataaagatgggaagagttaatctatggactggatattcaaggaaatcgtt
aattcaacaaagaacaaagaacaaattaaatgtccagcacaggaacaggccctgcggccctccaagcccgtgctgaccatgctgcccgactaaactacaatcttctacacttcctgggtccatatccctcaattcccatcctattcatgtatttgtcaagatgccccttaaatg
tcaccatcgtccctgcttcgaccacctcctccggcagcgagttccaggcacccactaccctctgcgtaaaaaacttgcctcgtacatctactctaaaccttgccccactcaccttaaacctatgcccccgagtaattgacccctctaccccagggaaaagcctctgactatcca
ttctgtctatgcccctcataattttgtagacctctatcaggtcgcccctcaaccttcttcgttccaatgagaacaaaccgagtttattcaaccactcctcatagctaatgccctccataccaggcaacattctggtaaatctcttctgcaccctctctaaagcctccacatcct
tctggtagtgtggcgaccagaattgaacactatactccaagtgtggccgaactaaggttctatacagctgcaacatgacttgccaattcttatactcaatgccccggccaatgaaggcaagcatgccgtatgccttcttgactaccttctccacctatgttgcccctttcagtg
acctgtggacctgtactcctagatctctttgattttcaatactcttgaaggttctaccaatcactgtatattccctacctgcattagaccttccaaaatgcattacctcacatttgtccggattaaactccatctgccatctctccgcccaagtctccaaacaatctaaatcct
gctgtatcctctgacagtcctcatcgctatccgcaattccaccaacctttgtgtcgtctgcaaacttactaatcagaccagttacattttcctccaaatgatttatatatactacaaacagcaaaggtcccagcactgatccctgcggaacaccactggtcacagccctccaat
tagaaaagcatccttccattgctactctgccttctttgacctagccagttctgtatccaccttgccagctgacccctgatcccatgtgacttcaccttttgtactagtctaccatgagggaccttgtcaaaggccttactaaagaccatatagacaacatccactgccgtacct
gcatcaatcatcttagtggcctcctcgaaaaactctatcaagtaagtgagacacaacctccccttcacaaaaccatgctgcctctcactaatacgtccatttgcttccaaatgggagtagatcctgtctcgaagaattctctccagtaatttccctaccactgaagtaaggctc
actggcctgtagttccctggattatccttgctacccttcttaaacagaggaacgacattggctgttctccagtcctccgggacatcacctgaagacattgaggatccaaagatttctgtcaaggcctcagcaatttcctctccagcctccttcagtattctggggtagatccca
tcaggccctggggacttatctactttaatattttttaagacacccacacctcgtctttttggatctcaatgtgacccaggctatctacacacctttctccaggctcaacatctaccaatttcttctctttggtgaatactgatgcaaagtattcatttagtacctcacccattt
cctctggctccacacatagattcccttgcctatccttcaatgggccaaccctttccctggctaccctcttgctttttatgtacgtgttaaaagccttccttaaccctatttgccaatgacttttcatgacccccttctagccctcctgactccttgcttaagttccttcctactt
tccttatattccacacaggcttcgtctgttcccagccttttagccctgacaaatgcctcccttttctttttgacgaggcctacaatatcactcattatccaaggttcccgaaaattgccgtatttatccttcttcctcacaggatcatgccggtcctgaattcctttcaactga
cacttgaaagcctcccacatgtcagatgttgatttgccctcaaacatccgctcccaatctatgttcttcagttcccgcctaatattgttataattagccttcccccaatttagcacattcatcctaggaccactcttatccttgtccaccagcactttaaaacttgctgaattg
tggtcactgttcccgaaatgctcccctaccgaaacatctaccacctggccgggctcattccccaataccaggtccagtaccgccccttccctagttggactgtctacatattgttttaagaagccctcctggatgctccttacaaactccgccccgtctaagcccctggcacta
agtgagtcccattcaatattggggaagttgaagtctcccatcaccacaaccctgttgtttttactcttttccaaaatctgtctacctatctgctcctctatctcccgctggctgttgggaggcctgtagtaaacccccaacattgtgactgcacccttcttattcctgatctct
acccatatagcctcactgccctctgaggtgtcctcccgcagtacagctgtgatattctccctaaccagtagcgcaactccgccaccccttttacatcccccctctatcccgcctgaaacatctaaatcctggaacgtttagctgccaatcctgctcttccctcaaccaggtctct
gtaatggcaacaacatcatagttccaagtactaatccaagttctaagtttatctgccttacccgtaatacgtcttgcattaaaacatatgcacttcaggccaccagacccgctgtgttcagcaacttctccctgtctgctctgcctcagagccacactgtccctattccctagt
tctccctcaatgctctcaccttctgacctattgctcccatgcccacccccctgccatactagtttaaaccctcccgtgtgacactagcaaacctcgcggccaggatatttatgcctctccagtttagatggaacccgtccttatataggtcacacctgccccggaagagctccc
agtggtccaggtaacggaaaccctccctcctacaccagctgtttagccacatgtttagctgctctatcttcctatttctagcctcactggcacgtggcacagggagtaatcccgagattacaaccctagaggtcctgtattttaactttctgcctagctccctgaactcctgct
gcaggacctcatgccccttcctgcctatgtcgttagtaccaatatgtacaatgacctctgcctgtttgcccgccccccttcaggatgccctctacccgttcggagacatcctggaccctggcaccagggaggcaacataccatcctggagtctctttcacgtccacagaagcgcc
tatctgtgcccctgactatagagtcccctattactattgctcttctgcgctttgaccctcccttctgaacatcagagccagccgtggtgccactgctctggctgctgctgttgttttcccctgataggctatcccccccgacagtatccaaaggggtatacctgttcgagaggg
ggacaaccacaggggattcctgcactgactgcctgccctttctggtggtcacccatttctctgcctgcaccttgggtgtgaccacatttatataactgcgatctatgacgctttccgccacctgcatgctcctaagtgcatccaattgctgctccaaccgaaccatgaggtctg
tgaggagctccagttgggtgcacattctgcagatgaagccatccgggacgctggaagcctcctggacctgccacatctcacagtcagagtactgcacccctctaactgatattgcgtcaattaattaaaattaaaagtttttaaaaaaaatgtatatatttttttatttttta
aagttactgttaactatctgtttcctcgcactagatttctaatataaatgcgaaagctaaatatagtactctccgatctctggcttagatactcctctaaattatagttaagtaattatgtttaattagttaccaatgcttaatttttttaaatttaatgtagattcccaacca
gccactcaggtcacagcttttctgtgatgtctcttcagtttcccccgacacacacaatttgaaaaaggtataaaagtaaaaatgagtaaaaatcacttacttaccttcttaccttctgagtgtcttagatgttctcaggttctctccctgacagagactgctcctcctcc
gaacggctcccgaaacgaggccgcgatcttttaaaatctccgctctgactcgcagctcctgcgctttttaaatctcctgcgctttttaaatctcaggggctggataaacatgggaacagttattccatagactcgataaacacaggaatagttaatccagagtctggataaaca
caggaatagttcatcccgagtctggataaacatgggaatggttaatccaggagttggctaaaaacaggaatagttaatccagagtctggataaacacgggaatagttaatctcgagattgaatgaacatgggggcagcacggtagcacaagtgattagcactgtggcttcacag
cgccagggtcccaggttcgattccccgctgggtcactgtctgtgcggagtttgcacgttctccccgtgtccgcgtgggtttcctccgggtgctccggtttcctcccacagtccaaagacgtgcaggttaggtggattggccatgctaaattacccgtagtgtccataagggttg
ggaggggttattgggttgcggggataaggtggaagtgagggattaatgtgggtcggtgcagactcgatgggccgaatggcctccttctgcactgtatgttctatgtaatctatgtaatgggaatagttaattcaaagactggataaacttgggaatagtaatccagaaactgga
taaatggtttatagttaatccagagactggataaacatgccagctgttagtccagggactgggtgaatacagaaatagctaatccagcacccaatggagttaccgattgcaactctgcaggacaccaggaactcccccagccccctccccccagtggtccattctcaggtgcac
tgtctgcatgtaagcttattccatataaggtgatcttggtgccagaatgtctggcttcacctgggatatccacgtcaattgatccatttgaaaggggtcaattatctcctgatgccccgtttacaggactagtccagacttgctgcagctgtcggtctctgcacagtaaatttg
agtttgcctgtttgacttcccatcaggcccgtctatggttctgactgtggttttaatttaaagtgtccaattctgtattggtcctaagattcaggccattggccgttttaccacagtccctcctcttgatccaatgagcgtcagcgaactggatcacacaactctaaccaaatc
cccagacagggaagtaccagaatcacaggcaaacaactctaatctctaacccaatatatatacattctgtaggcatcataaatgaaatacaaaattagactactcatatatcaaaattaactgtaggaaatgtagatatgaccaaaatcaggcattttaaatggtctgttgtgc
aggctctattctcttatcctgcctggtgcaggacctttactgcagcctgcaagttgctacatttctgctttagttcctctactttttcttgcaccgcatcccgatcatgatcgcgcgctgccctttgtggtacaccttcagcactgagcatgaaactggctcataagtagaagg
ctggtttggttcctctgttctcaccccttgaactcgctttgcagtttctctatctgctatcttgcctcacccaattcctgtttaaatttcaaactctccgcttctctctcgcggagcggagcgatcacaatcacttcgcttaatttggaagcccatgcgcgtggtcccaactaa
tttttcccttggatcccagatctgctatgggggtgctcccatgttacactcacttgggccgtttcttattgaaatatcttctgagtgtttcttccctttcgacgtggcttgccattttaactctctctccctctctctctctcgctctctctctctccccttgggtttat
cacaggtgtgtaggaatcaatcagggtggcggttcggctttcacagaattccctcccacagagtttacccagcacccttgactatcctggttctgactacatagtttgttcacctggtgttacagacatctggtttatggcccaattactgttggatgcctctactctcgtcca
tccacttgaattctggctatcacagagggctactgctctcttcattcaggctcaagctggatctggataatacaaggttagtgggtttgaatttggagcaactaggtaacttcttatcgaagaccacacagtggggcaccaaatgttgtctgtaaaattcagacatatcgacca
gtttattattcaaatgtttggcccaggtgtccttatttctgagatgaacaaaagacagacacaggttaacaattcaacccaagtttattttcaaacacaatggccgggattctccgacccggcgccgggtcggagaatccccgagggagcgcgggaatcgcgcccggccgcccc
gacgccggctttcccattctccaacgccgattttcgggcacccgcagaattgccgccacgccggtcgggggccgttgacagcagccccctccagggattcttcggccccgaagggccgagtggccaacgagtttggccgagtcccgccggcgtgtgttacgcaggtcccacag
ggcgggatctggaaggtgtgtctgcgggagccgtcctggagtggggcggggtgatccgaccccgggggggggcccccacggtggcctggcccgcgatcggggcctaccgatcggcgggcgggctggttccatggggcctactttactccgcgccgggcccctgtcggctctg
ccatattgcccgggggccagcacggagaagggaacccccacacatgcgcggaagtatgctggctgttccgcgcatgcgcaaactcgtgcgggccgttccgcgccggctggagctgcccgctaggaaggggagaattcctcactttcagggatcattgacgcaggagtggttggc
gccggttttcacgccagcttgggagcatagccccattattagagaatccaccggtcttctccggtgcctctgcatcaagtgtttgctggggaggatctctagatgcctccttcttatccccccacttcacacttccagaagcttctctggccctcaagcaatgtggtcatggtgc
agggatcgcaacacccaatggggttgtcgattgcaactctggaggactcctggacatctcctcagaggtccatcctcaggtgtattatctgcatgtaaccttatcccatatcaggtgatctaggtgccagaacatctggcgtcatctcggaaatccataacaattgaaccattt
gaatggggctaattaacttctaatgccctgtttacaggacaagctcagacttgccccagctgtttctggctatttgacttcccatcaggcccgtctgtgattctgactgtggctttaatttgaagtgtccaattcggtatcaggcctaagattcaggacattgaccattttacc
acgacgggaagagttaatccagggtctgggtaaacacaggaataattaatgcaggaagtcaataaacacaggaagagttattttggggattagataactaagggaatagttaattcagggacttgataaacatgggaatagttaatcctgggactcaataaacatgggaatagt
taatccagagactggacgaatacaggaatagttaaaacaggagttatataaacatgggatagttaatccaaggattggatcaacacgggaatggttaattcagggactcgatgaacatgggtatacctcatccagggaattgctaaatatgagaatagttaagccagagatggg
gcaaacacaggaatagttaattcagagatagtataaattcaggaaaagtccagagccgggatagatgtgggaatagttaagacaggggcagtattaacatgggaacagttaaaccagggactgggaaatatgttgaactgagcgctgatccctgattctgtgttcactattaaa
cgttggtctgaatcacaccaaactgactacaaatttttatcccattcttagttggccacctcgagtttggatacaaagtgaagcagtcccacgatcgaactgggaacaagtgccccctgtcagaaggccataaacatcatcaagaggaaatggaaagatggtaatttctgaaagt
aaaaatagagtagaagtcagtcaattgaaacataatgtgtggttggtgtagcatgcttgagaggtgcagatgactgcggacctatggttcccagagctctccatcactggacaatgtccccttttgctcatcatagaatccttacagtgcagagggaggccacatagcccatc
gagtctgcaccgaccttttgaaaaaccactgtacctaggctcaaccccaggtatatgggatcttgctgtgtgtgaattggctaccgcttttcctaaacagtgactacaattcaaaaagtctgccactggctataaagtgctttgggaaatcttcatttcgcgaattgtctgtat
aaactgatgttttcctccacctgggactgtaggtgtctcattgagagacattgagatccactattagtcaagccttgtcatgtccaaacaaaagagtgaataaatgactgagtaatctgtttctggtggtgttgactgagagtccaggctcccaggatattttcttcataatta
aataatcacttctgtccaactgagtaaggacagaggatcttaaattaatctcatctgaacaacacgccaaacagtgtggtgcaccctcagtactgcaccgaggtgtcacaactgatcaggtgatcacaacctggagtggcatttgaacctgtaacgttctgattcatgtaccga
tatgagctgaaccaagttgacaagacacctcaccaatggcctgtcatgtatgagacccctcaaaaaccctctcatcctttaaagggagtgttccttggttgggaggtgtagcttttaaagagaaaacactccattcaatgggctggggaagaaaacccatttgtaatttgagtg
tttggcattcgagaataacgattgtcatgaaaaagttgactggttgaccaatggcctattcagtctggcctacacctgatgtaaagccctcagaattggtccaatcagcctaccattcccatcaactcctcagagtaactgggcaatacgtctggaaggacggtggcaatgtcc
tgcaaaacccagacaccaacaataaccactcaccagtgataaacaacaagtgccagtcccttgcccattgtgcccgaaaactgaacattgccaaaaaaatgggggggaggaagaaccaagtgcacctttgacattgtaaaacatctcacgggtgcttcacagaaaaggtagcaga
caaaatctgatgccaaaacacagaaaggaatcagaatatgtggttaaaccttggtcaaagagcaagttaagggagagagagaggttagagagccatagaggtttaggaagagaatcctagaccttagaccccaagtggccaaggggtgatgggcaggagggcagatttggagga
tggcttggaaagataaaatgtgacttgaacgtatttatttggaggttacaggtgctgttggcgaggtcagcatttccatcgaatgcatagaatccctacattgcagaaggaggtcattcggcatatcgagtctgctctgaccctgcgaaagaccacccgacctgtgctcaatcc
ccacccccccctttccctgtaaccccacctaacgttttggacacttaggggcaatttagcgtggccagtctaactaaccgcacatctttggacagtgggaggaaaccagagcacctggaggaaacccatgcagacactgggagaatgtggaaactccacacattcagtgacccag
agtcggtatcaaacctgggtccctggtgctgtggggcagcagtgctcaccactatgccaccgtgccaccctttattgcccatcccttgttgctgattaaaggcataaggcactctttcagcgggtcagtgcagacttaatggaccgaatggcccctttctgcattttagaggtt
ctgtgattttctgattctatgaaagtgcaaactccacacagactgtcacccaagttcggaatcgaacccgggtccctggcgctgtgaggcagcaatgccaaccactgtgccacaatgcttccctgagttctcagattttggccgagcaacccagtgccgaagaagggctgtgta
tccaagcaaggatggtatacgactgagaggggaatttgcaggtggtgatgttcccatgtgtctcactcccatctcctagattgtttagggtgtgggtttagaacttgctgccgaaggaggtttggcaaggtgctgaggtgcatcttgtacatagtgtacactggtgccattggg
tgccaagactgggaggagtgataactaaataagtttctcggcctccactaaacacttacaatggttgtttcttttttgtagttgttttatcaaggaagggaaggacaaaacgttactgcctgaagccagggaaaactctttggagaggtcagaggcgcttctactgtgtggag
tcatggcacagagtgaagaaatgcaagtcctgtaatccaatcagctttatcccaactccgctttaaactgaatcctcaaacctatccaagcaggagcttcaatacggccaattaaacgatccttatgaagagtatcatgattcctaaacattaactccaatatcccacaaaccc
agagagattcttcctggattcagccatggttctgactctgacacacctcttgtcccacccgcagatgtgagccaataatccacgtcgacgatccttgtgcctgtactgtagaaatgccgcactgtcagaggtgccacttttagaatgcaacattaaacagaggaccagtttcac
tgtcagaagatcccatggccactattccaaagaagagctggaatgttctccccggtcttctgaggcaatatttatcccttgggcaacattccttaaaaacgcaggcaaacatacaatctcgtcattgatcccattgctgcttttgggatcatgctgagtcaattggctgctgca
tctcctacattataacagcgccgttgcaatgctctcagtatagcattatgtacatatcggtgaagcaattcttcagcttgttggcaagagatggaaaatcacgatcaaaatacccttgattttccaataagctgcgagaaagttcccatccgaggtgcggagctacaaaattac
tcggtcataattcccatgtccacagctgtaacaatgctttcccagatctgttgatgatcgtacgtttcaatcagcaaggtataatcaagaccactttcatgatcaatatgtatcagaagctttccatctagtagcgctgaaattcattgggtaatcagaaagtcaacatcagtc
acgcattcctcagcttcccctcccccaccaaaacctacgcccccaccccaacctcacctctcaaatccacaccatgttacctgatagacagaacatccaaataattgcaaagctcttcccattaattccccaccagttgcaacccacaaacagagaatggttcccagttgag
atggaggtatggatggaggacggacagcgtggacaagggtgagccatggggactgagccagttcagtctctggtgaaaggaggcaagaagcaaaataattcaagaaaggaagcattggaatgttaggaacagaaggaaatcagttatcccctcttcaccctattaaatggtga
cacaatgaggccatttagccttctgagcttgttgcacaggaacagaaggaaaccattagttccttcaaatccactgcatggggacaggagaaggccaatcagcccgttgagtctgttatttgggaacaaggggctactcaggtattcttgactgttagataggaacaggagaag
gccattcagccccagaacctgttatctatgaacaggaggaggccatttaactcctcgtgtttgcgacatggaaacaggagaaagccattcatccccccttgggcatgtttggtgcaataggagtagaccattcagcctttcaaacttgttctgccattcagttagatcacacctg
atccgtatgtcaaatcccatcgactgtctttaccccatgtcccttgagagccctccccaacacaagtccacagctctcagtattgaaagcttcacattattcagcatctacaacattttgagaattccagatgtccacatccctctgcatttaaaagcacttccagatgtcaca
cacacgcacctcagagttctaattttaagattgtgcccctgtgttctgctttcctccatcagaggaaatagattatttgaaaatatcttagtgaatcctcgcatcggcgtaaatacctcaatccatatcagtgatgggcaacctaggctggtgagtggcagcatgattggccc
tccttcaactcaatggactgcaagatcaaaattggacttgttcactaaccatgacccatgaataagattgaatacatttgatacatgccaattatttcacagcaagttctagaaaatgcttaatcatttatattctgatggaactgtcatcaacttcaaatggtaaaatcaggt
tgcatatttacacttcggacacagaaggagtgcacggttttcacagttaatgttgggctgaattctccgatcccctagccacgtgtttcttggcggcgcactgttcgctggcggcgggatcctctcttcccgcagcttgtcaatgggattttcccattgaagccaccccacact
gctagcaaacccacaggcagggttgcactgccagcaggaaaagagaatcccaaagaacagagaattccgtccgttgtgagcaatcagcacgcacctcacttcacttgcccttgcttaacgtgcgtatcacttcagtcataccggtaggagaaagctacatggatgaaaatcagt
cgaatttggacaacagtcaaaggcggcacggtggcgcagtggttagcactgctgcctcacggcgatgaggacccgggttcgatcccggccccgggtcactgtccgtgtggagtttgcacattctccccgtgtctgcgtgggtcccaccctcacaacccaaaggtgtgcaggctg
ggtggattggccacgctaaattgcccttaattggaaaaaagataataattgggcactctaaatgtatttaaaaaaataatttggacatcagtcaacacaatattttgtaaccacactcagaacccaggatgcgtgtgtctatctgccacatgcatcccctgggcctcagggtac
tcaccactgatctatatccttcctgaggtgtagacccctgaggggtccaaccagggctgtagacaaccctaccccttttccctttgtagcccagaaccctggagtcagggtcaagtctgtgagcaaagaaaatgtcatccacaaagagccaccactgtcagttcacactaatgag
cctttccccaccaacgtgatgcttattcattgaatacgacgcttaagcttcaggttataaatgctgtactaatgtgtttgcacagggagagagcctctgccagcaggacaggatgtggatcttaaacaggatccaggagcagttccaacatacactgctcccaaaactctaact
gatgattggaatgacagaaagtgtcgaactgttgccttcctgttcttaatgaaagggagggccctgcagcttaaaacatggaggcttggattttcattgttattacccaggcatttaacagtgagcgaattggaaaactaggcatgggcagcgtgaaatctcttccattccatt
ttcttttattgatccactcatgggatgtggacatgcctggcgaggtaatttattatcaaaccaaattgctattcagaaggtgatggtgggctttcttcttgaactctgctgtccgtgtggaggaggtgctcccacaatgccattgggttacagggttttgatccagtgacgatg
aagaaatggccaaaaaagttccaagccaggatggtgtgtgacttgcatagaaaataggagcaggagtagtccattcggccctgctagctgcaccgctattcaatatagtcatgggtataaaaccatcgatcccattgaatcaggttaatattaaaattagtgagcatagagagc
aaaatgacctgctttttccaggttgacagtgaggccattactttaaaagcacgttagatcttatattccctccctccactggattctgatggcgaggctctctattgcctggggttccggttgctgagtgcgacctattgtttaacatgtattgcttgtgctgataaaatcaat
gtatccaatgttacaaatttactgcatagctgctatcaattaaaaagttgttcattcattaatcagctccaaattgtatgtatctttattggtatctggagtgttcttgaagaaagcccagtaacgatacagtgcaaatctctgcactgattcctcgacgcaacgtga

**C:** Sequence of cDNA (coding region in red and highlighted in yellow)

gagggagcaggtgcagggctgtcggggatctgcatgactgataatgggagagatataagtcttccaacattcagaagcactgattaaactttcgatcgtgccacttctggatcagtccttaaatctgaagatgcagctggtttccctgattgctgtgtttgccttggccacaat
atctgtcctcacagatgatacaactcagctgaaagaggatgaggcaaagagggtactgacaacaaccagggtgaaatctcctgccagtgaggagagccgctgtgtgtgcaaagttggccacctcgagtttggatacaaagtgaagcagtcccacgatcgaactgggaacaagtg
cccctgtcagaaggccataaacatcatcaagaggaaatggaaagatgttgttttatcaaggaagggaaggacaaaacgttactgcctgaagccagggaaaactctttggagaggtcagaggcgcttctactgtgtggagtcatggcacagagtgaagaaatgcaagtcctgtaa
tccaatcagctttatcccaactccgctttaaactgaatcctcaaacctatccaagcaggagcttcaatacggccaattaaacgatccttatgaagagtatcatgattcctaaacattaactccaatatcccacaaacccagagagattcttcctggattcagccatggttctga
ctctgacacacctcttgtcccacccgcagatgtgagccaataatccacgtcgacgatccttgtgcctgtactgtagaaatgccgcactgtcagaggtgccacttttagaatgcaacattaaacagaggaccagtttcactgtcagaagatcccatggccactattccaaagaag
agctggaatgttctccccggtcttctgaggcaatatttatcccttgggcaacattccttaaaaacgcaggcaaacatacaatctcgtcattgatcccattgctgcttttgggatcatgctgagtcaattggctgctgcatctcctacattataacagcgccgttgcaatgctct
cagtatagcattatgtacatatcggtgaagcaattcttcagcttgttggcaagagatggaaaatcacgatcaaaatacccttgattttccaataagctgcgagaaagttcccatccgaggtgcggagctacaaaattactcggtcataattcccatgtccacagctgtaacaat
gctttcccagatctgttgatgatcgtacgtttcaatcagcaaggtataatcaagaccactttcatgatcaatatgtatcagaagctttccatctagtagcgctgaaattcattgggtaatcagaaagtcaacatcagtcacgcattcctcagcttcccctcccccaccaaaac
ctacgcccccaccccaacctcacctctcaaatccacaccatgttacctgatagacagaacatccaaataattgcaaagctcttcccattaattccccaccagttgcaacccacaaacagagaatggttcccagttgagatggaggtatggatggaggacggacagcgtggaca
agggtgagccatgggggactgagccagttcagtctctggtgaaaggaggcaagaagcaaaataattcaagaaaggaagcattggaatgttaggaacagaaggaaatcagttatcccctcttcaccctattaaatggtgacacaatgaggccatttagccttctgagcttgttgc
acaggaacagaaggaaaccattagttccttcaaatccactgcatggggacaggagaaggccaatcagcccgttgagtctgttatttgggaacaaggggctactcaggtattcttgactgttagataggaacaggagaaggccattcagccccagaacctgttatctatgaacag
gaggaggccatttaactcctcgtgtttgcgacatggaaacaggagaaagccattcatcccccttgggcatgtttggtgcaataggagtagaccattcagcctttcaaacttgttctgccattcagttagatcacacctgatccgtatgtcaaatcccatcgactgtctttaccc
catgtcccttgagagccctccccaacacaagtccacagctctcagtattgaaagcttcacattattcagcatctacaacattttgagaattccagatgtccacatccctctgcatttaaaagcacttccagatgtcacacacacgcacctcagagttctaattttaagattgtg
cccctgtgttctgctttcctccatcagaggaaatagattatttgaaaatatcttagtgaatcctcgcatcggcgtaaatacctcaatccatatcagtgatgggcaacctaggctggtgagtgggcagcatgattggccctccttcaactcaatggactgcaagatcaaaattgg
acttgttcactaaccatgacccatgaataagattgaatacatttgatacatgccaattatttcacagcaagttctagaaaatgcttaatcatttatattctgatggaactgtcatcaacttcaaatggtaaaatcaggttgcatatttacacttcggacacagaaggagtgcac
ggttttcacagttaatgttgggctgaattctccgatcccctagccacgtgtttcttggcggcgcactgttcgctggcggcgggatcctctcttcccgcagcttgtcaatgggattttcccattgaagccaccccacactgctagcaaacccacaggcagggttgcactgccagc
aggaaaagagaatcccaaagaacagagaattccgtccgttgtgagcaatcagcacgcacctcacttcacttgcccttgcttaacgtgcgtatcacttcagtcataccggtaggagaaagctacatggatgaaaatcagtcgaatttggacaacagtcaaaggcggcacggtggc
gcagtggttagcactgctgcctcacggcgatgaggacccgggttcgatcccggccccgggtcactgtccgtgtggagtttgcacattctccccgtgtctgcgtgggtcccaccctcacaacccaaaggtgtgcaggctgggtggattggccacgctaaattgcccttaattgga
aaaaagataataattgggcactctaaatgtatttaaaaaaataatttggacatcagtcaacacaatattttgtaaccacactcagaacccaggatgcgtgtgtctatctgccacatgcatcccctgggcctcagggtactcaccactgatctatatccttcctgaggtgtagac
ccctgaggggtccaaccagggctgtagacaaccctaccccttttccctttgtagcccagaaccctggagtcagggtcaagtctgtgagcaaagaaaatgtcatccacaaagagccaccactgtcagttcacactaatgagcctttccccaccaacgtgatgcttattcattgaat
acgacgcttaagcttcaggttataaatgctgtactaatgtgtttgcacagggagagagcctctgccagcaggacaggatgtggatcttaaacaggatccaggagcagttccaacatacactgctcccaaaactctaactgatgattggaatgacagaaagtgtcgaactgttgc
cttcctgttcttaatgaaagggagggccctgcagcttaaaacatggaggcttggattttcattgttattacccaggcatttaacagtgagcgaattggaaaactaggcatgggcagcgtgaaatctcttccattccattttcttttattgatccactcatgggatgtggacatg
cctggcgaggtaatttattatcaaaccaaattgctattcagaaggtgatggtgggctttcttcttgaactctgctgtccgtgtggaggaggtgctcccacaatgccattgggttacagggttttgatccagtgacgatgaagaaatggccaaaaaagttccaagccaggatggt
gtgtgacttgcatagaaaataggagcaggagtagtccattcggccctgctagctgcaccgctattcaatatagtcatgggtataaaaccatcgatcccattgaatcaggttaatattaaaattagtgagcatagagagcaaaatgacctgcttttttccaggttgacagtgaggc
cattactttaaaagcacgttagatcttatattccctccctccactggattctgatggcgaggctctctattgcctggggttccggttgctgagtgcgacctattgtttaacatgtattgcttgtgctgataaaatcaatgtatccaatgttacaaatttactgcatagctgcta
tcaattaaaaagttgttcattcattaatcagctccaaa

**D:** Amino acid sequence (signal peptide shaded)

MQLVSLIAVFALATISVLTDDTTQLKEDEAKRVLTTTRVKSPASEESRCVCKVGHLEFGYKVKQSHDRTGNKCPCQKAINIIKRKWKDVVLSRKGRTKRYCLKPGKTLWRGQRRFY
CVESWHRVKKCKSCNPISFIPTPL

**Fig. S4.** Gene position and architecture (**A**), genomic DNA sequence (**B**), cDNA sequence (**C**), and encoded protein sequence (**D**) of the unannotated *cxcl17* gene in the NCBI reference genome of *Scyliorhinus torazame* (cloudy catshark). The information was downloaded from the NCBI gene database (https://www.ncbi.nlm.nih.gov/gene).

### A: Gene position and architecture

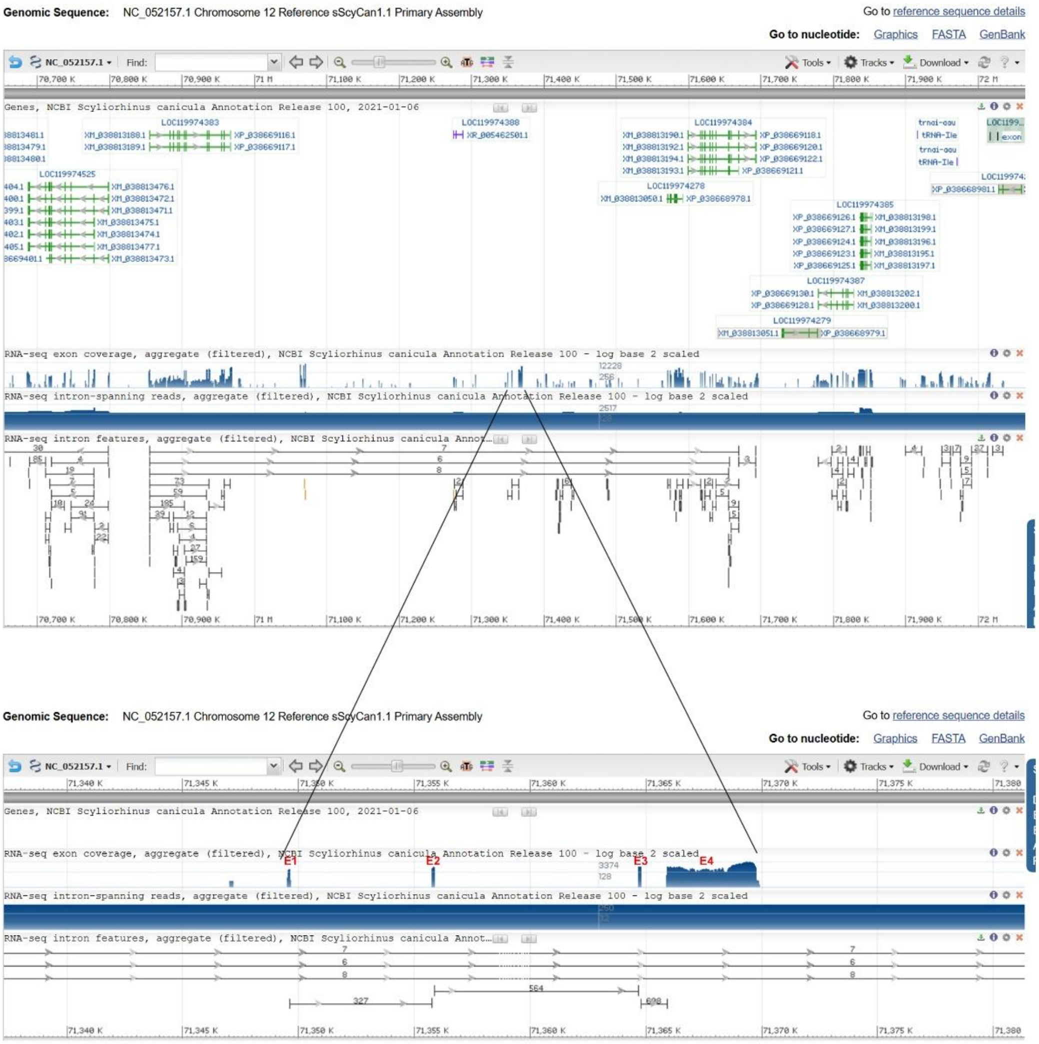


### B: Sequence of genomic DNA (exons in red and highlighted in yellow, coding region underlined)

ggccgggggaggggaaacttgcagcccctcctctgcactggggagggagcaggtgcagggctgtcagggatgactgataataggagcgatataagtcttccaacattcag**aagcgctgattaaactttgatcgcgccacgtctggatcagtcctgaaatctgaagatgcagct**
**tgtttccctgattgttgtgtttgccttggtcacagtatctgtcctcacagatg**gtaagaatttaaatctgtgcaattatttgctttatggtttatcacatgtctattcagatagataattttaaaaatatattatttatatttggaacgagagggttgctggctaaaccagggc
tgaaattctccgcttgttgcgattcaattttcgtgccagcaacgtacatccaccaacggtttcatggcagctcggagtagttacgatgggaaatcccattgacaatcggtgggaattggggaagctgccatagaggcgcgcagttgagaaactggagaaagctgccccaaatta
gttgcaaatttctagttacaatagagaggctttctcagtcgcttctgtttcggtatctaacagcatatattccaccagcaattctgaatgtagttaacaatgcgcagttaaatgaggaataaattccctggctgttcctcttcaaagtaatgtcatgggatccttcatatcaat
ttgatgaggcagatgacttaaggtctcacccgaaagattgctcctgacagtgtagtgccccttctatactgcactggagtgtcagccttaatttccatgcactcaaacattggagtgggatttgaaatcagaatctcggtgttcagaggtgagggagctgtcatctgagctaag
gtgacatgcttgccgcgatactaagaatctatcacacagttgtggaactggagtctcatttcagccagtccgttttaggatggtaggttttgttaactgaaggacatcagtgatcaagttgggtttttacaacaatctgacaatgtcaccgtcactttgactcgtaccagtttt
tttcttttcagacatagctttaaactttatttaaatgctgaacttacaagagtgatatttgaacccaagttacctggggtttaatgtgaacgtgaataattgcattcatttagcgcctttcatgaggtttgcatgtcccaaagtgctttctaattgattaattgcattcagagc
tggagtcactgttgtcctgtaagaaattcaaagctagtttgtacacagcaagctcccagaaacaacaatgtttaagatcagaattgagcagtaaggattgaccacaaaatgggggacaccactgttcttctctgaagagtgactgtgacatttcttttgtttgttgcaaaaata
tactttatttgtaaaatatctgaaagaacatgacaaacatttcaaagtggccatcacagaaagtgcaatcatattcaaattttctacataagtcacgggctggattctctagtcccccctagctgcgagtttctcgatggcgcgctgttcgctgtcggtggaattttcccttccc
gctgcttgtcaatgggatttcccattgaagccagcccatgccgccgggaatcccattggtgggggtgggttgccagcaggaaaagggaatcccaatgactggagtattctggcccatgttgcactctggggtacttcaatacagtcaaagaaacattacagacatttcaaaatg
gtaattacatacagtgcaatggtattttagtttactacataaatcaagttgcattatgaggtgcttcaatacaattctgatacatgtaatattcactgtatacattcagtgtgagtcatacggcccgaggggggttctgtataattccctttccttcagttcactagggctgaaa
gattttaggcaatgaccatccccattgcgcctctacggcagcttccctaagcttcagtgcctccctcagtacgtagtcctgggttttggttgtccaactgagaggttcaaagtctcatctgaacgatactacctcagccagtgcagctcccccctcagtgctgccctgcagctcc
agcctagattatgcgctcaaatctctggagtgggtttcatttagggaagagttgaagaaacaactgttttgttctggaatgcactgcatgagaggacaatgaaagcagattcaatcgcaacttcagggaatcaggtaaatatttaaagggagaaattcttcttctgactgctcc
tcacagcaaggagaaacttgccacacagtatagcccatagcaagaacctgcgcatagagtcatacagttttacaggaagaaacctttggcctatcgtgtctgtgtcggccatcgagcaccaatctattctaaacccattttcctgcacttggtctgtagtcttgtatgctatgg
catttcaagtgctcatctaagtgcttcttaaatgttgtgagggttcctgtctctacaaccctttcaggcagcgagttccaggttcccgccatcctctgggtggaaaagacttttctcaaatcccctcaaaatctcctgccttactttaaatctatgctcctggttatcgacccc
tctactgaggggaaacgtttgttcctgtctatcctctctgtctctcataattgtgatgtggagatgccggctttggactggggtgagcacagtaagaagtcttacaacaccaggttaaagtccaacaggtttgtttcaaacacgagctttcggagcgcagctccttcttcaggt
gacaggagaggtatgatccagaaacatttatatagacaaagtcagcgatgccggacaatgcttggaatgcgagaatttgcaggtaatcaaatcattacagatccagagataggggtaaccccaggttaaagaggtgtgaattgtctcaagccagaacagttggtggaattttgc
aagcccaggtcagatggtggggagtggatgtaatgcgacatgaatccaagatcccggttgaggccgcactcatgcgtgcggaacttaactatacgtttttgctcgcaattctgcgttgtcgcgtgaaagattgctcctgacagtgtagtgcaccctctatactgcactggagtg
tcagccttaatttccatgcgctcaaacattggagtgggatttgaaatcagaatctcggtgttcagaggtgagggagctgccatctgagctaaggtgacatgcttgccgctatactaaaaatctatcacacagttgtggaactggagtctcatttcagccagtccgttctataca
cctagattggttcccctctgcagccttctctgctctaaggagaacaaccccccagcctaaccaacccctcttctcagctgaaactctccagcccaggtaacatcctggtgaatctcctctgcatcctttccagtgcaatcacatctgtcctaaactgtggcaaccagaactgcca
cacagtatctagctgtggcctgacgagcgttttatacagattcatcatagcctgtctgcacttatattctatgcctcggcttattatggcaagtatatcatatgccttctttaaccaccttgtctacctgtcctgctgccttcaggggtctttggatactcaagtccctctggc
cctctgtacatcctagtgtccgaccattcattgtgtgttctcttgctttgttagtcctaccgaaaagcatcacgtcatacatttcagggttaaattccatttgctaggcttgatttaatcgaatttgaaatagtttatagggcgcacctgagtaagtccagaatgagtcgcttc
tttgcatgggtggaagacacgtgcgtgctctgctcaaggcgtctggccaaccaggttctgctctttccccaagcttgtgggttctctaggtctccttcttcagcaccatgtcggcgattctgaaaattcaactggagccctgtcaacttggtggccaaattttgagtgtcagtc
gtgctcgagctgcagacaagtgtgggggccgataccttggccttcgcctcgctggtttcctggaggtgggtactgctggggtggaggtctgcttctccgcccagtacctcagtgtggctgggagatctgatggagtttctatacctcgcgaaagtcaagtacaccgcgcgggga
gcaattgaggggttttactgaagatgacagccgtctgttgtttatttcaaggaattggccatggttcattgctcagaaggggagaggggaggaagaagagaaggggggaggggttaggttttgtgaggggtgggctatgttttggaagctttatgtttgtattagtgtattattg
tgttgtgttgttttgtattatgtataaaatgaaaaataaaaatatatttttaaaaattccattgccatggttctgcacatgtgaccaacctgactatatcatcctgtaatcacagagtctgttgtgtggcggctaccacattgttcagtccagccaggagactatcccactctt
actctctgccttagacttattggacagattttcaggttgacctccttctctcctccatgatagttaaagacttggttagttatagcaatgagcagttggatccatatgaacaagcacctcttccctggaaccttatctgaactgaaattgttctgttccgttggcagtgcacca
caatcagatagatgctgtacaagtatgagctgcgggtactgttggactcggcgattaagtggatgtgatctatgagtcactgctgaggttctcaaattctcgaaacattttataaactcaaacatggccccccttacaatttgttgtttggtctgtcactgctctatagcaaatc
tctcctataattagctttgaatctccaactccaagatgcatcaagggacaaagcctagaatcttgccacctctgtggtttaattgtatcttagcttgcaacaagaaatccccagggtcatcctaggggcagggcaagacgacgttccagaaaatcccaggaattacaagtcaga
aaataagaattcaggaattggtccatcaaaataaaaggtccatttctaccatcttcagcatgtaaatgttgcaattaatctctgtcgatcaatctttaactggaggtgaagcaaagcccaagtggtacagagttttgagaacagcagctcttgctcctcaaaaacatgttccac
tcaccacagttttgttactcaaatgggggcgtcgctgactgagtgacttacaaggccatttcagaggatagttatgagtcacatgtaggccagaccagctaatgatgacaggtttgcttccctgaagtgaacgagatgggtttttacaacaatcatttcatgataacttattca
ttccagattgttttctactgaattaaattttgactgactgctaaactgggatttggacctgggtccccaatttcctaggtcactggattgcaggttcggtgacaataccactacgctaatgcctccttggttaatgtcaggatagtctcggccagtgaccgatttccctaagta
gtcctgggttatggggcaggaatcagtcagttttctttttttcatgttctctgaacaaccccaacaacaacttccactcattttacagagcctttaacagaagaaaaggtcccaccaaactacttcagaggagcgttatatcaataaacaccatgtcagacaggatgatgttgga
cgcatcaccaaatgtttagcgaaagtggtagattttcaaggaggtggagagtaaagtagagaagacgataggtttacagggagggaacaaggattagcttttaaaatccaattgtttcaaaactaagaatcagtgtaaatcggcaatgactggaacattacaatcaaaagaaaa
aattgtcagggcgacacggtggtaaaaggttagcactgttacctcatagcaccaagaacccgggttcaatcccggctctgggtcactgtccgagtggcgattgcacattctctccatgtctgcgttattctcaccccccacaatccaaagacgtgcagggtaggtggatttgcc
acgctaaattgctccttacttagaaataaaaagaattgggcactttaatctttttttctaaactatcagtatcattgaacaaagtgaatgtctgtcccagcctctattcagtcagtgacatcacgtcaaatggactggacaggtctgtcaagcatgttttttagggatcaccacg
atgacaagaaaaggtcacaaaaattgctttgattgtgaatataaattaccagggtttatatttttacaggcatgtttattatttccag**atacaactcagctgaaagagggtgaggccttgagggaactgacacaaccagggtgaaatctcctgccagtgagagccactgtgtc**
**tgcaaag**gtaggaaggcacattcagacctcgattactcaccaacctgatcaacaagataatgaaatcagagactgggatgatcaaacccaaaactgggttaaccatggggcagataaaccagggacacggtgaatagggaaacagttcaaataaaaggactagataaagatgg

gaatagttaatccagagattggataaccacgggaacagttaatccagggactggataaacacataaatagttaatccatggactggataaacatgggaatagttaatctgtagactggataaagacaggaatagttaatccatggactggataaacatgggaatagttaatctg
tagactggataaagacaggaatagttaatccatggactggataaacatgggaatagttaatctgtagactggataaaggcaggaatagttaatctgtagactggataaacacgagaatagttaatccatagactggataaacatgggaatagttaatccataggctggataaac
atgggaatagttaatccataggctggataaacatgggtatagttaatccatagactggataaacatgggaatagttcatccatagactggataaacacgggaatagttcatccagagactggataaacatggaattggttaatccagggactgggtaatcatagaaatcattaa
ttcagggctggataagcatgggaacagttaatccatagactcgataaacacaggaatagttaatccagagtctagataaacacgggaatagttaatccagagttaaatgaacatgggaatagttaatccaaaaactggaataaatttgggaatagtaatccaaaaactggatc
aacgctttgtagttaatccagggactggataaacatgccagctgttagtccagggactgggtgaatacaggaatagctaatccagagatttgataaacacggtggtgttgtctgtaagattcagaacttcaaccagtttattattcaagcgttttaaccagatgtccatatttg
cgagataagcaaaggactaacacaagataacaattcaaccaaagtttattttcaagcattataaaatagtttatccccaaaattatcctaactatagtatgccaaagggtcttaattctacacccgccgatgaacttgatcagctggtcgaatccagtgagtctcagtcttct
cagggcctttgtctgcgaaggtgagtctcacatgttgcttacttctgtcctctggtctgccgctgagtcgtctctgctgcctctgcatcgtgtcttcactcaggtgatctctagatatctcttgctcatcctcttcctcacacccttccagacgtttctccgcatcttcagggg
ctaggtcggtgccagagtggtttgcgctgcgtcagccagcgtggaaggggcttggcgccacaccaaccggcgccgaagggcctctgccagctggcgtgagttggcgcatgcatgggaacgccagcgtgtgctggcgtcatcctagcgcatgcacagaggggggttcatctccgcg
ctggccatcgcagaccggtacacaagccgggcgcggaggaatagagtgcccccacggcacaggcctgcacgcagatcggtgggctccaatcgtgggccaggccaccttgggaacagcacccagggccagatccccccacgccccccccgaggacccagaggccgcccgcggagc
caggtcctgccagtaagtacctgttgtaacatacgccggcgggaccggccgaaaacaggtggccactcggcccatcgcaggctggagaagcgctggggtagccgttggcagcagccgccgaccggcacggtgcgattcccacaccgccaaatccccggcgccagagaatgtgg
cagccggtgggtgcgggattcccgcccaccccccagtgattctccaacccggttggggggggggggtcggagaattccgcccctagtttattgcccaatcactgttggatgcctgtaccctcgtccacccacttgaatcctggccatcacagagggctattgctctcttcattcag
gctcaagctggatctggattatacaagattagtgggtttgaatttggagctactaggtaacttctttgaagaccactcagaggagcaccaagtgtctgtaaaattcagacatatcgaccagtttattattcaaatgtttgacccaggtgcccttatttctgaggatgagcaaa
ggaccgacacaggttaacaattcaacccaagtttattttcaaacacattggccgggattctccgtcccagtgccgggttggagaatccccgagggagcgcgagaatcgcgcccggcagccccgacgccagctccccattctcagacgctgattttcgggcacccgaaggatcg
ccgttgacagcagcccccccccagcgattctccgggccccgatgggccgagtggccgacgagtttggccaagtcccgccggcgtgtgttattcaggtcccacatggcgtgatctggaaggtgtgtctgcaggggccgtcctggaggggggggtgggggatccgatccgggggg
gatccgacccgggggggggcgggggcacggtggccatgcccgcgatcggggcctaccgattggcgggcggctggttccgtgcgggcctattttactctgcaccaggcccctgtaggctccgccatacttcccgggggccggtgtggagaagggaaccccgcgcatgcgcgt
aaaaacgccggctgttccgcgcatgcgcaaatttgcgcggcccattccgcggcagctggagctgcccactaggaagggagaatttcctcacctttcaggctgttgatgccggagtggttggcaccggttttcacgccagcgtggggcatagccccattattagagaatcccgc
ccaatgaaacagtttgtccccaaattatcccaaatagaatctggctaagatccttaatactgcccgtgccgatgaactggattgagttgtgggtcgaatcctcgagtttagattgtctcagggcctttgactgcgaaggtgggtgtcgcatgctgcttccttctgccctctg
gtccaccattgagacttctccgctggctctgcattgagtctttgctggggaggatctctagatgcctccttcttacccccacttcacacttcctgaagcttctctggccctcaaccaataaggtcacggtggtggttgcaacacccaatggagttgtcgattgcaactctg
caggactcctggacctctcctcagaggtccattctcaggtgcattatctgcatatgaccttatcccatatcaggtgatctaggtgccagaacatccagcagagggcagcagagcagagctactgatcagctgttctggggaaatttgcatacgtgcagtgcggtcagcctaagt
tgaaggtgaatctgcgaagaagacccgaggagtttgagtgaaggcaatcatccagcagagggcagcagagcagagctactgatcagctgttctggggaaatttgcatacgtgcagtgcggtcagcctaagttgaaggtgaatctgcgaagaagacccgaggagtttgagtgaag
gcaatcatccagcagagggcagcagagcagagctactgatcagctgttctggggaaatttgcatacgtgcagtgcggtcagcctaagttgaaggtgaatctgcgaagaagacccgaggagtttgagtgaaggcaatcatccagcagagggcagcagagcagagctactgatcag
ctgttctggggaaatttgcatacgtgcagtgcggtcagcctaagttgaaggtggtttgtggagaggctgttggcaagtgacagttaaacctgaagcactttgtgagtgtttcccaccctacctcctcctctaaccaacccccccacccacagtggttggaagcgggagcagg
ggcctgtcgtgaaggtgagtgagtgcctttaaatttgcttacctttcagcgggagcagggtttgaagtaaatatcaggtaagctcttccttctttttctttttcttgtttttttttaaatctagaggtgatgtcagggaaggcagtacaatgctcctcctgcagaatgtttgag
gtgagggacgccgtcagcgtccctgctgatttcatctgtgggaagtgcacccaactccagctcctcaaaaaccgtgttagggacctggagcttgagctggatgaacttcggatcattcgggaggcagagggggtcatagataggagcttcagggaagtagttacaccaaagact
ggagatagatgggtaactgtaagagggactgggaagaagcagtcagtgcagggacccctgcggtcgttcccctgagtaacaagtatacgttttgatacttgtgggggggacgacttaccaggggtaagcctcatcggggtacaggcctctggcacggagtctgtccctgttgct
cagaagggaagggggaaaggagtagaacattagtaattggggactcaatagtcaggggcatagataggagattttgcgggagcgagagagactcacgtttggtatgttgcctcccaggtgcaagggtacgtgatgtctcggatcgtgttttccgggtccttaagggggagggg
gagcagccccaagtcgtagtccacattggcactaacgacataggtaggaaagggacaaggatgtcaggcaggccctttagggagctaggatggaagctcagagcgagaacaaacagagttgttatctctgggttgttgcccgtgccacgtgatagtgagatgaggaataggag
agagagcaattaaacacgtggctacagggatggtgcaggcgggagggattcagatttctggatgactggggctctttctggggaaggtgggacctctatagacaggatggtctacatctgaacctgaggggcaccaatatcctgggggggagatttgttagtgctctttggggg
ggtttaaactaattcagcaggggcatgggaacctggattgtagtttggggtacgggagattgagagtatagaggtcaggagcacagatttgacttcgcaggagggtgccagtgttcaggtaggtggtttgaagtgtgtctacttcaatgcgaggatacgaaataaggtag
gggaactggcagcatgggttggtacctgggacttcgatgttgtggccatttcagagacatggatagagcagggacaggaatggttgtgcaggttccggggtttaggtgtttttagtaagctcagagagggggcaaaagagggggaggtgtggcgctgctagtcaaggacagta
ttacggtggtggaaaggatgatagatggggactcttcttctgaggtagtatgggctgaggttagaaacaggaaaggagaggtcaccctgttgggagttttctataggccacctaatagttctaggatgtagaggaaaggatggcgaagatgattctggaaaagagcgaaagta
acagggtagttgttatgggagactttaactttccaaatattgactggaaaagatatagttcgagtacattagatgggtcattctttgtacaatgtgtgcaggagggtttcctgacacaatatgttgacaggccaacaagaggcgaggccacattggatttggttttgggtaatg
aaccaggccaggtgttagatctggaggtaggtgagcactttggaaacagtgaccacaatcggtgaccttacattagtgatggaaaggataagtatacccgcagggcaagagttatagctggggaaggcaattatgatgccattagacatgacttaggatgtgttggtt
ggagaagtaggctgcaagggttgggcacactggatatggagcttgttcaaggaacagctattcatgttcttgataagtacgtaccagtcaggcagggaggaagggtcgagcgaggaaccgtggtttaccaaagaagtggaatctcttgtaaggaagaagaggcct
atgtgaagatgaggcgtgaagtttcagttggggcgcttgatagttacaaggaagcgaggaaggatctaaagagagagctgagacgagcaaggaggggacatgagaagtctttggcaggtaggatcaaggaaaacccaaaagctttctataggtatgtcaggaataaaagaatga
ctagggtaagagtagggccagtcaaggacagtggtgggaagttgtgtgtggagggctgaggagataagcgagatactaaatgaatacttttcgtcagtattcactcaagaaaaagataatattgtggaggagaatgctgagacccaggctattagaatagatggcattgaggtgc
gtagggaagaagtgttggcaattctggacaaggtgaaaatagataagtccccggggccagatgggatttatcctaggattctctgggaagccagggaagagattgctgagcctttggctttgatttttaggtcatcattgtacagaatggtgccagaggactggaggatag
caaatgtggtcccttttgttcaagaaggggagtagagataaccccggtaactatcaggccggtgagcctaacgtctgtggtgggtaaagtcttggagaggattataaaagatacaatttataatcatctagataggaataatatgattaggacagtcagcatggttttgtgaagg
gtaagtcatgcctcacaaacctatcgagttctttgagaaggtgactgaacaggtagacgagggtagagcagttgatgtggtgtatatggatttcagtaaagcgtttgataaggttccccacggtcgtctattgcagaaaatacggagctggggattgagggtgatttagaga
tgtggatcagaaattggctagttgaaagaagacagagagtggtagttgatgggaaatgttcagaatggagttcagttacgagtggcgtaccacaaggatctgttctggggccgttgctgtttgtcattttttataaatgacctagaggagggcgcagaaggatgggtgagtaaat
ttgcagacgacactaaagtcggtggagttgtagacagtgcggaaggatgttgcaggttacagagggacatagataagctgcagagctgggctgagaggtgcaaatggagtttaacgtggagaagtgtgaggtgattcacttttggaaagaataacagaaatgcgaatatttg
ctaatggtaaaattcttggtagtggatgagcagagggatctcggtgtccatgtacatagatccctgaaagttgccaccaggttgataggttgtgaagaaggccctatggtgtgttggcctttattggtagagggattgagttccggagccatgaggtcatgttgcagttgt
acaaaactctagtacggccgcatttggagtattgcgtacagttctggtcgcctcattataggaaggacgtggaagctttggaacgggtgcagaggagatttaccaggatgttgcctggtatggagggaaaatcttatgaggaaaggctgatggacttgaggttgttttcgttag
agagaagaaggttaagaggtgacttaatagaggcatacaaaatgatcagagggttagatagggtggacagcgagagccttctcccgcggatggaggtggctagcacgaggggacatagccttaaattgaggggtaatagatataggacagaggtcagaggtgggtttttacgc
aaagagtggtgaggccgtggaatgccctacctgcaacagtagtgaactcgccaacattgagggcattaaaaatttattggaagcatatggatgataagggcatagtgtaggttagatggcctttagatttttccatgtcggtgcaacatcgagggcgaaggccgtac
tgcgctgtatcgttctatgttctatgttctatctggagtcatctgggagatccacaacaattgatccatttgaatgtggctaattatcttccaatgtcctgtttacaggacagctcagacttactccgactgtctctctctttccagtgtatacaagtttgactatttgactt
cccatcaggcccgtctgtgattctggctttaatgtaaagtgtccaattcggtatcaggcctaaaataagggacattgaccattttaccacaacgtgaagagttaatccagggactggataaacacaagaataattactgcaggaacacaataaacacaggaagagttattctgg
ggattagataactaagggaatagttaattcagggacttgataaacatgggaatagttaatccagggactcaataaacatgggaatagttaatccagagactgtatgaattcaagaatagttaaaacaggagctatttaaacatgggatagttaatccatgaactggatcaacac
gggaatggttaattcagggattcgatgaacatgggtatacatcatccagggaactgataaatatgagaatagttaagccagagatggggcaaacacaggaatagttaatccagagatagtataaattcaagaatagtccagagccagatagatatgggaatagttaagacaggg
acagaattaacatgggaacagttaaaccagtgactgggaaatatgttgaactgatcactgatcactgattctgtgttcactatttaacgttggtctgaattacacccaaactgactacaagtttttatcccattcttagttggccacctcgagtttcgacacaaagtgaagaagt
cccacgatcgaactgggaacaagtgcccctgtcagaaggctaaaaagatcatcaagaggaaatggaaagatggtaatttatgaaagcataaagtaggggaagaagtcagtcaattgaaacataatgtgtggttggtatagcatgcttgagaggtgcagatgactgcggacctatg
gttcccagagccctccatcactggacaatgtcccccttttgctcatcatagaatccttacagtgcagagagaggccacatagcccatcgagtctccaccgacctttcgaaaaaccactgtacctaggcccaatcccaggtatatgggatcttgccgtagaatcccgacagtgca
gaaggaggtcattcggcctatcgagtctgctctgttcctgcgaaaaaccaccctacctgtgctcaatcacgccgcccttcccccgaaccctacctcacctaacgtttttggacacttacgggaatttagcgtggccagtccaactaacctgcatgtcttcagatagtgggaggaaaccg
gcacatctgggaaatccacgcagtcactcggagaatgtgcaaatttcacattcagtgacccagagtcggtatcaaacctgggtccctggtgctgtgggatagcagtgctaaccactgtgtcaccgtgcacccttattgcccatgccttgttgctgattaaaggtatatgg
cactctttcagagggtcagtgcagacttaatggaccgaatggcccctttctgcattatagaggttctgtgatttttatgattctatgaaagtgcaaactccacacagactgtcacccaagatcagaatcgaactcgggtccctggcactatgccaaccattgccaccatgctgc
cctgagttctccgattttgaccaaacaacccagtgccgaagaagggatgtgtatccaagcgaggatggtatgccactgagacgggaatttgcaggtggtgatttcccatgtgtctagctcccatctcctagattgtatagggtctgggtttagaacttgctgctgaaggagatt
tggcaaggtgctgaggtatcttgtacatagtatacgctggtgccattgggtgccagtactgggaggagtgataactaaataagtttctcggcctttactaaacactgacaatggttgtttctttatttgtagctgttttatcaaggaaaggagggacaaaacgttactgcctga
agcctgggaaaactctgtggagaggccagaggcgcttctattgtgtggagtcatggcacagaatgaagaaatgcaagtcctgtaatccaatcgactttatcccaactccgctgtaaactgaatcctcaaacctatccaagcagggagcttcaatacggccaattaaacaatccc
tatcaagagtatcatgattccgaaacattaactccaacatcccacaaaccagagagattccttcctggattcagccatggttctgactctgacacacctcttgtcccacccgcagatgtgagccagtaatccacgtcgacgatcctcgtgcctgtaccgtagaaatgctgcac
tctcagaggtgtgacttttagaatgcaacactaaacagaggcccaatttcactggcagaagatcccatgcccactattccaaagaagagctgggatgttctccctggtcctctgtggcaatatttatccctttgagcaacattccttaaaaacgtaggcaaacacacaatctcgt
cattgatcccattgctgcttttgggatcatgctgtgtcaattggctgctgcatctcctacattataacagcgtcgttgcaatgctctcagtgtagcattatgtatatatcagtggcaagagatggaaaatcacaatcaaaatacccttgattttccaataatctacgagaaagt
tcccatccgaggtgcagagctacaaaattactcggtcataattcccattacggctgtaataatgctttcccagatctgttgatgatggtacgtttcaatcagcaagggtataatcaaggccactttcacgatcaatacgtatcagaagctttccatctgatagcggtgaaattc
attgggtaatcaaaaagtcaacatcagtcactccttcctcagcttcccctcccccaccaaaacctaactccacccccccaaactcacctctcaaatccacaccatattacctgattgatagaacatccaaataattgcaaagtcttcccattaattcccaccagttgcaac
ccacaaacagttgagatggaggtatggagtgggacagacaacatgagccatggggactgagcctgttcagtctctggcgaaaggagacaagaagcaaaataattaagaagcattggaatgttatgaacattatgtcaccctattaaatggtgacataatgaggccatttggcct
tctgagcttgttgcacaggaacagaaggaaaccattggttccttcaagtctactgcatggggataggagaaggccaatcagcccgttgagtctgttatctgggaacaagaggctactcaggccttctttactgttagataggaatagaggaacgccaatcagccctggaacctt
ttatctatgaacaggagaaagccattcatccccctgggcatgtttgtggaataggagtagaccattcagcctttcaaacttgttctgccactcagttagatcacacctcgtcgtacgtcaaaacccattgactgtctttacccatgtcccttgagagccctccccgacaca
agtccaccgctctcagtattgaaagcttcacattattcagcatctacaacatttggggaattccagatttcacttccctcgcatttaaaagcacatcagatatcacacacgcacctcagagctctaatttaagattgtgcccctgccttctggtttcctccatgaaggga
aatagattatttgaaactatcttaccgaatcccagcatcggtgtaaatacctcaatttatatcagtgatgggcaacctaggctggtgagtgagcagcatgattggccctccttcatctcggtggactgcaagatcgaaatcggacttcttcacgaaccatgacccgtgaataag
attgaatacatttgatacatgctgattatttcatagcaagttctagaaaatgcttaatcgtttatatttctgatggacatcatctttaaatggtaaaatcagattggatatttacactttggacacagagtgagtgcacggttttcacagttaatgtcgggctgaattctctggt
ccccagccatgtgtttctcggcagaactgctcgctggtggcgggatcctctcttcccgcagcttgtcaatgggatttccattgaagccatcccacactgccagcaaaccacacaggcagggttgcattgccagcagaagagagaatcccaaagaacagagaatccggc
cattgtgagtaatcagcacacacctcacttcacttgcccttgcttaacgtgcgtatcacttcagtcatacggtaggaaaaagcgcagcatggatgaaaatcagtcaaatttggacaacagtcaaaggtggcatggtggcgcagcggttagcactgcctgcctcacggcgctcagga
cccgggttcgaccccagtcccgggttactgtccatgtggagtttgcacattctcccatcgtctgcgtgcgtcccaccccccacaactcaaagatgtgcaggttgggtgcattggccacgctaaattgcccttaattggaaaaaagataataattgggcactctaaatttattta
aaaagaatttggacaacagtcaacacaatattctgtagccacactcagaatccaggatgcgtgtgtctatctgctacatgcatcacctgggcctcagggtactcagcactgatctgtatccttcctgaggtgtagaccccctgagggggtccaaccagggctgtagacaaccct
acccctttcccttttgtagcccagaaccctggagtcacggtcaagtctgtgagcaaaggaaatgtcatccacaaagagccaccactgattcacactaatgagctttcccccaccaatcggatggtgcttatacattgaatacgacgcttaagcttcaggttataaatgctgtact
aatgtgtttgcacagggagagagcctctgccagtaggacaggatgtggatcttcaacaggatccaggagcagttccaacatacactgctcccaaaactctaaatgagcgttggaatgacagacagtgcgaactgttgccttcctgttcttaatgaaagggacggccctgcagc
ttaaaacatggaggcttagattctcattgttattacccaggcattgaacagtgagcgaatttggaaaactaggcatgggtagcgtgaaatctcttccattccattttcttttattgatccactcatgggatgtggacatgcctggcaaggtaatttattatccattccaaattgc
cattcagaaggtgatggtgggcttccttcttgaactctgctgtccctgtggaggaggtgctcccatatatgccattggattacagggttttgatccagtggcaatgaagaaatggccaaaagagttccaagccagggtggtgtgtgacttgcatagaaaataggagcaggagtag
tccatttggccctgcgagccgcaccgcattcaatataatcatgggtataaaccatcatgcccattgagtcaggttaatattaaaattagtgagtatagggaggagaaattacctgctttttccaggttgacagtgaggccattactttaaaagcaagttagatcttatattcc
ctccactggtatctgatggcgaggttctctctcgcctgggttccggttgctgagtgcgacctactgtttaacatgtactgcttgtgctgataaaatcaatgtatccaatgttacaaatttactgcatagctgcgatcaattaaaaagttgttcattcattaagcagctccaaat
tgtctgtatctttattggtatctggagtgttcttgaagagagcccagtgacgatacagtgcgaatctctgca

### C: Sequence of cDNA (coding region in red and highlighted in yellow)

aagcgctgattaaactttgatcgcgccacgtctggatcagtcctgaaatctgaagatgcagcttgtttccctgattgttgtgtttgccttggtcacagtatctgtcctcacagatgatacaactcagctgaaagagggtgaggccttgagggaactgacaacaacccaggtga
aatctcctgccagtgagagccactgtgtctgcaaagttggccacctcgagtttcgacacaaagtgaagaagtcccacgatcgaactgggaacaagtgcccctgtcagaaggctaaaaagatcatcaagaggaaatggaaagatgctgttttatcaaggaaaggagggacaaaac
gttactgcctgaagcctgggaaaactctgtggagaggccagaggcgcttctattgtgtggagtcatggcacagaatgaagaaatgcaagtcctgtaatccaatcgactttatcccaactccgctgtaaactgaatcctcaaacctatccaagcagggagcttcaatacggccaa
ttaaacaatccctatcaagagtatcatgattccgaaacattaactccaacatcccacaaaccagagagattccttcctggattcagccatggttctgactctgacacacctcttgtcccacccgcagatgtgagccagtaatccacgtcgacgatcctcgtgcctgtaccgtagca
gaaatgctgcactctcagaggtgtgacttttagaatgcaacactaaacagaggcccaatttcactggcagaagatcccatgcccactattccaaagaagagctgggatgttctccctggtcctctgtggcaatatttatccctttgagcaacattccttaaaaacgtaggcaaac
acacaatctcgtcattgatcccattgctgcttttgggatcatgctgtgtcaattggctgctgcatctcctacattataacagcgtcgttgcaatgctctcagtgtagcattatgtatatatcagtggcaagagatggaaaatcacaatcaaaatacccttgattttccaataat
ctacgagaaagttcccatccgaggtgcagagctacaaaattactcggtcataattcccattacggctgtaataatgctttcccagatctgttgatgatggtacgtttcaatcagcaagggtataatcaaggccactttcacgatcaatacgtatcagaagctttccatctgata
gcggtgaaattcattgggtaatcaaaaagtcaacatcagtcactccttcctcagcttcccctcccccaccaaaacctaactccacccccccaaactcacctctcaaatccacaccatattacctgattgatagaacatccaaataattgcaaagtcttcccattaattcccc
accagttgcaacccacaaacagttgagatggaggtatggagtgggacagacaacatgagccatggggactgagcctgttcagtctctggcgaaaggagacaagaagcaaaataattaagaagcattggaatgttatgaacattatgtcaccctattaaatggtgacataatgag
gccatttggccttctgagcttgttgcacaggaacagaaggaaaccattggttccttcaagtctactgcatggggataggagaaggccaatcagcccgttgagtctgttatctgggaacaagaggctactcaggccttctttactgttagataggaatagaggaacgccaatcag

ccctggaaccttttatctatgaacaggagaaagccattcatccccttgggcatgtttgtggaataggagtagaccattcagcctttcaaacttgttctgccactcagttagatcacacctgatccgtacgtcaaaacccattgactgtctttaccccatgtcccttgagagcc
ctccccgacacaagtccaccgctctcagtattgaaagcttcacattattcagcatctacaacattttgggaattccagatttccacttccctctgcatttaaaagcacatccagatatcacacacgcacctcagagctctaattttaagattgtgcccctgccttctggtttcc
tccatcagaggaaatagattatttgaaactatcttaccgaatcccagcatcggtgtaaatacctcaatttatatcagtgatgggcaacctaggctggtgagtgagcagcatgattggccctccttcatctcggtggactgcaagatcgaaatcggacttcttcacgaaccatga
cccgtgaataagattgaatacatttgatacatgctgattatttcatagcaagttctagaaaatgcttaatcgtttatattctgatggacatcatctttaaatggtaaaatcagattggatatttacactttggacacagagtgagtgcacggttttcacagttaatgtcgggct
gaattctctggtcccccagccatgtgtttctcggcagcgaactgctcgctggtggcgggatcctctcttcccgcagcttgtcaatgggattttcccattgaagccatcccacactgccagcaaacccacaggcagggttgcattgccagcaggaagagagaatcccaaagaaca
gagaattccggccattgtgagtaatcagcacacacctcacttcacttgcccttgcttaacgtgcgtatcacttcagtcataccggtaggaaaaagcgacatggatgaaaatcagtcaaatttggacaacagtcaaaggtggcatggtggcgcagcggttagcactgctgcctca
cggcgctcaggacccgggttcgaccccagtcccgggttactgtccatgtggagtttgcacattctcccatcgtctgcgtgcgtcccacccccacaactcaaagatgtgcaggttgggtgcattggccacgctaaattgcccttaattggaaaaaagataataattgggcactct
aaatttattttaaaaaagaatttggacaacagtcaacacaatattctgtagccacactcagaatccaggatgcgtgtgtctatctgctacatgcatcacctgggcctcagggtactcagcactgatctgtatccttcctgaggtgtagacccctgagggggtccaaccagggct
gtagacaaccctacccctttccctttgtagcccagaaccctggagtcacggtcaagtctgtgagcaaaggaaatgtcatccacaaagagccaccactatcagttcacactaatgagcctttccccaccaatgtgatgcttatacattgaatacgacgcttaagcttcaggttat
aaatgctgtactaatgtgtttgcacagggagagagcctctgccagtaggacaggatgtggatcttcaacaggatccaggagcagttccaacatacactgctcccaaaactctaaatgagcgttggaatgacagacagtgtcgaactgttgccttcctgttcttaatgaaaggga
cggccctgcagcttaaaacatggaggcttagattctcattgttattacccaggcattgaacagtgagcgaattggaaaactaggcatgggtagcgtgaaatctcttccattccattttcttttattgatccactcatgggatgtggacatgcctggcaaggtaatttattatcc
attccaaattgccattcagaaggtgatggtgggcttccttcttgaactctgctgtccctgtggaggaggtgctcccataatgccattggattacagggttttgatccagtggcaatgaagaaatggccaaaagagttccaagccagggtggtgtgtgacttgcatagaaaatag
gagcaggagtagtccatttggccctgcgagccgcaccgctattcaatataatcatgggtataaaaccatcgatcccattgagtcaggttaatattaaaattagtgagtatagggagggaaattacctgctttttccaggttgacagtgaggccattactttaaaagcaagttag
atcttatattccctccactggtatctgatggcgaggttctctctcgcctgggttccggttgctgagtgcgacctactgtttaacatgtactgcttgtgctgataaaatcaatgtatccaatgttacaaatttactgcatagctgcgatcaattaaaaagttgttcattcattaa
gcagctccaaattgt

**D:** Amino acid sequence (signal peptide shaded)

MQLVSLIVVFALVTVSVLTDDTTQLKEGEALRELTTTRVKSPASESHCVCKVGHLEFRHKVKKSHDRTGNKCPCQKAKKIIKRKWKDAVLSRKGGTKRYCLKPGKTLWRGQRRFYC
VESWHRMKKCKSCNPIDFIPTPL

**Fig. S5.** Gene position and architecture (**A**), genomic DNA sequence (**B**), cDNA sequence (**C**), and encoded protein sequence (**D**) of the unannotated *cxcl17* gene in the NCBI reference genome of *Scyliorhinus canicular* (smaller spotted catshark). The information was downloaded from the NCBI gene database (https://www.ncbi.nlm.nih.gov/gene).

**A:** Gene position and architecture

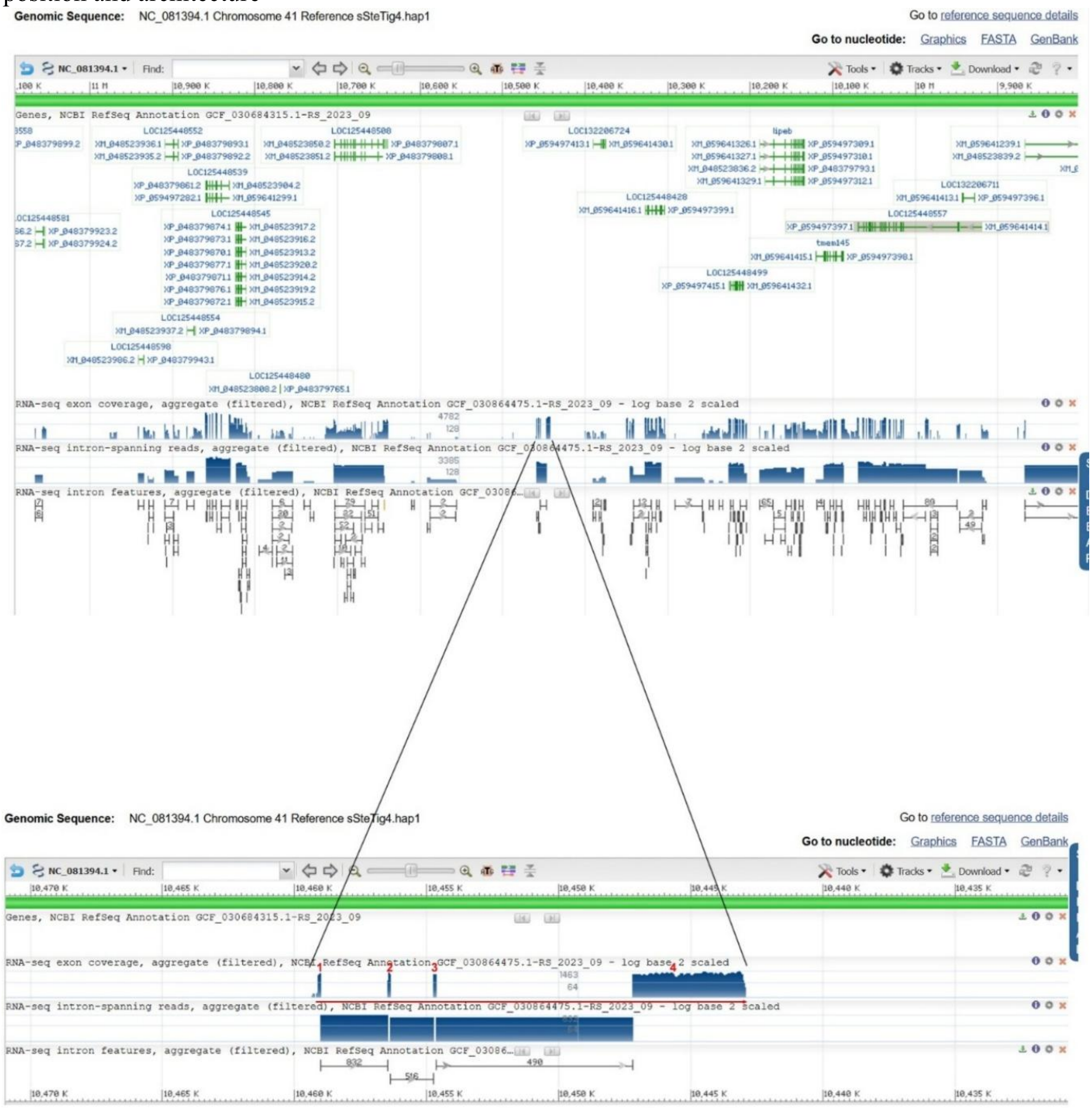


**B:** Sequence of genomic DNA (exons in red and highlighted in yellow, coding region underlined)

tctctggagggcaggggatgggtgggaggtctgtcccagtgcagagctgtcagggataagcatgattgatttcaggagagatataaacttccctgtgctcagaggctctgattaaacagctcgggaccagtgaacacccgaagatgcagatcacactcctgcttattgcctttg
cctttgcatcgatccctgtgctcaccgacggtaagactggattgcaacagtttgctttagagagaatgtaaagctcgttgttcattcgagagtgcagtacttacaatcattctcaaaggaacgggtggtgtggggaggcagggtgctgggggtcattggtaatgtcagcgataa
gtcctgacattttactcaaggatgtgaccctcccatgcagctccggggcaggtaagggcttaggcctagggatcatagatgctcagtggtaggcacactaccattccaccacacacgcccgcagcggaggcatgtctttgagactcaaacacgcaacggcgaccttgggttcaa
catttggccagctcagcagatcaatggcaggttcggttccctgaaggacgttgatgcttccaattggatatttcgccagccttcgcccgaatttgcaaacgagcaggcttccgtgggagcagccaccccacgacctgggcaggcgcagctgcattggcgtagcgcctcatgccg
ggctcctttcgagggaccccagagccacaggaggccaagctttactgggacgatcagccgcacctcctgaggtgcccgtttcttggggtcgacatctgaattgtcgacaaatattccagcccgacgtggccgctccagaatccggcatgagctgcgattatcggtgaagggctc
ctgaaccgtggcttgtacagagaatgcaagctgcagtaatgggttcatctgtggtttcaacactacagtgtggaaccaggccattcagcccattcggtcacacaccgacccttcaatgagcctcctaaccagaaccacaccttctaccctaacccggtaagtctccatttcgaa
tggctagcccaccgagcctgtgggctgtgggaaggagtctgtgtgggtgtgaggaaaatatacaaaactctacacagtcacacaagggtggaatcgaacccgggctcttggcgctgtgaggtggcagtggtaaccaccgagcctcgtgccacatcgaaaggtggtgattgctct
ggcggggtgggtatagaggaaatttaccaggatgctgcctcagaatgaagtacttcatcaatgaagagggacttgaggaggttacagagatagggagggggtgtaggggctggaggaggttacagagatagggagggggtgtaggggctggagggggtgacagagatagagagg
gggtgtaggggccggagggggttacagagatagggagggggtgtagggactggaggaggttacagaactagggagaggtgtaggggctggagggggttacagaactagggagggggtggaggggctggagggggttacagagatagggagggggtgtaggggctggagcgggtt
acagaggtagtgacaggttgaggggctggatgtggttacagagattgggaggggtgtgggagctggagggggttacagagataggaagggggtgtaggggctagaggaggttacagaactagggacaggtgtaggggctggaggaggttacagagatagggagggggtggaggg
ggctggaggaggttacagagatagggagggggtgtaggggctggagggggttacagagatagggagggggtgtaggggctggagggggttacagagatggggatgggttgtatgggctggacggggttacagagatagggagggggtgtaggggctgtaggggattacagagat
agggagggggtgtaggggctggaggacattacagagatagggagggtgtgtaggggctggaggcagttacagaggtggaggggtgaaggggctggagggggttacagagatagggagggggtgtaggggctggaggaggttacatagatagggaggagtataggggctgtaaga
ggttacagaaatagggaggggatttagggactggagggggttacggagatagggagggggtgtaggggctgggggggttacagggatagggaggggttggatgggctggacggggttacagagatagggagggggtgtaggggctgtagggggttacagagatagggagggggt
gtaggggctggaggaggttatagaggtgggggtgaaggggctggagggggttacagagacagggaggggtgtaggggctggagggggttacagagatagggagggggtgtaggggctggagggggttacagagatagggagggggtatagggtctggatggggttacagagat
ggggggctgtaggggctgggaggggatacagagatagggagtgggtgtcggggataacagagatagagaaatatgaggtgaaaaagcaacttgaagagacccattgaaagtttaaggtcagtgcggacatccataaacaggaaaacttcaagtgaataaacaattagcagagc
aattgttaaaagtggtgctcagtctttgaggacggaaatgcatattcagcatgtttcaaagaagaaaacaggcaggtctcaactgtcatgctttgctcgattgtaaagctgtggtcagaggaggcatatcattaacaggcattttcttattcatttcagatacgttggagccga
aggtggttgaagtcagtgtggaggggaatgcaggcggagggagggccctcctcactgatggcaactattgtgagtgtaaaggtaagcagtaatgtcatagcaacttcgagaataggcagtttgcaaactaaagataaaataagggattgcataaacactggaaacagtcacacc
aggggctggataaacacaggaaacagtcacaccagtggctggataaacacaggaaaagtcgcaccaggggctggataaacacaggaaacagtcacaccaggtagagggtaaacacaggaaacagtcacaccaaggattggataaacacaggaaaagtcgcaccaggggctggat
aaacacaggaaacagtcacaccaggtagagggtaaacacaggaaacagtcacaccaaggattggataaacacaggaaatagtcacaccagtggctggataaacacaggaaaagtcgcaccaggggctggataaacacagaaacagtcacaccagggataggataaacacaggaa
acagtcacaccaggtagagggtaaacacaggaaacagtcacaccagggataggataaacacaggaaacagtcacaccaggtagagggtaaacacaggaaacagtcacaccaaggattggataaacacaggaaatagtcacaccagtggctggataaacacaggaaaagtcgcac
caggggctggataaacacagaaacagtcgcaccagggactggataaacacaggaaacagtcacaccaggcagagggtaaacacaggaaacagtcacaccaggaacaggataaacacaggaaacagtcacaccagggctggataaacacaggaaacagtcacaccagggactgga
taaacacaggaaacagtcacacccgggattggataaacacaggaaacagtcacaccagggactggataaacacagaaactgtcacaccagggattggataaacacaggaaacagtcacaccagggtttatataaacacaggaaacagtcacacccgggattggataaacacagg
aaacagtcacaccaggaacaggataaactcaggaaacagtcacaccagggactggataaacacaggaaacagtcacacccgggattggataaacacaggaaacagtcacacgagggactggataaacacagaaactgtcacaccagggattggataaacacaggaaatagtcac
accagggtctatataaacacaggaaacagtcacacccgggattggataaacacaggaaacagtcacaccaggaacaggataaacacaggaaacagtcacaccagggactggataaacacaggaaacagtcacagcaggaacaggaaacagtactacatagaaatggcaagcaga
tagtaacaactccttccaagttccctttctgaaagaaaatcagaacattctggaaaagctctgcctgtctggcaacatctgtgaagaaaaaaagtcagagttgttctgagaaaagggtcagtgggcccaaactgttaactctgatttttttttcactatgctgcccgacccact
gggtttttccagcagcttctgtttttgctcctgatacacagcattccgcggtttctctcggttcccctttctgaaatggcaccgggctctttttttgaattgttgacaacagctggttgccctattctctacagttggccacacccagcttcgatacaaaccacaacagcccctt
cacagggcaggaaagaagtgcttgtgtcagcaaccaagcaggacccgccagaggaattggaaaaacggtaaatgctcattttttggtctgattcgtgtgcagcaggattctccctgtgagatcccgggattactccgggttttaaaactctcttttactgtaggcagccttgctt
tgtttgtctatttgtgatggcattgtatctagtgccttttggcttttaaggtgttgggttcgaaccccactcctgatggttacattctggaaaggtatcatcattacttgggggggggggtagctctgtatttatgagagattaatttatctgcatgacgtgtcaaacccata
taaggtaacatcatacgatgttatcctggccagtgtttatccctcagccgacctcattaaaacaggcgacctggaagttgtcactttactgttaatgggagcttgctgtgcgcaaactggtttcctacattgtagcagtgacaggcatcccagtctcattcaactgtctgcgcc
ccctgcaattatcccactggggtgcatgtagcagagttcagtttcacaattggctcccatgctgaggtcaggagtggggatgctcgctgatgatcgcgcaacgatcagcgccattcctgactcctcagatactgaagcagtccgtctccaagtgcagcgagaccaggacaatat

tcagatttaggctgacaagtaacgcttgcgcattcagtgtgattgagtcaaaatcatgctatccctcccaaattgcattatgcatctaaaagcagcatgtggactacagcgattcaagaaggcagctcatcttcaccttctcaatggcaactaggagtgggcaataaatgccgg
cccagctagtgacacccaagttttatgagtgagtaaaaacaaaatgccagacaaggactatctgaaatgcaaaagattctcaccatcattcccttggttttgagtgacattcccatcactgaatcccccactgtcagcactctgcaggttatcatcgaccaggcactctgctgg
attcaccatataagcacaatgtctacaacaccaggtcagacgcaaggaatactacagtgagtgactgacctcctggctccctaaagcctctctacaaggcacaagtcaggagtgtgatggaatactccccacttgcctgaatgagtgcagctccaataacactcaagaaacatg
atgccatccagggtaaagcagccccgtttgattggcatcacatcctcaaacatccaatccccccatcactgacgctcggtagcagcagtgtgcactgcctacaagatgcactgcagtaattcaccaaagatcctcagagcgcaccttccaagcccacaaccacttccattgag
aagaacaagggtaacagatacatcggaacactgcctcctgcaagttcccttccaaaccactcaccatcctgacttgaatgtataccgcaggtcaggcagggtgtaaacccaaaacgttgactctcctgctcctctgatgctgcctgacctgctgtgttcctgcagctccacact
gcattgaccctgactccagcatctgcagttcttggaatcttggaaatatatcgccctttcttgagtgtcgctgggtccgaatcagtggcatctccctacatcccaaagaatgcaacggttcaaggtagcagttcaccacctccttccgcagggcaacatgggaatgggcagtaa
attctggcccagcttgtgatgcccatatcccaagaatgaataatgcaatatatcaatgtcaaaaggcagatgtcagaacgcaggaggtgtaggctttgatcacgagtctcaagtcctctgtgatgctgggaaaagtttcaaaatgaaggagtctctgggtcatactaacaagag
aaagagaaagcccattgactcatcaagatctatccaaccacattgccaatttggaatgtagaggttccctgctacccgatcatccattaaccccatccttcaagaattaaaaaaagtcccagtttcatctcctcagactgatgaaacatgcagtgcagcattcccctcctaaac
caaataggagatatttgggtgggcaattatttcccattgactgagagtccagagtctcgtgtgctgtgccacgataaagagaacagttttcggatcagaaagaaggggccttaacataatttcatctgagcagcaggcacatataatgcagccctgcctcaggaattaccctgg
ctgatgaagtgctgtgacccacgagtgggtctcgaacccacggcctcccagaatgcagcccagtgtgagctgaggacagctgacttgtcatttaaggaaaggagtgtcttggttgggaggtggtacactactgaagagagccccccctgccttcccattgctggtgcccataaga
tgtaggagctgtttgtgggagctttctgtgcgtagactgtcaattacgcatcaagaaacaaactctggatttgctgtgaaccgcttcagaatgtcctgatgttgaaaaggcactggatatttccatctcgttaagataagaggagcagcagggccattcatcccactgagtctg
ctccaccattcatgactggtccgatcatccacacccccccacatccctgccgttgccctgttcacccttgattcccccttactggttaaaaatctctgtctgccccaccctttgcagtatagaactctgtgcattcattcctctctgagagatgaaattcctcctcatctctgt
cttaaatgggcaatccctcattctgagattatcttctttggtggtaggtaccctcccacagcctctccacatctaccctgccaaggcccctaaatatcttgtttgtttcagtaaaattatctccaattcttccagactgcaaagtgtacagccccaacttactcaaactctcct
cacacaaaaatgcctcatttcagggcctggcctttgtgagcacctatctccatggggaaacctactcgctttcccagtgtttttctcggactagtccagagtggtgtttctccttaagagtttccgaactgatgtcatcagcagaatactcccatcagactctgggcaagtagg
tcaggttgccactcttccctgaatctcaatacttatttaaataattcacaagggagaacataaaagccaggaccacgagtgggccagctggcccttcgagcctggtcctccattcaataagatcatggctgatttgtcatagactcaggtccacttgtctgactgttcaccaaa
accctcgattccttcactgttcagaaatctatctttgccttgaaaacattcaataaggtagcctcaactgggcagagaattctacagattcacaaccccttttggcgaagaagttcctcctcaactcagccctcaatctgctcccccttattttgaggctctgccccctcattc
tagtttcaatcccccccctcgccccggcccagtggaaacgacgtccttgcttccatgttatctattcccttcataatgttatgtttctataagatccactccccacgccattcttctaaattccaaccttgtgaacatcctctgcaacccctccagtgccagtacatcctttct
caagtcaggagatcaaagccatacgcagtactccagcgccctataaagctgcagcataacttccctgtttttaaactccatcccttgaggaatgaagaagaacattccatttgtatgggcaagtatgggtgtgaatccctcgccaaatgcctgtgcccatataggtgtcaagaa
gaacgatgtgcatttatatagcgataatggtgtcaaccccgaggagcttctccagaaatattagccaaaggattgcagacagcacagtggctcagtggctggtgctgctgcctcacagcggcaggggcctgggttcgattccaccttcaggtgactgtgtggagtttgcacatt
ctccctgtgtctgtgtggggttcctcccacagtccgaagatgtgcaggttagggtggatcggccatgctaagctgcccacagtgttcagggatgtgtagattaggtggggtacagaggaatgggtcagtgcgacttgttgggccaaagggtctgttcccacactgtagggattc
taagatatctgacgccaaataaaacaggttatgatagatttctgaagaagggccccgacccgaaacatcagctttcctgcttctctgatgctgcctggcttgctgtgctcctccagccccacgctgtgttatctcaggcttcagcaggttataattgactggtggaaggatacc
ttaaggaagggagggagttcgtggaggaggcccttgttgcaaaaggtaaattgatcagttagaagtacacgttgaagaagctgggactgttctcctttgagacaaaatggctgacgtgtgatataatggagggtttcaaaatcataagtgggttacaccaggtagatcaggaga
gactattcctactcataaaacaactgagagcaagagatttaaaggagtgtgcaaacataggaagggtgacatgaggaaatatcttttttttgcacagtgagtggtttgggtctggagtgcattgcctggaaatgtggtggaggcaggctccacgaagacattcacaatgttcctt
aggtacttatttggatcgaaatggtgtatagggattcgagtaaaatggcatgggttagaagtgattcctggtttagaaatgggttccttcctctgtacgctcacaattcatagagttgcacagccctttggtacaaccagaacagaccctcagtccaatttgtccatgctgacc
agacaccttaaattaacctcgttccatttgcaagcattggacccatatctctctaaacccttcctattcatgtatccatcaagagccttttcaatattgtcattgtactagcctccaccacttcttctagcagctcattccacacatgcaccaccctctgcgtgaaaacgttat
cccttgggtcttgtttaaatctttcccctctcaccttaagcctatgcactctggttttgaactcccctatcttgggaaaaacaacctggctattcaccctatccaagccccccatgttttcataaacccctttaaggtcccctctctgtctctgatgctccagggaaaatagcc
ccatcctctcccgaaagatcaaacactccaaccccggcaacatatttgtagatcttttccaaaccctttataatttcgcttcctatggtagggagaccaaaattgaatgcagtatttcaacagtgcctaactaatgtcctgtatagccccaacatgacctccaatactcaatgc
actgaccaataaagtaagcgtaccaaatgtcttttgtaccaccttgtctacctatgactccttcaaggaattatgaacgtgcactccttttgttcagcaacactcctcaggaacttagccttaagtgtataagctgagcttagttgccagaatacagcacctcatatttatcta
aattaaactccatctgccactccaagttgatttgtacacaagtcgaaacacaatgtggtacaaacggatgcgtgcatgtcagatcctttgaattaatctgaatacggaatggcgtccattagtgagagtgtttggatatttgtaaattggggactgtctgggtttagagccgga
gaagacatgatgggcagactgacctccatcctgaacgatatagtaccttatctctcatttgtttctttcggaggaagtggcacactggccagacacagcccttattgcctaattgccccttgagaaagtgggggtacgctgtcttcttgaattactgttgtccttgggggtatg
tagggtcacccatggggctgtcacgaaccagactggggagggatttctaggattttgatccaatggtggtgagttaacgaggatatttatctaagcaacggcgatgagcaactgaaaggacaatgcagagatgatggttttcccatctattcccagcagaaggggttttgcatg
ttttggaaggatgtaaaacaaaattggtgaatttctgtaatgtatcttatagatcttccttttaaccattcatgggatgaggctgtctctagctaagcatgtattaccaacccgtaaattgtactgaggacagttaagagtcaatcacatcaccgtgggtctgggaccacaata
aggccagaccagctaagggtggcagctttgttccctcatccaaataggctttgtccctccaacaatttctagccttgattccaaattttcctgtagcccaatttccaccatggtgggattcgaacacagctctccaggatgttacctgggtcactggattaacagtccaatggt
actaccactaggctgtcacctccccactagattcagagggtacactctgcaccttttaaagtggtcagcatcatggttttagaagctgctgtctaaggatcattgccaaatttgtgcagtgcatcttgtagatgttgctgttataatactatacttaacaggcccctttttttt
gaagagaagttttaaggcagaggcactgttcaatctaccatgtagcctcttgagtaaacagcttgtgagatgttgtttttttggttaagttgtaacaatagaagcagactggctgggtgtgatctagctccccatgaacctggatttttacttttagctgttagcagtagactt
gaagaactggaagctattttgccctgtctcggtattgccaaaagctgggggggggggggggggggggagtacggggtaggtgcggagggggtggttctcctcctgctgctggagttgcatgtgaggtaatctgttttctgattttgccttttttttgccaggagtgggatattac
aatattggaacagttaattacaaatagtttctgtgtctgttatccttgatatcagtgataatgggaactgcagatgctggagaatctaagataatgaaatgtgaagctggatgaacacagcaggcccagcagcatttcaggagcacaaaagctgaccaagggcctaggcccgaa
acatcagcttttgtgctcctgagatgctgctgggcctgctgtgttcatccagcctcacatttcattatcttgtgtcagttatcctgtttagtttttgcaatagagtcgagctcttccaattccttctttctttcgttgcattttaactattgtggatgaataaaatatgttcta
ctttaaatctggaagcttgacaaatcaaattgcaactggaatgccacaccttacatttacttttacaataaaaaaagttggggtctagactttcttctcaatactttgaagggtttggtctagtctataacagcacacactgcaaaatttagattggaaactcatgacttttat
tctgtttttccttgtgtagctttcctccccaagctgcgaaaacagaaacgttactgcctgaaaccagggaaaactatatcgaaaggcaagcgacacttctattgtgtggagacttggcacagaatgaaaaaatgcacccccctgtaaaccaatggattttatccccacgcccctt
taagattgcatgctgaggcagggagatcttacgcggcaaattagacagacgatccctgcaccaaaagttgcaaattcttggaatataaagccattatcctcacaagccaggagagagcaatcactagattagaccccactctgctgccttcccagacccattcttagattatgt
aggccagtgatggacacccccagtgcttatgctgagtgagtgcagcactgcctgactctctcccagctaacttccttccccatacaccacttctctaccctggatgcaaaaatccatggttcactctcttccgaaacagaaaaccagaataaaaaggccgggtgggcactgtct
cctcggaaccatgatgcaaaatgtcttcaatgtccctcttcaaaaagtttaaaaggaaaatgcacatagatacagtgatcccatcactcatgggcctgtgagatgttactgtgcagtagcctggtgtcccacccaaaaatggagtctttgcattagacacaaagcctgacctta
cgaaacggcaataagtcctcccatcacaagatttataatccattttgtgagctgagccatcattcccagtccgttgaatattcacgagtctccagaacacagcaatctcttgtcagctaggatctagccagcgaaactttaacctgggattgtaacgtggagattttcctcacc
agaccttcggagatgatctggaggcatgagtgggatttgaaaccaggcatagcctcaatctaagaaactcccttggaggctatcattgagccgaactcactcaaccctgtttcccagccccttgtcctgtgttttctttatatccaactgaaatctagactcataccaggaatg
agtaggttgtcttatgaggaataacaaaacagactaaacctccatactttggagtttagatgaatcagggctgactttatttgaaacatataggaccctgagggatgcttaactgactggatgtggagagaatgttcccacttttaagaaaatctagaaaaaaacgggttcatg
tttaaaattaagggagcactcattgaagacagaggtggggaaacatttctttctctatcagagtgttctgaatctttggaattcccatcttcaaaaggcaatggatgttctaatctttaaatatttcgaaggcagaggtgggtagaatcttgacgaccaatggggagtcagaga
ttatcggggacgatgcaggaacgtagagttgaggttaaaatcggatcagccatgaccctattgagtggccgagcacccatttgttcctgagataatgggaactgcagatgctggagaatccgagataacaaaatgtggagctggatgaacacagcaggccaggcagcatcttag
gagcacaaaagctgatgtttcaggcctagaccctttatcagagagggggatggggaagggttctgaaataaacagggagagagggggaggcggaccgaagatggatagaggagaatataggtggagaggagacagacaagttaaatggatggagccagtagaagtgagtgtagg
tggggaggtagggaggggataggttagccatgcttgtgatgtcggaagggcatttcctgcctgtgccagccatggcaatttaaccagattttgagctctcagccacttcaggatctctctgatggatccatgaccttccaagtccatactgaagagaggtcttaagttggcatt
gtgtcatctattttgttaaacctcgggaggctaaggaaattggttatttccataaagactcttaccaatttttatggatgctccataaaaagcattccaactcgatgtattacagcttggtacagctgctgctctgtccagcattgtaagaaactacagagatttgtgaacaca
gcccagtctatcacacacaccagccttccatcctttgactccatctgtacttcccacggccttgggaaagcagccaacatcattaaagacccctcctaccctggttatactctcgtccgacctcttctattgggtagaagcgacaaaagcgtaaatactaaaacggattcgcgg
acagtttcttccccattgttattaaaattctgaatgaatctctcaaattttaaatttaatgttgattttgctctctacgtaccttctctacagccgaaacattatattcctcactctgttctgttgccctaatgtactttgaatggtatgatctgcctggactgcactcagaac
aaaacatttcgctgcacttaggtacatgtgacactaataaatcaaaccaaaccaagtcaaatgtagatgcttctcacaggaaaccaacaaaggctatggtcgagcttaatggtctgctagaatgctgagtctgggtctagccccttgatacagcttgtatagtgttgctacagg
gagtcctcactatcagtttacactaatgagtctttccccaccaatgcgatgtttagtcgttgagtggaacataggctgtaaattggtgtaactccaggggtctctgaacagagcaggagtcttaaccagaagaaagacaggaagagaatcgaaggcattcctgatcattcctg
ccaccaaaatgcactcaacgttgatagggagggtcaaagtgacaccttcctgtttgtagtggaagggaagggagttacagtcgaaaacatgtagggttggattctcatctgtgtcacattgaaccagtgtgtgaatgccaccctcactgggatgggcatcaattttactctctc
catggattttgacataagcccagcgttagctgctcagcgctgatggccctttttctgatctcagcagccacttagttggcacgattttggcccggcaaagaattaaacgacatgatgccagcttaagagggcatttttttttccttatgcctttgtgggatgtagaccgaactgg
ctagaccagcatttactcaaactccctagttgccccacaaataggagtcaaccatgtcactgtgagtctggagtcacatgtaggcttgacagagtaagatggcagatttcctttcttaaggttagacattcagttgttttttttcaaaataatggctaatggtcatcatttctg
tttgagttaaacttttaccatctgctattgtgagattcaaagtcatgtttccagaatattaccctgaatttctgaaccaacgacagtgccactaaaccactgcctgctctgaaagaggttcttagtttttcaatggagcccacatgtcatttctccacatgtatcgcttgtgct
gagaaaaaatatttactatatcagatagttacagtaagattgctatcctttaaagaattgttcatttattcattagtcagctacaaactatgtccttatgctctgatgtaaagcgtacaattcatctatattttaatattgcttagtttgaatttttggattttgaactggtgg
cgtctgggattaaatgattaacatgtgagtatttctggtgattgcttttaatatatcatataagagacagattttgtggagagtaatggagttttgttttcattgagtattacaagcgaacagaataaacagtttagcattgggtttcggctggtaggatcttgtaattacttt
atttttgactcagaaggagccccccattgattaaagaaagggccatcggcttttgttttcagcagatgtggttgtcactgggcatgcaaggtagttgctgtcaaacagttaagaaatagattgcctcagtgtaaggaatttcatgtgaatgcaccagaccagaagggcttggt
taaaatcctgaagcagtaccgcaaagctgagaaaatctaagcttgcataaagatcaaggaagaggctatcaaattctctataccagcaactgaagccaaagttaaatcaaactctgtaagtgatgagaagggaatttttttctggcatctgaatgctttaacaggttactgttg
ggattaatgggattatgttttattgcattttaaaatcttaaatgtgtgtaatttgaaaacacctttgctttattctgtgttactttcattcattttgcataataaacttctgttctat

### C: Sequence of cDNA (coding region in red and highlighted in yellow)

agaggctctgattaaacagctcgggaccagtgaacacccgaagatgcagatcacactcctgcttattgcctttgcctttgcatcgatccctgtgctcaccgacgatacgttggagccgaaggtggttgaagtcagtgtggaggggaatgcaggcggaggagggccctcctcac
tgatggcaactattgtgagtgtaaagttggccacacccagcttcgatacaaaccacaacagccccttcacagggcaggaaagaagtgcttgtgtcagcaaccaagcaggacccgccagaggaattggaaaaacgctttcctccccaagctgcgaaaacagaaacgttactgcct
gaaaccagggaaaactatatcgaaaggcaagcgacacttctattgtgtggagacttggcacagaatgaaaaaatgcacccccctgtaaaccaatggattttatccccacgcccctttaagattgcatgctgaggcagggagatcttacgcggcaaattagacagacgatccctgc
accaaaagttgcaaattcttggaatataaagccattatcctcacaagccaggagagagcaatcactagattagaccccactctgctgccttcccagacccattcttagattatgtaggccagtgatggacacccccagtgcttatgctgagtgagtgcagcactgcctgactct
ctcccagctaacttccttccccatacaccacttctctaccctggatgcaaaaatccatggttcactctcttccgaaacagaaaaccagaataaaaaggccgggtgggcactgtctcctcggaaccatgatgcaaaatgtcttcaatgtccctcttcaaaaagtttaaaaggaaa
atgcacatagatacagtgatcccatcactcatgggcctgtgagatgttactgtgcagtagcctggtgtcccacccaaaaatggagtctttgcattagacacaaagcctgaccttacgaaacggcaataagtcctcccatcacaagatttataatccattttgtgagctgagcca
tcattcccagtccgttgaatattcacgagtctccagaacacagcaatctcttgtcagctaggatctagccagcgaaactttaacctgggattgtaacgtggagattttcctcaccagaccttcggagatgatctggaggcatgagtgggatttgaaaccaggcatagcctcaat
ctaagaaactcccttggaggctatcattgagccgaactcactcaaccctgtttcccagccccttgtcctgtgttttctttatatccaactgaaatctagactcataccaggaatgagtaggttgtcttatgaggaataacaaaacagactaaacctccatactttggagtttag
atgaatcagggctgactttatttgaaacatataggaccctgagggatgcttaactgactggatgtggagagaatgttcccacttttaagaaaatctagaaaaaaacgggttcatgtttaaaattaagggagcactcattgaagacagaggtggggaaacatttctttctctatc
agagtgttctgaatctttggaattcccatcttcaaaaggcaatggatgttctaatctttaaatatttcgaaggcagaggtgggtagaatcttgacgaccaatggggagtcagagattatcggggacgatgcaggaacgtagagttgaggttaaaatcggatcagccatgaccct
attgagtggccgagcacccatttgttcctgagataatgggaactgcagatgctggagaatccgagataacaaaatgtggagctggatgaacacagcaggccaggcagcatcttaggagcacaaaagctgatgtttcaggcctagaccctttatcagagagggggatggggaagg
gttctgaaataaacagggagagagggggaggcggaccgaagatggatagaggagaatataggtggagaggagacagacaagttaaatggatggagccagtagaagtgagtgtaggtggggaggtagggaggggataggttagccatgcttgtgatgtcggaagggcatttcctg
cctgtgccagccatggcaatttaaccagattttgagctctcagccacttcaggatctctctgatggatccatgaccttccaagtccatactgaagagaggtcttaagttggcattgtgtcatctattttgttaaacctcgggaggctaaggaaattggttatttccataaagac
tcttaccaatttttatggatgctccataaaaagcattccaactcgatgtattacagcttggtacagctgctgctctgtccagcattgtaagaaactacagagatttgtgaacacagcccagtctatcacacacaccagccttccatcctttgactccatctgtacttcccacgg
ccttgggaaagcagccaacatcattaaagacccctcctaccctggttatactctcgtccgacctcttctattgggtagaagcgacaaaagcgtaaatactaaaacggattcgcggacagtttcttccccattgttattaaaattctgaatgaatctctcaaattttaaatttaa
tgttgattttgctctctacgtaccttctctacagccgaaacattatattcctcactctgttctgttgccctaatgtactttgaatggtatgatctgcctggactgcactcagaacaaaacatttcgctgcacttaggtacatgtgacactaataaatcaaaccaaaccaagtca
aatgtagatgcttctcacaggaaaccaacaaaggctatggtcgagcttaatggtctgctagaatgctgagtctgggtctagccccttgatacagcttgtatagtgttgctacagggagtcctcactatcagtttacactaatgagtctttccccaccaatgcgatgtttagtcg
ttgagtggaacataggctgtaaattggtgtaactccaggggtctctgaacagagcaggagtcttaaccagaagaaagacaggaagagaatcgaaggcattcctgatcattcctgccaccaaaatgcactcaacgttgatagggagggtcaaagtgacaccttcctgtttgtag
tggaagggaagggagttacagtcgaaaacatgtagggttggattctcatctgtgtcacattgaaccagtgtgtgaatgccaccctcactgggatgggcatcaattttactctctccatggattttgacataagcccagcgttagctgctcagcgctgatggccctttttctgatc
tcagcagccacttagttggcacgattttggcccggcaaagaattaaacgacatgatgccagcttaagagggcatttttttttccttatgcctttgtgggatgtagaccgaactggctagaccagcatttactcaaactccctagttgccccacaaataggagtcaaccatgtca
ctgtgagtctggagtcacatgtaggcttgacagagtaagatggcagatttcctttcttaaggttagacattcagttgttttttttcaaaataatggctaatggtcatcatttctgtttgagttaaacttttaccatctgctattgtgagattcaaagtcatgtttccagaatat
taccctgaatttctgaaccaacgacagtgccactaaaccactgcctgctctgaaagaggttcttagtttttcaatggagcccacatgtcatttctccacatgtatcgcttgtgctgagaaaaaatatttactatatcagatagttacagtaagattgctatcctttaaagaatt
gttcatttattcattagtcagctacaaactatgtccttatgctctgatgtaaagcgtacaattcatctatattttaatattgcttagtttgaatttttggattttgaactggtggcgtctgggattaaatgattaacatgtgagtatttctggtgattgcttttaatatatcat
ataagagacagattttgtggagagtaatggagttttgttttcattgagtattacaagcgaacagaataaacagtttagcattgggtttcggctggtaggatcttgtaattactttatttttgactcagaaggagccccccattgattaaagaaagggccatcggcttttgttt
tcagcagatgtggttgtcactgggcatgcaaggtagttgctgtcaaacagttaagaaatagattgcctcagtgtaaggaatttcatgtgaatgcaccagaccagaagggcttggttaaaatcctgaagcagtaccgcaaagctgagaaaatctaagcttgcataaagatcaagg

aagaggctatcaaattctctataccagcaactgaagccaaagttaaatcaaactctgtaagtgatgagaagggaatttttttctggcatctgaatgctttaacaggttactgttgggattaatgggattatgttttattgcattttaaaatcttaaatgtgtgtaatttgaaaa
cacctttgctttattctgtgttactttcattcattttgcataata

**D:** Amino acid sequence (signal peptide shaded)

MQITLLLIAFAFASIPVLTDDTLEPKVVEVSVEGNAGGGRALLTDGNYCECKVGHTQLRYKPQQPLHRAGKKCLCQQPSRTRQRNWKNAFLPKLRKQKRYCLKPGKTISKGKRHFY
CVETWHRMKKCTPCKPMDFIPTPL

**Fig. S6.** Gene position and architecture (**A**), genomic DNA sequence (**B**), cDNA sequence (**C**), and encoded protein sequence (**D**) of the unannotated *cxcl17* gene in the NCBI reference genome of *Stegostoma tigrinum*. The information was downloaded from the NCBI gene database (https://www.ncbi.nlm.nih.gov/gene).

**A:** Gene position and architecture

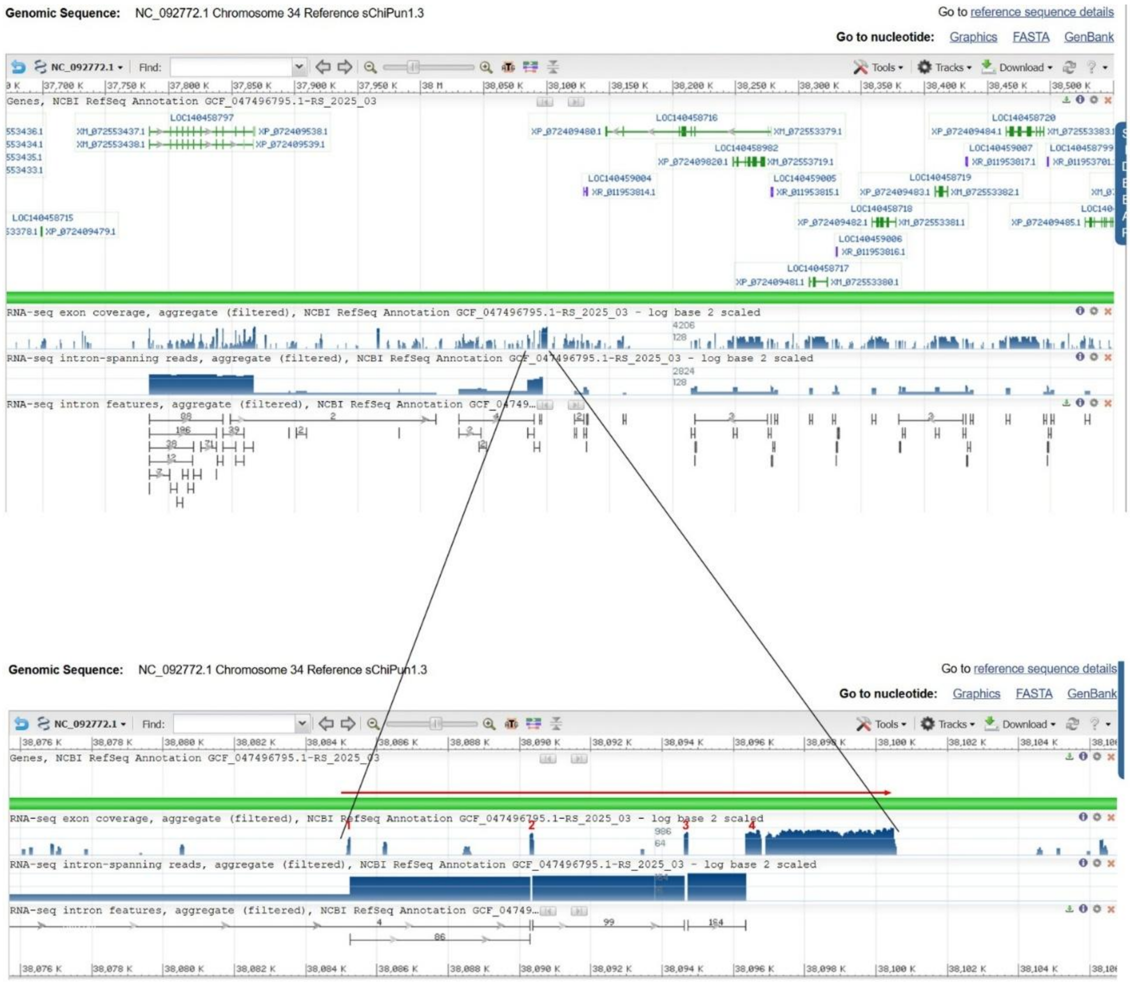


**B:** Sequence of genomic DNA (exons in red and highlighted in yellow, coding region underlined)

agatataaatctcccagcgctcagaggttcgcacgaaccagcctggggagcagggaaacaccccaaagatgcagcttgccttgctgctgctgatagtgtgcgtctccagctacaatcctgttctcatggagggtagggccaatttacaattaatcagttgggtagtcactgtct
gttcaggaatgcaggacttactggtagtgggggaggtgggggattggggaagggagaggaggaaggagggggggaaggggagaggggaaacgggacaggacaggaggtgggggtgtaaggggactggaggaggagggtggcaatgcccagggtggtcactggtcatggtca
gcgatcagtcagagcaattatttccccccacacttaccaactcagggatgggaccccctaattccctcccaatcctccctatatagcagggcttatacacttaatggttcggtcctagggagtgttgctgaacagagaccttggagtgcagatccatagctccttgaaagtggag
tcgcaggtagctaggataatgaagaaggtgtttggtatgctttcctttattggtcagagtattgagtacaggagttgggtgctgaaaatgtgttgctggaaaagcgcagcaggtcaggcagcatccaaggaacaggagaatcgacgtttgaagaagggcttatgcccgaaacgt
cgattctcctgctccttggatgctgcctgacctgctgcgcttttccagcaacacattttcagctctgatcgccagcatctgcagtcctcactttctgctacaggagtttgggaggtcatgttgaagctgtgcagggtattggttaggccactgttggaatattgcgtgcatttc
tggtctccttcctatcggaaagatgttatgaaacttgaaagggttcaggaaagatttacaagggtgttgccaggattggaggatctgagctacagagagagaggctgaacaggctggggctgttttccctggagcgtcggaggctgaggggtgaccttatagaggtttataaaa
tcatgagtggcatggatagggtaaatagacaaggtcttttccctggggttggggagtccagaactagaggggcataggtttagggtgagaggggaaagatataaaagagacctaaggggcaactttttcacacagagggtggtacgtgtatggaatgagctgccagaggatgtg
gtggaggctggtacagttacagcatttaaaaggcatctggatgggtatatgaacaggaagggttcagtggtatatgggccgggtgctggtaggtgggactagattgagttgagatatctggtcggcatggacgggttggaccgaaggtctgtttccggctgtacatctctct
gactcgatgactcttgtagcagccgtggagattaagcagaattaacgtttgggatcggtgcagacccaatggcttccttccaccccagagggatttcctgtctctagttccagcagctcacctgacctttaacctctgcactgaggtacttgccatgggctgagcgaactcagt
gaactcagaaaggcactccgggacagagaggggaaggcggggggggggggggaagagatttgaatgaccacttgcccgcctggtgcttgacctgccctggcctcctgtaaattcgaagaccgagccagtgactcaatctgattacaggccctgaatcagaccaggagagacggc
cagaaatcaggccattcaggcccatctagtctgctccaccatccaatcatggctgataaagcttctcaacctcccgctttctcccccctgacccttcacccccttgatactcaggaacctatccattttggtcttaactatactcaatgaccaggcccccacagccttcagtggc
agtgaatctcatagattccccactctctggctgaagacgtttctgcttatcttcattctaaaaggtcttccctttagtctaaggctgttccctcgtccctcttatcgatgggaacatcttcccaacatccactctgtccgggccattctataagtctcaattagatccccccc
ccatccttctaaattccatcgagtataaacccagagtcctcaaatgttcctcatctgttaagctttccattcccgggaccattctcgtgaatctcctgtgaaccggccagtgtatccttcctgagatatgacgcccaaaactgagaacaatcctccaaatctgacctgaccaca
gcctcagagacactcagaagtccatccctgcttttatattcaagtcctcccaagttaaatcacagcgttaggggtggcagggtggctcagtgggttagcactgcaacctcacaacaccaggatcccaggtttgattccagcctccgagcgactgtctgtgtggagtttgcacat
tctccccgtgtctgtgtgggtttcctccgggtgctccggtttcctcccacagtccaaaggtgtgcaggtcagggtggattggccgtgctaaattgtcccagagtgataggtgcattagtcagagggaaatgggtttgggtgggttactcttcggaggtcggtgctgactggtt
gggccgaagggcctgtttccaatctgtagagaatctaatctaatcttaaaaaaaaatctaatcgttgcatttgccttccgaatgactgactcaattgtcgacacctctttcattattagagtattcggaaacaagctcagacacagttataagctggagcaggagatctttatat
accagccagtacagagcaaactatcaataaatacacaggcagggagggctggagggggttacagggatagggagagggtgtaggggctgggggggttacagggatagggagagggtgtaggggctggagggggttacagagatagggagggggtgtaggggctggagggggtt
acagggatagggagagggtgtaggggctggagggggttacagggatagggagagggtgtaggggctggagagggttacagggatagggagggggtgtaggggctggagggggttacagggatagggagagggtgtaggggctggagggggttacagagatagggagggggtgta
ggggctggagggggttacagagatagggagacggtgtagggctggagggggttgcagagatagggagggggtgtaggggctggagggggttacagggatagggagggggtgtaggggctggagggggttacagggatagggagggggtgtaggggctggagggggttacaggga
tagggagggggtgtaggggctggagggggttacagagatagggagggggtgtagaggctggagggggttacagagatagggagggggtgtaggggctggagggggttacagggatagggagggggtgtaggggctggagggggttacagagatagggagggggtgtagaggctgg
agggggttacagggatagggagggggtgtaggggctggagggggttacagagatagggagggggtgtaggggctggagggggttacagagatggggagggggtgtaggggctggaggggggttacagagatagggtgggggtgtaggggctggagggggtttactgggagagaga
gtgagtgtaggggttggaggggctgacagagatagggagggatgtaggggctggagagggttacagagatagggagggtgtgtaggggctggagggggttacagagatagggagggggtgtagggtctggagggggagacagagataggagggggtgtaggggctggaagaagtt
acggagatagggagggggtgtaggggctggaagaagttacggagatagggagggggtgtaggggctggagggggttacagagattgggagggggtataggggctggagggggttacagagatagggagggtgtgtaggggctggagggggttacagagatagggagggggtgtagg
ggctggagggggttacagagatagggatggggtgtaggggctggagggggttacagagatagggagggggtgtaggggctggagggggttacagggatagggagggggtgtaggggctggagggggttacagagatagggagggggtgtaggggctggagggggttacagagat
agggagggggtgtaggggctggagggggttacagagatagggagggggtgtaggggctggagggggttacagagatagggagggggtgtaggggctggagggagttacagagatagggagggggtgtaggggctggagggggttacagggatagggagggggtgtaggggctgga
gggggttacagagatagggagggggtgtaggggctggagggagttacagagatagggagggggtgtaggggctggagggggttacagagatagggagggggtgtaggggctggagggggttacagagatagggagggggtgtaggggctggagggggttacagggatagggaggg
ggtgtaggggctggagggggttacagggatagggagggggtgtaggggctggagggggttacagggatagggagggggtgtaggggctggagggggttacagggatagggagggggtgtaggggctggagggggttacagggatagggagggggtgtaggggctggagggggtt
acagagatagggagggggtgtaggggctggagggggttacagagatagggagggggtgtaggggctggagggggttacagagatagggagggggtgtaggggctggagggggttacagagatagggagggggtgtaggggctggagggggttacagggatagggagggggtgta
ggggctggagggggttacagagatagggaaacatacggtcatcgatcaattgaaagaaaataatgaattgatgaagttgaagggatttccagagcagaacgcaacgtgattgtccataatcaggaacgctgcaactggagataaatctatcaggatttcagcaagaagtgatgt
tgaatctttgctcagacgaagagatgttccgcatcattcaacaaaacaatgcccaggtttggaacgtagaggttggttggatgatccggtaaagttcagtggtggcacatttttttcaccggcattgtgttcatttcagatattcgggatgggaaggaagataaacccagggtgg
aactcagtgcaaacacaacaggcaccttggaacccggcaggaaacattgcctgtgtaaaggtgggtgcagacagcccagcaacttcgagaccacgcgatactgtgctcagaaaaacaattacacattggtcacaccggggaccgggtcaacatggggaagggtcacaccgggta
aacacagggaagggtcacaccggggactgggacaacacggggaagggagacaccggggaccgggtcaacacggggaagggtcataccgggtaaacacggggaagggtcacaccggggaccgggtaaacacggggaagggacacaccggggaccgggtcaacacggggaagggtc
acaccggggaccgggtcaacatggggaagggtcacaccgggtaaacacagggaagggtcacaccggggactgggacaacacggggaagggagacactggggaccgggtcaacacggggaagggtcataccgggtaaacacggggaagggtcacaccggggaccgggtaaacacg
gggaagggacacaccggggaccgggtcaacacggggaagggtcacaccgggtaaacacagggaagggtcacaccggggactgggacaacacggggaagggtcacaccggggactgggacaacacggggaagggtcacaccggggaccgggtcaacacggggaatggccacaccg
gggaccgggtcaacacggggaagggtcacaccggggaccgggtcaacatggggaagggtcacaccggggaccgggtaaacacggggaagggtcacaccggggaccgggtaaacatggagaaggatcacaccggggaccgggtcaacatggggaagggtcacaccgggtaccggg
tcaacatggggaagggtcacaccgggtaccgggtcaacacggggaacggtcacaccggggactgggtcaacacggggaagggtcacaccggggaccgggtaaacacggggaagggtcacaccggggaccggtcaacacggggaagggtcacaccggggaccgggtcaacacgg
ggaagggtcacaccggggaccgggtaaacacggggaagggtcacaccggggaccgggtaaacacggggaagggtcacaccggggaccgtgtaaacacggggaagggtcacaccggggaccgggtcaacacggggaagggtcacaccggggaccgggtcaagacggggaagggtc
acaccggggaccgggtaatcacggggaatggtcaaaccggggaatgggacaacacggggaagggtcacaccggagaccgggtcaacacggggaagggtcacactggggaccgggtcaacacggggaagggacacaccggggaccgggtaaacacggggaacggtcacaccgggg
accgggtaaacacggggaagggacacaccggggaccggataaacacggggaacggtcacaccggggaccgggtaaacaaggggaacggtcacaccggggaccgtgtcaacacggggaagggtcacaccggggaccgggaaaacacggggaagggtcacaacggggaccgggtaa
tcacggggaatggtcacaccggggaccgggtaaacacggggaagggtcacaccggggaccgggtaaacacggggaagggacacaccggggaccgggtaaacacggggaacggtcacaccggggaccgggtaaacaaggggaacggtcacaccggggactgggtcaacacgggga
agggtcacaccggggaccgggtaaacacggggaagggtcacaccggggaccgggtaaacacggggaagggacacaccggggaccgggtaaacacggggaagggtcacaccggggaccgggtaaacacggggaacggtcacaccggggaccgggtaaacacagggaacggacac
tccggggaccgtgtaaacacggggaagggtcacaccggggaccgggtaaacatgggaaagggtcacaccggggaccgggtaaacacggggaagggtcacaccggggaccgggtaaacacgaggaagggacacaccggggaccggggaaacacggggaatgggtcacaccgggga

ccgggtaaacacggggaacgggacacaccggggaccgggtaaacacggggaaagggtcacaccggggaccgggtaaacacggggatgggtcacaccggggaccgggtaaacacggggaagggtcacaccggggaccgcgtaaacacggggaagggtcacaccggggaccgggta
aacacggggaaagggtcacaccggggacagggtaaacacggggaagggtcacaccggggaccgggtaaatacagggaagggtcacaccggggaccgggtaaacacggggaagggtcacaccggggaccgggtaaacacagggaatgggacacaccggggaccgggtaaacatgg
ggaagggtcacaccggggaccgggtaaacacggggaagggacacaccggggaccgggtaatcacggggaacggtcacaccggggaccgggtaaacacggggaaagggtcacaccagggaccgggtaaacacggggaagggtcacaccggggaccgggtaaacacgggcaacggt
acacaccggggaccgggtaaatacggggaagggtcacaccggggaccgggtaaacacggggaacggtcacaccggggaccgggtaaacacggggaagggtcacaccggggaccgggtaaacacggggaagggtcacaccggggactgggtaaacacagggaagggtcacaccgg
ggaccgggtaaacacggggaagggtcacaccagggaccgggtaaacacggggaagggtcacaccggggaccgggtaaacacggggaagggtcacaccgggtaaacacggggaagggtcacaccggggaccgggtaaacacggggaagggtcacaccggggaccgggtaaacacg
ggaaagggacacaccggggaccgggtaaacacggggaagggtcacaccggggactgggtaaacacggggaagggacacaccggggaccgggtaaacacggggaacggtcacaccggggaccgggtaaacacggggaagggtcacaccggggaccgggtaaacacggggaagggt
cacaccggggaccgggtaaacacggggaagggtcacaccggggactgggtaaacacggggaagggacacaccggggaccgggtaaacacggggaacggtcacaccggggaccgggtaaacacggggaaagggtcacaccagggaccgggtaaacacggggaagggtcacaccgg
ggaccgggtaaacacggggaagggtcacaccggggaccgggtaaacatggggaacggtcaccccggggacctggtaaacacggggaagggtcacaccggggaccgggtaaacacggggaagggtcacaccggggaccgggtcaacacggggaagggtcacaccggggagcaggt
taacacggggaagggacacaccggggaccgggtcaacacggggaagggtcacaccggggaccgggtaaacacaggaaagggtcacaccgagaaccgggtaaacacggggaagagtcacaccggggaccgggtaaacacggggaagggtcacaccgggtaaacactgggaagggt
cacaccggggaccgggtaaacacggggaagggtcacaccggggatcgggtaaacacggggaaaggacacaccggggaccgggtaaacacggggaacggtcacaccggggaccgggtaaacacggggaaagggtcacaccagggaccgggtaaacacggggaagggtcacaccgg
ggaccgggtaaacatggggaacggtcacaccggggactgggtaaacacggggaacggtgacaccggggaccgggtaaacacggggaagggtcacaccggggaccgggtaatcacggggaagggtcacaccggggaccgggtaaacacggggaagggtcacaccggggaccgggt
caacacggggaagggtcacaccggggaccgggtaaacacggggaagggtcacaccggggaccgggtaaacacggggaagggtcacaccggggaccgggtaaacacggggaagggtcacaccagggaccgggtaaacacggggaatggtcacaccgaggaccgggtagattataa
cagggaagatgttaaaatttggcccagtgtgttctgcccctgctcacaagctcccttgttgaagagttgctctctttaaccgaagggccatctctgatctgcctttctcctgtagctggtcacctcccgttacactaccgcgcaaagcagccattcgtaacggctggcagaaag
tgcttctgtcagaaacgaagccagatcaccaggaaaaagcacaaggatggtaaattgttctttttcactctgtctccagggctgagagggagggtgtcattggtgtacatcccttgggatttgagtgttcctgcgcatagcaattttgaactccatgagtctgaaagctctcca
tctccggatacagccttgccttttgtgattatctgcgacggcattgcatcccacaccctctagctcttcagatctcatgttctgatcccaaatctgatatgattctcatccaggctgtcattatcagcgcttgggtgtagctctgccctgatgagaggttaattcatttgtata
agttaaggtggtttgataatattctgccaatgtttatctctgctccgcagaaacagatgagctggtcattgtcacattgctgattgtgggatcttgctgaccccaaactggcgtcctgcattacagcagtattcaacatcctttcgatggtctgcgcccatgagattattccac
caggggcgctgttgttgacatgtagagaggaatgcggtctcataatcatgcgaaccacatcaaacccattgttaaagtgggaattggggatattcgccgatgattgcaacaatgctcagcaacatttgtgacagctcagacactgagacaatccatgtccaaatacagcaagac
ccggacaattcccacgctgggactgacaagtgacaggtcacattgacggtagcaccatctggtcgaaatccaagaattccctccctcggggcattgtgcccacagtggagggactgcagcggttcagccctgtgtctgcgtgggtttcctccgggtgctccggtttcctcccac
agtccaaaatgagcacgttagggtggattggccgtgctaaattgtcccatagtgcccagggatgtgtaggttagggtggattggccatgctaaattgtcccatagtgcccagggatgtgtaggttagggtggattggccatgctaaattgtcccatagtgcccagggatgtgca
ggttagggtggattggccgcgctaaattgtcccatagtgtccagggatgtgtaggttagggtggattggccatgctaaattgtcccatagtgcccagggatgtgcaggttagggtggattggctgcgctaaattgtcccatagtgttcagggatgtgtaggttaggttggattg
gccatgctaaattgtcccatagtgttcagggatgtgtatgttagggtggattggccgcgctaaattgtcccatagtgcccagggatgtgtaggttagggtggattggccatgctaaattgtcccatcgtgcccagggatgtgtaggttagggtggattggccatgctaaattgt
cccattagtgcccagggatgtgcaggttagggtggattggccgcgctaaattgtcccatagtgcccagggatgtacaggttagggtggattggccacgctggtatgagtgggtgaggtgcgtttttcagaggtcaacgtccattaattgggtcgggggaatggttgtggatta
ctgttcggagggtcggcgttggcatcttgggcggaagggcctggttcgtgcagggattctatgagtcatgcgtgtttttaaaaatgtcctttccacagctctcttcgcgagacagcgccaagggaaacgttactgcctgaaaccaggaaaggccgtgtaccaaggccagcgaca
attgtattgcgtggagagctgggccagctggagaaagtgtgccccctgtgacccggtcgcccttatccccaccccactttaagacagcgcgctgaagtatagggagctcccaatgccgcccggagactccctgcacggaccgttcggaattcccagacagcgaagccgatcatc
cccggctccaccaccctccggcctccccgcgacgccgcccagtgctggacatcccccagtgcccgctctgagggagtgcagcactcgcccgtgtctctcccacctcaccctaacaccgggatgcaaaacaaaaaccccacagttcgctctcttccgaaacggaaaatgaaaatg
gccgggaggagacggagtgtgtggggggggggtgggggggggggggggggggggtggggggggtgggggcgggggttctccctccagcgaaccttgaatcaaatatccctctcccggaaagttcaaaaggaaaacacacacgggcgcatggtttcccagcattggtgggatct
tgctatgcagcgtggtcccacctcacgacagagaccctgtgcaaagtttgacccttacgaaatggacggttcaaagttgagcccttcctgttcacggtgaccgtcacgagcaaaggtccccgattcctgggcattttctggaatgttctcctcagaacattcgtccattttccc
cccctccccaccccaccccaccctcggtcactctcagctacgatctagccagcgaaaaccttttgcgtggagagttccctcgccaactccactgggagatgttcccaaggtgtaggtgggatttgactcccactctcctgacccagaaggagagtcattacccgtcttccacct
gagaaattctggtatgactgagtcaaacacactctactccatctcccaaccctttgttttgagtttgttatatccaactaacacctcgacgcatgccaggaatgagtgaatctgctccagaggattcaacgttgtcaggggtcaacgtccattaattgggtagggggaatggt
tgggatcctctggtgactagaagagtcaggggtgacttttgttatggactaggccagaccactcaaaacattcttaagcagacagcccagaccataactttgcaatttgtgtcagtaaaataacccagagtatgttagctaggttgaccaccaggttttaaaacagaggaaaat
ctttagattcgtttttttagattagattcccgacagtgtggaaataggcccttcggctcaacaagtccacacttattcacggagtgggaggcaaagaatccttatagaacccagtcacccccaactcaactgaatatgcacaacggtcccaataaacaaacccctttaaacaca
gtgaaaatgaaacaaaggtttacaggtcgagggtagaaggacagagagtctagcggcctgtttccacacagcccactgccgaactgactcaaacaacactccagtttctaggactgactactcctctttcattatacagttcacgaccaaaacatggccactttggcttgaggt
ctcatcggtttacagatgaacagaaggcctctcaaaatcctttctcatctctgtaccaaactggccacctcggagctgggagctggtttacggcccctctgaaaacaaaaccaagtgccattgagaaaagaagcagctttgtgacactttattaaaacatatcggctcctgagg
gacacctgactggctggcggtggagggaatgctgggagaatctgaaacaaaggcattcctggttaaaatcaagggggcggccattttaaccagagctgagggagcatttttctttctctgagaggcattgtgagtctgtggaatatcccgtcctccacaggccgtggatggaga
aactttaagcgttttgaaggagaggtagattgaatctcaacgaccaatcagaagtgaggaattgtcggcggggaaggatgtgcggtggagatgtaggaatggagacctgatgttgggatcagatcaaatgcgatctcattgaagggcacagcaggttgacgagtaccaattgtt
acagaggtaggaggtttcctacctgtctccctttgaaccccccaatttttttctaaaacgcctcaacaggaccactctcccatcgcccctgacagcccacggtaatccaaccagattttgagctctcagccacttcaggatctctctggtggatccatggcacaccctttccat
aaggccagactggagtaagttcttaaattggcattgtgttgtctgttatggctctgtggttagcactgcttcctcacggtgctagggacccagattcgattccaccctcggggcgactgtctgtgtggagtttgcacgttctccccgtgtctgagtgggtttcatcacggtgct
ccggtttccttccacaaaccaaagatgtgcaggtcaggtaaattggccatgctcaattgtcccataatgcccagggatgtgcaggttagggtggattggccgtgctaacttgtcccatagtgcccagagatgtgcaggttagggtggattggccatgctaaattgtcccatagt
gcccagagatgtgtaggttagggtggattggccatgctaaattgtcccatagtgcccagggatgtgcaagttaggatggattggccatgctaaattgtcccatagtgcccagagatgtgcaggttagggtggattggccgtgctaacttgtcccatagtgcccagagatgtgta
ggttaggatggattggccatgctaaattgtcccattgtgcccagagatgtgtaggttaggatggattggccatgctaaagtacatatggtgttcagggatgtgtcagttaggtgcattagtcagcagtaaatgttgggaaattggtttgggtgggttactctttggagggtcag
tgtgggtgtgttggatcaaatggcctgtttacgtactgtcggaattctatttgattctaaacttcaggaatctgaggatctctttgtgcgtcccacttgctaggctagcgattactggcaatccctagttgtcccagagggcaattaagagtcaaccacatcactgtgggcctg
gagtcacgtctcggccagaccgagtaagggtggcagatttccttcccaaatttaaatattcaccaaccagttgtgtgtttaacaatcagaaatggttcctatttctgttggaattaaacttttaccatctgccattgtgggatttgaactcataaccacagaatattaccctga
attattagtccaatggcagcaccactgcctgctgtgttagaggcccctagcttctcaatgaggcctccatgccattgcttcagatgtttagtttgtgctgagaaaataaatgtacctaatgtccaatatttaccacaagagcattatccattaaagaatttttcatttatttgt
taatcaactgcaaactatgtcttgctgttctgatggaaagtatatcgctcatgtatattttaatgccacttagtttggatttttggattttgaacgagtggcatctgggattaaatgattaacttgccggtatttctgtagccattgtgattacatttaaaatatcatgtgagt
gacagacttaacggagtgcaatgggtgttttattttcattggctgagttttaagatgtagctttaaaggtctggtaatgacttaattcttggctcagaatgaccacactggattgacgaaaaaccttttttggctttaggtttccagcagacttggctgaagctggagttgctcc
taaagaagggagatgggtgtagtacagttaagaaatagtgtagggaaattaacttgaatgtaccagaccagaggtatttggttgaaatcctgaatcagttctgcgaagctgagaaaatctaagctcacaaggaagaggctattaaactctctgtgccaacaattggaacccata
gttaaatcaaaccctacaagtgattgagtggggaattttctctagtatttgagtgtgtattggagtactgctgggattaaaggggtttaattttattgcatattgatagcttaactatgtgtaactttgtttactgtatgttagttttgcaataataaagttctgctctttgat
tcaaacccagtctc

### C: Sequence of cDNA (coding region in red and highlighted in yellow)

accccaaagatgcagcttgccttgctgctgctgatagtgtgcgtctccagctacaatcctgttctcatggaggatattcgggatgggaaggaagataaacccagggtggaactcagtgcaaacacaacaggcaccttggaacccggcaggaaacattgcctgtgtaaagctggt
cacctcccgttacactaccgcgcaaagcagccattcgtaacggctggcagaaagtgcttctgtcagaaacgaagccagatcaccaggaaaaagcacaaggatgctctcttcgcgagacagcgccaagggaaacgttactgcctgaaaccaggaaaggccgtgtaccaaggccag
cgacaattgtattgcgtggagagctgggccagctggagaaagtgtgccccctgtgacccggtcgcccttatccccaccccactttaagacagcgcgctgaagtatagggagctcccaatgccgcccggagactccctgcacggaccgttcggaattcccagacagcgaagccga
tcatccccggctccaccaccctccggcctccccgcgacgccgcccagtgctggacatcccccagtgcccgctctgagggagtgcagcactcgcccgtgtctctcccacctcaccctaacaccgggatgcaaaacaaaaaccccacagttcgctctcttccgaaacggaaaatga
aaatggccgggaggagacggagtgtgtggggggggggtgggggggggggggggggggggtggggggggtgggggcgggggttctccctccagcgaaccttgaatcaaatatccctctcccggaaagttcaaaaggaaaacacacacgggcgcatggtttcccagcattggtgg
gatcttgctatgcagcgtggtcccacctcacgacagagaccctgtgcaaagtttgacccttacgaaatggacggttcaaagttgagcccttcctgttcacggtgaccgtcacgagcaaaggtccccgattcctgggcattttctggaatgttctcctcagaacattcgtccatt
ttccccccctccccaccccaccccaccctcggtcactctcagctacgatctagccagcgaaaaccttttgcgtggagagttccctcgccaactccactgggagatgttcccaaggtgtaggtgggatttgactcccactctcctgacccagaaggagagtcattacccgtcttc
cacctgagaaattctggtatgactgagtcaaacacactctactccatctcccaaccctttgttttgagtttgttatatccaactaacacctcgacgcatgccaggaatgagtgaatctgctccagaggattcaacgttgtcaggggtcaacgtccattaattgggtaggggga
atggttgggatcctctggtgactagaagagtcaggggtgacttttgttatggactaggccagaccactcaaaacattcttaagcagacagcccagaccataactttgcaatttgtgtcagtaaaataacccagagtatgttagctaggttgaccaccaggttttaaaacagagg
aaaatctttagattcgtttttttagattagattcccgacagtgtggaaataggcccttcggctcaacaagtccacacttattcacggagtgggaggcaaagaatccttatagaacccagtcacccccaactcaactgaatatgcacaacggtcccaataaacaaacccctttaa
acacagtgaaaatgaaacaaaggtttacaggtcgagggtagaaggacagagagtctagcggcctgtttccacacagcccactgccgaactgactcaaacaacactccagtttctaggactgactactcctctttcattatacagttcacgaccaaaacatggccactttggctt
gaggtctcatcggtttacagatgaacagaaggcctctcaaaatcctttctcatctctgtaccaaactggccacctcggagctgggagctggtttacggcccctctgaaaacaaaaccaagtgccattgagaaaagaagcagctttgtgacactttattaaaacatatcggctcc
tgagggacacctgactggctggcggtggagggaatgctgggagaatctgaaacaaaggcattcctggttaaaatcaagggggcggccattttaaccagagctgagggagcatttttctttctctgagaggcattgtgagtctgtggaatatcccgtcctccacaggccgtggat
ggagaaactttaagcgttttgaaggagaggtagattgaatctcaacgaccaatcagaagtgaggaattgtcggcggggaaggatgtgcggtggagatgtaggaatggagacctgatgttgggatcagatcaaatgcgatctcattgaagggcacagcaggttgacgagtaccaa
ttgttacagaggtaggaggtttcctacctgtctccctttgaaccccccaatttttttctaaaacgcctcaacaggaccactctcccatcgcccctgacagcccacggtaatccaaccagattttgagctctcagccacttcaggatctctctggtggatccatggcacaccctt
tccataaggccagactggagtaagttcttaaattggcattgtgttgtctgttatggctctgtggttagcactgcttcctcacggtgctagggacccagattcgattccaccctcggggcgactgtctgtgtggagtttgcacgttctccccgtgtctgagtgggtttcatcacg
gtgctccggtttccttccacaaaccaaagatgtgcaggtcaggtaaattggccatgctcaattgtcccataatgcccagggatgtgcaggttagggtggattggccgtgctaacttgtcccatagtgcccagagatgtgcaggttagggtggattggccatgctaaattgtccc
atagtgcccagagatgtgtaggttagggtggattggccatgctaaattgtcccatagtgcccagggatgtgcaagttaggatggattggccatgctaaattgtcccatagtgcccagagatgtgcaggttagggtggattggccgtgctaacttgtcccatagtgcccagagat
gtgtaggttaggatggattggccatgctaaattgtcccattgtgcccagagatgtgtaggttaggatggattggccatgctaaagtacatatggtgttcagggatgtgtcagttaggtgcattagtcagcagtaaatgttgggaaattggtttgggtgggttactctttggagg
gtcagtgtgggtgtgttggatcaaatggcctgtttacgtactgtcggaattctatttgattctaaacttcaggaatctgaggatctctttgtgcgtcccacttgctaggctagcgattactggcaatccctagttgtcccagagggcaattaagagtcaaccacatcactgtgg
gcctggagtcacgtctcggccagaccgagtaagggtggcagatttccttcccaaatttaaatattcaccaaccagttgtgtgtttaacaatcagaaatggttcctatttctgttggaattaaacttttaccatctgccattgtgggatttgaactcataaccacagaatattac
cctgaattattagtccaatggcagcaccactgcctgctgtgttagaggcccctagcttctcaatgaggcctccatgccattgcttcagatgtttagtttgtgctgagaaaataaatgtacctaatgtccaatatttaccacaagagcattatccattaaagaatttttcattta
tttgttaatcaactgcaaactatgtcttgctgttctgatggaaagtatatcgctcatgtatattttaatgccacttagtttggatttttggattttgaacgagtggcatctgggattaaatgattaacttgccggtatttctgtagccattgtgattacatttaaaatatcatg
tgagtgacagacttaacggagtgcaatgggtgttttattttcattggctgagttttaagatgtagctttaaaggtctggtaatgacttaattcttggctcagaatgaccacactggattgacgaaaaaccttttttggctttaggtttccagcagacttggctgaagctggagtt
gctcctaaagaagggagatgggtgtagtacagttaagaaatagtgtagggaaattaacttgaatgtaccagaccagaggtatttggttgaaatcctgaatcagttctgcgaagctgagaaaatctaagctcacaaggaagaggctattaaactctctgtgccaacaattggaac
ccatagttaaatcaaaccctacaagtgattgagtggggaattttctctagtatttgagtgtgtattggagtactgctgggattaaaggggtttaattttattgcatattgatagcttaactatgtgtaactttgtttactgtatgttagttttgcaataataaagttctgctct
ttgattcaaacccagtc

### D: Amino acid sequence (signal peptide shaded)

MQLALLLLIVCVSSYNPVLMEDIRDGKEDKPRVELSANTTGTLEPGRKHCLCKAGHLPLHYRAKQPFVTAGRKCFCQKRSQITRKKHKDALFARQRQGK
RYCLKPGKAVYQGQRQLYCVESWASWRKCAPCDPVALIPTPL

**Fig. S7.** Gene position and architecture (**A**), genomic DNA sequence (**B**), cDNA sequence (**C**), and encoded protein sequence (**D**) of the unannotated *cxcl17* gene in the NCBI reference genome of *Chiloscyllium punctatum* (brownbanded bambooshark). The information was downloaded from the NCBI gene database (https://www.ncbi.nlm.nih.gov/gene).

### A: Gene position and architecture

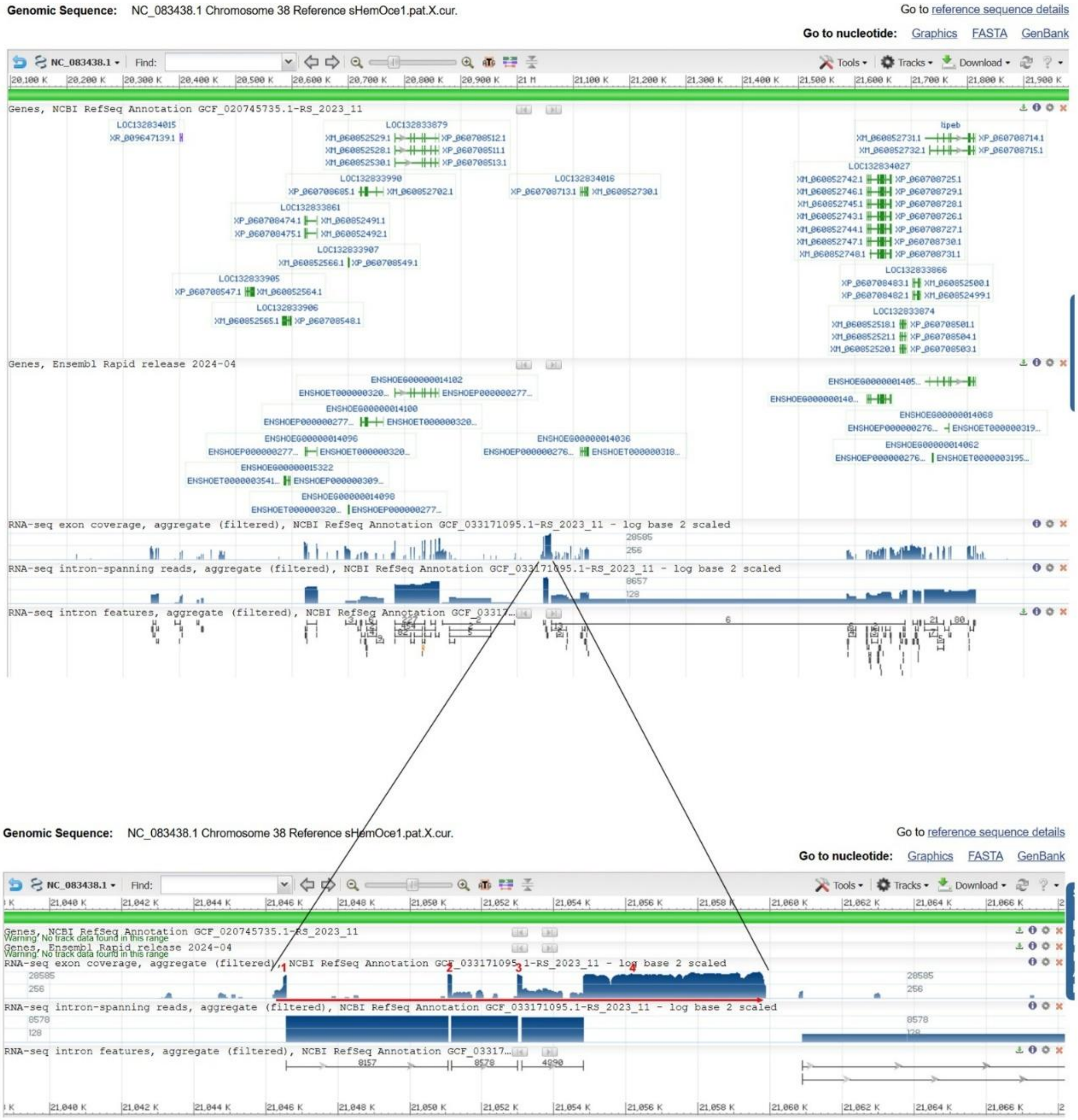


### B: Sequence of genomic DNA (exons in red and highlighted in yellow, coding region underlined)

gaatttcaggaacttgccagtaatgatgggttcacaggcagagaaattggcagcagtttctctggaggggagtggtgcaaggttggggggtggtctgttgaaccgcagagctatcagggataggcaggattgccttcaggagagatataaatcttccagcgctcagaggttcgca
cgaattagcctggggaaacatcccgaagatgcatcttgtctttctgctgctgatcgtgtgtgtctccagctacaatcctgtcctcatgcagggtaggaccaatttacaataactcagttgggtcgtcactgcctgttcacgaatgcaggacttcctgggagtagggggggaggtg
ggggatcggggaaggtgggggaggtgggggatcggggaaggtgggggaaaggggagaggggaaacgggacaggacgggaggttggagggggggtaaagggactggaggaggagggtggtaatgcccagggtggtcactggtcatggtcagcgatcagtcagaacaattatttc
cccccacacttaccgactcagggatgggacccctacttccctggagaaagtgaggtctgcagatgctggagtatcagagtggaaaatgtgttgctggaaaagcgcagcaggtcaggcagcatccaaggaacaggaaattcgacgtttcgggcataagcccttcatcaggaattc
ctgatgaagggcttatgcccgaaacgtcgaatttcctgttccttggatgctgcctgacctgctgcgcttttccagcaacacattttcaaccctacttccctcccaaccctccctgtatcgcagggcttatacacttaatgctaaggtcctagggaatgttgctgaacaaagaga
ccttggagtgcaggttcatagctccttgaaagtggagtcgcaggtagataggatagtgaagaaggcgtttggtatgctttcctttatcagtcagagtattgagtacaggagttggggggtcattttgcagctgtacaggacattggttagggcactgttagaatattgcgcgca
attctggtctccttcctatcggaaagatgatgtgaaacttgaaagggttcagaaaagattgacaaagatgttgccagggttggagggtttgagctacagggagaggctgaacaggctggggctgttttccctggagcgtcggagactgaggggtgaccttctagaggtttataa
aatcatgaggggcatgcataggataaatagacaaggtcttttccctggggtgggggagtccagaactagagggcataggtttagggtgagaggggaaagatataaaagaaacctaaggggcaactttttcacacagagggtggtgcgtgtgtggaatgagctgccagagaatgt
ggtggaggctggtacaattacagcattaaaaaggcatctggatgggtatatgactaggaagggtttggagggatatgggccaggtgctggcaggtgggactagatcgagttgggatatctgttcggcatggacgggttggaccgaagggtctgtttctgtgctgtacatctctc
tgactcaatgactcctgtagcagccgtggagattgagcagagtgaacgttgggggtcggtgcagacccaatggcttccttccaccccccggagggatttcctgcctctagttccagcagctcacctgacctttaacctctgcactgaggtactcgccatgggctgagcgaactca
aaaaggcactctgggacagagagaggaaggcggggatgggggggggagagatatgaatgaccatttgcctgcctggcgcttgatctgccctggcctcctgtaaattcgaagaccgagccagtgactcaatctgaatacaggctgtgaatcagacctggagagacaggggcagaa
atcaggccattcgggcccatctagtctgctccaccatccaatcatggctgataaagtttctcaacctcccgctttctccccctgacccttcacccccttgatactcaggaacctatccatctcagtcttaaatatactcaatgaccgggctgtcacggcctacagtggcagtga
atctcagactccccactctgtggctgaagaactctcttcttatctccattctaagtggtcttccctttactctaaggctgttcccttgggtcctcgtccctcttatcgacgggaacatcttcccaacatccactctgtccaggccactcaattagatgccccactcatccttct
aaattccatcgagtacaaacccagagtcctcaaatgttcctcatccgttaagctatccattcccgggaccattcccgtgaatctcctgtgaacctgctccaaggccagtgtatcctccctgagatatggcgcccaaaactgagaacaatcctccaaatctgacatggccacagc
ctgagggacactcagacgtccatccctccttttatattcaagtcctcccaagttaaatcacagcattcggggcggcagggtagctcagtgggttagcactgctgcctcacagcaccagggacccgggttcgattccagcctcgggtgactgtctgtgtggagtttgcacattct
ccccgtgtctgcgtgggtttcctcccacaaccaaagatgtgcaggtcagggtggactgagcatgctaaattgtcccatagtgtccagggatgtgcaggttagggtggattggccatgctaaattgtcccatagtgtccagggatgtgcaggttagggtggattggccatgctat
attgtcccatagtgacaggtgcatacgtcagagggaaatgggtctgggtgggttactcttcgaagggtcggtgtgggctggttggccgaagggcctgtttccacactgtagggaatctaatctgatctaaaaaaaatctaatagttgtatttgccttcggaatgactgactca
attgtcgacacctctttcgttatgagagtattcggaaacaagctcagacacagctataagctggagcaggaaatcttcatatcccagcaagtacagggcaaactatccatacatacacaggcagggagggctggagggggttacagagatagggagggggtgtaggggctggag
gggtcacagagatagggagggggtgtaggggctggagggggtcacagagatagggagggggtgtaggggctggagggggtcacggagatagggagggggtgtaggggctggagggggtcacagagatagggagggggtgtaggggctggagggggtcacggagatagggaggg
ggtgtaggggctggtgggggtcacggagatagggagggggtgtaggggctggagggggttacagagatagggaagggtgtaggggctggagggggttactgagatagggagggggtgtaggggctggagggggttacagagaaagatcgggaaagggatgtaggggctggaggg
ggttacagagatagggaagggtgtaggggctggagggggttacagagatagggagggagtgtaggggctggagggggttacagagatagggagggggtgtatggctggaggaggttacagagatagggagggggtgtaggggctggagggggttacagagatagggagggggtgt
aggggctggcggaggttacagagatagggagggggtgtaggggctggcggaggttacagagatagggagggggtgtaggggctggaggaggttacagagatagggagggggtgtaggggctggaggaggttacagagatagggagggggtgtaggggctggagggggttacaga
gatagggtgggggtgtaggggctcgggggggttacagagatagggagggggtgtaggggctcgagggggttacggagatagggagggggtgtaggggctggagggggttacggagatagggagggggtgtaggggctggagggggttacggagatagggagggggtgtaggggc
tggagggggttatggagatagggagggggtataggggctggagggggttatggagatagggagggggtgtaggggttggagggggttacagagatagggagggggtgtaggggctggagggggttacagagatagggaggggggtaggggctggagggggttacagagataggg
agggggtgtaggggctggagggggttacagagatagggaagggggtaggggctggagggggtgacagagatagggagggggtgtaggggctggagggggttacagagatagggagggggtgtaggggctggaggggtttacagagatagggagggggtgtaggggctggaggg
ggttacagagatagggagggggtgtaggggctggcggaggttacagagatagggagggggtgtaggggctggagggggttacagagatagggagggggtgtaggggctggagggggttacagagatagggaaatttgcagtcatcgatcaatttaaagaaaatacgaactgat
taagttgaagggttttccagatccgaacacaacttgtttgttcataattaggaacgcggcatgtggagaagataattaatgctgaatctttgctcatagaaacggatgttcagctggattcgacaaagcaatgccggggtttgggaggtagaggttgggtggaggatcaggtaa
ggttcagtggtggcacatattctcacagacattgtattcatttcagatattcggggagggaaggaagataaacccagtgtggcactcggtacaaacagaacaggaacccgccaacatggtgggaaatattgccagtgtaaaggtgggtgctcacaccagggaccgggcaaacac
agggaacggtcacaccggggagcgggcaaacacggggaacggtcacaccagggacctggtaaacaccgggaacggtcacaccggggacagggtaaacaccgggaacggtcacaccggggaccgggtaaacacggggaacggtcacaccggggactgggtaaacaccgggaacgg
tcacaccggggaccgggcaaacacagggaacggtcacaccggggagcgggcaaacacggggaacggtcacaccagggacctggtaaacaccgggaacggtcacaccggggacagggtaaacaccgggaacggtcacaccggggaccgggtaaacacggggaacggtcacaccgg
ggactgggtaaacaccgggaacggtcacaccggggacagggtaaacacggggaacggtcacaccggggaccgggtaaacacggggaacggttacaccggggacagggtaaacacggggaacggtcacaccagggaccgggcaaacaccgggaacggacacaccggggaccgggt
aaacacggggaacggtcactccagggaccgggtaaacacggggaacgatcacaccggggaccgggtaaacacggggaacggtcacaccggggaccgggtaaacacggggaacggtcacaccggggacctggtaaacactgggaacggacacaccggggatcgggtaaacacggg
gaacggacacaccggggaccgggtaaacacggggaaccgacacaccggggacagggtaaacacgggaaacggtcacaccggggacagggtaaacacggggaacggtcacaccggggaccgggtaaatacggggaccggacacaccggggacagggtaaacacggggaacggtca
caccggggacagggtaaacacggggaacggtcacaccggggaccgggtaaacacggggaacggtcacaccggggacagggtaaacacggggaacggtcacaccggggaccgggtaaacacggggaacggtcacaccggggacagggtaaacacggggaacggtcacaccgggga

cagggtaaacacggggaacggtcacaccggggaccgggtaaacacggggaacggacacaccggggaccgggtaaacacggggaacggacacaccggggacagggtaaacacggggaatggtcacaccaggcaccgggtaaacacggggaacggtcacaccggggacagggtaaa
caccgggaacggacacaccggggatcgggtaaacacggggaacggtcacaccggggaccgggtaaacacggggaacggtcacaccggggacagggtaaacacggggaacggtcacaccggggaccgggtaaacacggggaacggtcacaccggggacagggtaaacacggggaa
cggtcacaccggggaccgggtaaacacggggaacggtcacaccggggaccgggtaaacacggggaacggtcacaccggggaccgggtaaacacggggaacggacacatcggggacagggtaaacacggggaacggacacaccggggatcgggtaaacacggggaacggtcacac
cggggaccgggtaaacacggggaacggtcacaccggggaccgggtaaacacggggaacggtcacaccggggaccgggtaaacactgggaacggtcacaccggggaccgggtaaacacggggaacggtcacaccggggacagggtaaacacggggaacggtgtgttctgtccctg
ctgacaaactccctggttgaagagttcctctctttaatcgctgggccatctctgatctgcctttctcctgtagctggtctcctcccggttcactcccgcccaaagcagccaatcgtaacggctggccgaaagtgcttctgtcagaaacccagccagatccccaggagaaagcac
agggatggtaaagtgttctttttcactctgttttcggggctgagagggagggtgtccttggtaggcatcccctgagatctgtgggttcctgcgcatagcactttagtcctccatgagtctgaaagctccccatctccggatacgcccttgccttttgtgattatctgtgatggc
tttgaatcccacaccctctagctctgcagatgttatgttctgatgccaactctgatcagattctcagccaggctgtcgttatcagtgcttgccctgatgagaacataactcatttgtgtcagttaaggtggcttgataatattccgccaatgtttatctctgctccacagaaac
agaggagctggtcgtggtcacgttgctgattgctgacctcagactggggtcctgcgttacagcagtattcaacgtcccttcgatggtctgcgcccatggggttcttccaccagggggtgctgtaatcaggcgaaccacatcaatcccaatgttaaagcgggagttggggatatt
cgccaatgactgggaactgacaagtgacgggtcacattgacaacggtagcaccatccgggcgaaatccaggaattccctccctcagggcgttgtgcccacagcagagggactgcagcggttcagcccagtgtctgtgtgggtatcctccaacagtccaaagatgagcaagttag
ggtggattggccgtgctaaattgtcccatagtgcccagggatgtgctggttagggtagattggccatgctaaattacccatagtgcccagggtgtgtaggttagggtagattggccacgctaaattgtcccatagtctccagggatgtgcaggttagggtggattggccatgct
aaattgtcccatagtgcccaggaatgtgcaggttagggtggattggccatgttaaattgtcccatagtgcccagggatgtgcaggttagggtggattggccgtgctaaattacccatagtgcccagggatgtgcaggttagggtggattggccatgctaaactgtcccatagtg
cccagggatgtgcaggttagggtggtttggccatgctaaattacccatagtgcccaaggatgtgcaggttagggtagattgaccatgctaaattgtcccatagtgttcagggatgtgcaggttagggtggattggccatgctaaattgtcccatagtgttcagggatgtgcagg
ttagggtggattggccatgctaaattgtcccatagtgcccagggatgtgtaggttagggtggattggccatgctaaattgtcccatagtgttcagggatgtgcaggttagggtggattggccatgctaaattgtcccatagtgcccagggatgtgtaggttagggtggattggc
catgctaaattgtcccatagtgcccagggatgtgtaggttagggtggattgaccatgctaaattgtcccatagtcaccagggatgtgtaggttagggtggattgaccatggtaaattgtcccatagtcaccagggatgtgtaggttagggtggattgaccacgctggtatgtgt
ggatgaggtgcgtttgtcagggttcgacgtccattaattaggtagcgggggaatggttgtggattactgttgggagggtcggtgttgatatgttgggcgaaagggcctagttccttcagggattctatgagtcatgtggtgtttttaaaaatttcctttccacagctctcttc
gcgaggcagcgccaagggaaacgttactgcctgaaaccaggaaaggctgtgtaccgaggccagcgacagttctattgcgtggagagccgggccagctggagaaagtgcgtcccatgtaacccaatcgccccttatccccaccccccctttaagacggcgcgctgaagcacggggag
ctcctaacgccgcccggagactccctgcgccggccgtccggaatccccagacagcaaagcccatcatccccggctccaccaccctccggcctccccgcgacgccacccagtgctggacatcccgctctgagggtgtgcagccctcgcctgagtctctcccaccttgccccaaca
tcgggatgcaaaaaaaaatccgcagtccgctctcttccgaaaaggaaaatggaaatggtcagggaggagacggtggggggcttctccctccagctaaccgggaatgaaacatccctctcccggaacgttcaaaaggaaaactcacatgggtgcatggtttcccggcattggtgg
gatctcgctgtgcagcgtggtcccacctcacgacagagtccctgtgcaaagtttgtcccttacgaaacggacggttcaaagttgagcccacggtgaccgtcacgagcaaaggtccctgattcctgggcagcgtcatcattttctggaacgttctcctcagaacatttgtccatt
ccacctcccccccccccccccaacctctccccccccccccccccccccccccccctaccctcagtcactgtctgctacgatctagccagcgaaaaccttttaggtggagatttccctcaccaactccactgggagatgttcccaaggtgtaggtgggatttagctccaactctcc
tgacccagaagtagagtcattacccgtcttccacctgagagattctctcagagggtacgactgagtcaaacacactctcctccatctcccaacccttttgagtttgttatatccaactaacacctcgacgcaggccaggaatgagtgaatccgctccagacgaattgttagaca
gactaggccgggatcctctgatgactagaagagtcaggggtgacttttgtatggactaggccagaccactcaaaacattattaagcaggcagctcagaccataactttgcaatttgtttcagtgaaataactcagagtaagttagctaggttttaaaagagagagaaattttta
gatttgttttcttagattagaaacaggcccttcagcccaacaaatccacacttattcacggaatgaacccctatagaacccagtcaccccaaactcaactgaacatgtgaaacagtcccaataaacaaacccctttaaacacagtaaaaatgcaacaaaggtgtacaggtcaag
ggcagaaggagagagagtctagcagcctgtttccacacagcccactgctgaactgactcaaagaaaactccagtttctaggaccgactgctcctctttcgttatacatggtcactcccaaaacatgaccactttggcttgaggtctcatctgtttacatctgaacaaaaggcct
ctcaaagtcttttctcatccctgtaccacactggccacctcggagctgggagctggtttatgacccccctctgaaaaaaaaaaccaagggcccttgagaaaagaagcggctttgtgacactttgtcgaaacgtaacggctcccgaaggacccctgactggctggcagtggagaga
atggtgggagaaacaaaggcggtccatgctaaaatcaaggggggcggccattttaatcagagccgagggagcattttcctttctctgagaggcgttgcgagtctttggcatattcccgtcctccacaggccgtggatggaggagctttaagcgttttgaagggagaggtagattg
aatctcgacgaccaatcagaagggaggaattgtcggtggggaaggatcggcggtggagatgtaggaatggagacctgatgttggaatcagatcaaacgcgatctcattggaggccacagcgggtttgatgagttgcagaggtaggaggtttcctacctgtctgcctttgacccc
ccgccatttttctaaaacgcctcaacgggatcactctcccatcgtccctgtcagcccacggtaatccaaccagattttgagctctcagctacttcaggatctctctggtggatccaaggctcaccccttccataaggccagtctggagtaagttcttaaattggcattgtgttg
tctgttatggcccagtggttagcactgctgcctcacagtgctagggacccagattcgattccaccctcgggcaactgtctgtgaggagtttgcacgttctccccgtgtctgtgtgggtttcatcacagtgctccagtttcctcccacagtccaaagatgtgcgggtcaggtgaa
ttggccatgctaaattgtcccatagtgcccagggatgtgcaggttagggtgaattggccatgctaaattgtcccatagtgtccagggatgtgcaggttagggtggattggccatgctaaattgtcccataatgcccagggatgtgtaggttagggtggattggccatgctaaat
tgtcccacagtgtccagggatgtgtaggttaggtgaattggccatgctaagttacatatggtgttcagggatgtatagtttaggtgcattagtcagcagtaaatgtagggaaatgggtttgggtgggttactttttggagggtcggtgtgagcatgttgggcctaatggcctgt
ttccgtactgtaggaattctatttgattctgaacctcaggaacctggggatctcggcatgtggctctcaccaatgttgatagatgtaccacagagagaattctatctggatacatcacagtttgggattacaactgctctgtgcaggactgtgagtaactgcagaaagttgtaa
acacagcccaggccatcacactaaccaaccttccatccattgacctccatctttaattctcagtcaggcagccaatgtcatgaaagacccctcccaccctattctaatctcctccaacctcttctattgggtagaagacaccaaagtttaaagtcaggtgccaacagatttgag
aacagcttcttccccactgttattgtacttctgagtggacccctcaaatttaacttcgatcttattctctgcactccttctctgcagccataacccactctgttctcttactcccaatgccctttctatggcacaatctgcaggcaatacaaaacttttcactgtacttgggta
catgtgacagtaataaaacaaaccaagccaaatcaaatcaacctgaacgtagctactccagatagttatccaacaaagactacggccaagccttctgatctgcttgaatgctagtgttgggtcgagcccctagataggagcattctgttgcacagggagagtcctcgctctcag
ttcacagtaaagagccttttcccactaatatcatcgaatagggcacaggatgtaaatcggtgcaggggtccctgcacagagtcttaaccaaagtaaaacggaggggaatcaagggatatctctgatcgtccctgtccccaaagcccaagttgataaggagggacagagagacgc
cctactgtttctaatggaagggaagggttggattcccgtcagtatcacgttgaccagtgagggggtacctttggctaccctcactggaacggatgtcagtttattctctctctgtggatttcgatgtggccatcaaagcccagaattggctgctcatcgctggtggcccctttgt
gagttcaggagtcattcagttggcataactttgacccagccaacacagtaccaacttcaaatggcacaccggcttctgtcatctctttgtgggatgggcgtcccacttactgggccagcgattactggcaatccctggttgccccagagggcaattaagagtcaactacatcac
tgtgggcctggagccacgtctaggccagaccgagtaagggtggcagattcccttcccaaatttaaatattcgccaaccagttgtgcgtttaaacaatcagaaatggtccctatttctgttggaattaaacctttaccagccgccattgtgggatttgaactcataaccacagaa
tattaccctgaattgttagtccaatggcagcaccactgcctgctgtgtgagaggcccctggtttctcaatgaggcctccatgccattgcttcagatatttagcttgtgctaagaaaataaacgtacctaatgtccaatatttaccacaagagcactatccaataaagaattttt
catttatttgttaatcaactatgtcttgttgttctgatggaaattatatagctcatgtatattttaatgccacttagtttggatttttggattttgaactagttgcatctgggattatatgattaacttgccactatttctgtagccatggtgattgcatttaaaatatcatgt
gagagacagactttagcggagtgcaatgggtgttttattttcattggctgagttttaagatgtagctttaaaggtctggtaattacttaattcttggcacagaaggagcacattggattgaagaaaaaccttttttggctttaggtttccagcagatttggctgaagttggagtt
gctcctaaagaagggagatgggtttcgtaccgttaagaaatagtgtagggaaagtaacctgaatataccaggccagaggtatttggttgaaatcctgaatcagttctgcgaagctgagaaaatctaagctcgcaaggaagaggctatcaaagtctctgtgccaacaattggaag
ccattgttaaatcaaaccctatgagtgattgattttctctagtatttgtgtgtgtattggagtactgttgggattaatggggtttaattttattgcatgttgatagcttaactatgtgtaactttgtttactgtatgtcagatttgcaataataaagttatgctctttggttca
aacccagtctccagctttgtgtgctcccgtatgggaaaacctcataaataagctatcaagccatactctcgtctgtga

**C:** Sequence of cDNA (coding region in red and highlighted in yellow)

gaggttcgcacgaattagcctggggaaacatcccgaagatgcatcttgtctttctgctgctgatcgtgtgtgtctccagctacaatcctgtcctcatgcaggatattcggggagggaaggaagataaacccagtgtggcactcggtacaaacagaacaggaacccgccaacatg
gtgggaaatattgccagtgtaaagctggtctcctcccggttcactcccgcccaaagcagccaatcgtaacggctggccgaaagtgcttctgtcagaaacccagccagatccccaggagaaagcacagggatgctctcttcgcgaggcagcgccaagggaaacgttactgcctga
aaccaggaaaggctgtgtaccgaggccagcgacagttctattgcgtggagagccgggccagctggagaaagtgcgtcccatgtaacccaatcgccccttatccccaccccccctttaagacggcgcgctgaagcacggggagctcctaacgccgcccggagactccctgcgccggc
cgtccggaatccccagacagcaaagcccatcatccccggctccaccaccctccggcctccccgcgacgccacccagtgctggacatcccgctctgagggtgtgcagccctcgcctgagtctctcccaccttgccccaacatcgggatgcaaaaaaaaatccgcagtccgctctc
ttccgaaaaggaaaatggaaatggtcagggaggagacggtggggggcttctccctccagctaaccgggaatgaaacatccctctcccggaacgttcaaaaggaaaactcacatgggtgcatggtttcccggcattggtgggatctcgctgtgcagcgtggtcccacctcacgac
agagtccctgtgcaaagtttgtcccttacgaaacggacggttcaaagttgagcccacggtgaccgtcacgagcaaaggtccctgattcctgggcagcgtcatcattttctggaacgttctcctcagaacatttgtccattccacctcccccccccccccccaacctctcccccc
ccccccccccccccccccctaccctcagtcactgtctgctacgatctagccagcgaaaaccttttaggtggagatttccctcaccaactccactgggagatgttcccaaggtgtaggtgggatttagctccaactctcctgacccagaagtagagtcattacccgtcttccac
ctgagagattctctcagagggtacgactgagtcaaacacactctcctccatctcccaacccttttgagtttgttatatccaactaacacctcgacgcaggccaggaatgagtgaatccgctccagacgaattgttagacagactaggccgggatcctctgatgactagaagagt
caggggtgacttttgtatggactaggccagaccactcaaaacattattaagcaggcagctcagaccataactttgcaatttgtttcagtgaaataactcagagtaagttagctaggttttaaaagagagagaaatttttagatttgttttcttagattagaaacaggcccttca
gcccaacaaatccacacttattcacggaatgaacccctatagaacccagtcaccccaaactcaactgaacatgtgaaacagtcccaataaacaaacccctttaaacacagtaaaaatgcaacaaaggtgtacaggtcaagggcagaaggagagagagtctagcagcctgtttcc
acacagcccactgctgaactgactcaaagaaaactccagtttctaggaccgactgctcctctttcgttatacatggtcactcccaaaacatgaccactttggcttgaggtctcatctgtttacatctgaacaaaaggcctctcaaagtcttttctcatccctgtaccacactgg
ccacctcggagctgggagctggtttatgacccccctctgaaaaaaaaaaccaagggcccttgagaaaagaagcggctttgtgacactttgtcgaaacgtaacggctcccgaaggacccctgactggctggcagtggagagaatggtgggagaaacaaaggcggtccatgctaaaa
tcaaggggggcggccattttaatcagagccgagggagcattttcctttctctgagaggcgttgcgagtctttggcatattcccgtcctccacaggccgtggatggaggagctttaagcgttttgaagggagaggtagattgaatctcgacgaccaatcagaagggaggaattgtc
ggtggggaaggatcggcggtggagatgtaggaatggagacctgatgttggaatcagatcaaacgcgatctcattggaggccacagcgggtttgatgagttgcagaggtaggaggtttcctacctgtctgcctttgaccccccgccatttttctaaaacgcctcaacgggatcac
tctcccatcgtccctgtcagcccacggtaatccaaccagattttgagctctcagctacttcaggatctctctggtggatccaaggctcaccccttccataaggccagtctggagtaagttcttaaattggcattgtgttgtctgttatggcccagtggttagcactgctgcctc
acagtgctagggacccagattcgattccaccctcgggcaactgtctgtgaggagtttgcacgttctccccgtgtctgtgtgggtttcatcacagtgctccagtttcctcccacagtccaaagatgtgcgggtcaggtgaattggccatgctaaattgtcccatagtgcccaggg
atgtgcaggttagggtgaattggccatgctaaattgtcccatagtgtccagggatgtgcaggttagggtggattggccatgctaaattgtcccataatgcccagggatgtgtaggttagggtggattggccatgctaaattgtcccacagtgtccagggatgtgtaggttaggt
gaattggccatgctaagttacatatggtgttcagggatgtatagtttaggtgcattagtcagcagtaaatgtagggaaatgggtttgggtgggttactttttggagggtcggtgtgagcatgttgggcctaatggcctgtttccgtactgtaggaattctatttgattctgaac
ctcaggaacctggggatctcggcatgtggctctcaccaatgttgatagatgtaccacagagagaattctatctggatacatcacagtttgggattacaactgctctgtgcaggactgtgagtaactgcagaaagttgtaaacacagcccaggccatcacactaaccaaccttcc
atccattgacctccatctttaattctcagtcaggcagccaatgtcatgaaagacccctcccaccctattctaatctcctccaacctcttctattgggtagaagacaccaaagtttaaagtcaggtgccaacagatttgagaacagcttcttccccactgttattgtacttctga
gtggacccctcaaatttaacttcgatcttattctctgcactccttctctgcagccataacccactctgttctcttactcccaatgccctttctatggcacaatctgcaggcaatacaaaacttttcactgtacttgggtacatgtgacagtaataaaacaaaccaagccaaatc
aaatcaacctgaacgtagctactccagatagttatccaacaaagactacggccaagccttctgatctgcttgaatgctagtgttgggtcgagcccctagataggagcattctgttgcacagggagagtcctcgctctcagttcacagtaaagagccttttcccactaatatcat
cgaatagggcacaggatgtaaatcggtgcaggggtccctgcacagagtcttaaccaaagtaaaacggaggggaatcaagggatatctctgatcgtccctgtccccaaagcccaagttgataaggagggacagagagacgccctactgtttctaatggaagggaagggttggatt
cccgtcagtatcacgttgaccagtgagggggtacctttggctaccctcactggaacggatgtcagtttattctctctctgtggatttcgatgtggccatcaaagcccagaattggctgctcatcgctggtggcccctttgtgagttcaggagtcattcagttggcataactttga
cccagccaacacagtaccaacttcaaatggcacaccggcttctgtcatctctttgtgggatgggcgtcccacttactgggccagcgattactggcaatccctggttgccccagagggcaattaagagtcaactacatcactgtgggcctggagccacgtctaggccagaccgag
taagggtggcagattcccttcccaaatttaaatattcgccaaccagttgtgcgtttaaacaatcagaaatggtccctatttctgttggaattaaacctttaccagccgccattgtgggatttgaactcataaccacagaatattaccctgaattgttagtccaatggcagcacc
actgcctgctgtgtgagaggcccctggtttctcaatgaggcctccatgccattgcttcagatatttagcttgtgctaagaaaataaacgtacctaatgtccaatatttaccacaagagcactatccaataaagaatttttcatttatttgttaatcaactatgtcttgttgttc
tgatggaaattatatagctcatgtatattttaatgccacttagtttggatttttggattttgaactagttgcatctgggattatatgattaacttgccactatttctgtagccatggtgattgcatttaaaatatcatgtgagagacagactttagcggagtgcaatgggtgtt
ttattttcattggctgagttttaagatgtagctttaaaggtctggtaattacttaattcttggcacagaaggagcacattggattgaagaaaaaccttttttggctttaggtttccagcagatttggctgaagttggagttgctcctaaagaagggagatgggtttcgtaccgtt
aagaaatagtgtagggaaagtaacctgaatataccaggccagaggtatttggttgaaatcctgaatcagttctgcgaagctgagaaaatctaagctcgcaaggaagaggctatcaaagtctctgtgccaacaattggaagccattgttaaatcaaaccctatgagtgattgatt
ttctctagtatttgtgtgtgtattggagtactgttgggattaatggggtttaattttattgcatgttgatagcttaactatgtgtaactttgtttactgtatgtcagatttgcaataataaagttatgctctttggttcaa

**D:** Amino acid sequence (signal peptide shaded)

MHLVFLLLIVCVSSYNPVLMQDIRGGKEDKPSVALGTNRTGTRQHGGKYCQCKAGLLPVHSRPKQPIVTAGRKCFCQKPSQIPRRKHRDALFARQRQGKRYCLKPGKAVYRGQRQF
YCVESRASWRKCVPCNPIALIPTPL

**Fig. S8.** Gene position and architecture (**A**), genomic DNA sequence (**B**), cDNA sequence (**C**), and encoded protein sequence (**D**) of the unannotated *cxcl17* gene in the NCBI reference genome of *Hemiscyllium ocellatum* (epaulette shark). The information was downloaded from the NCBI gene database (https://www.ncbi.nlm.nih.gov/gene).

### A: Gene position and architecture

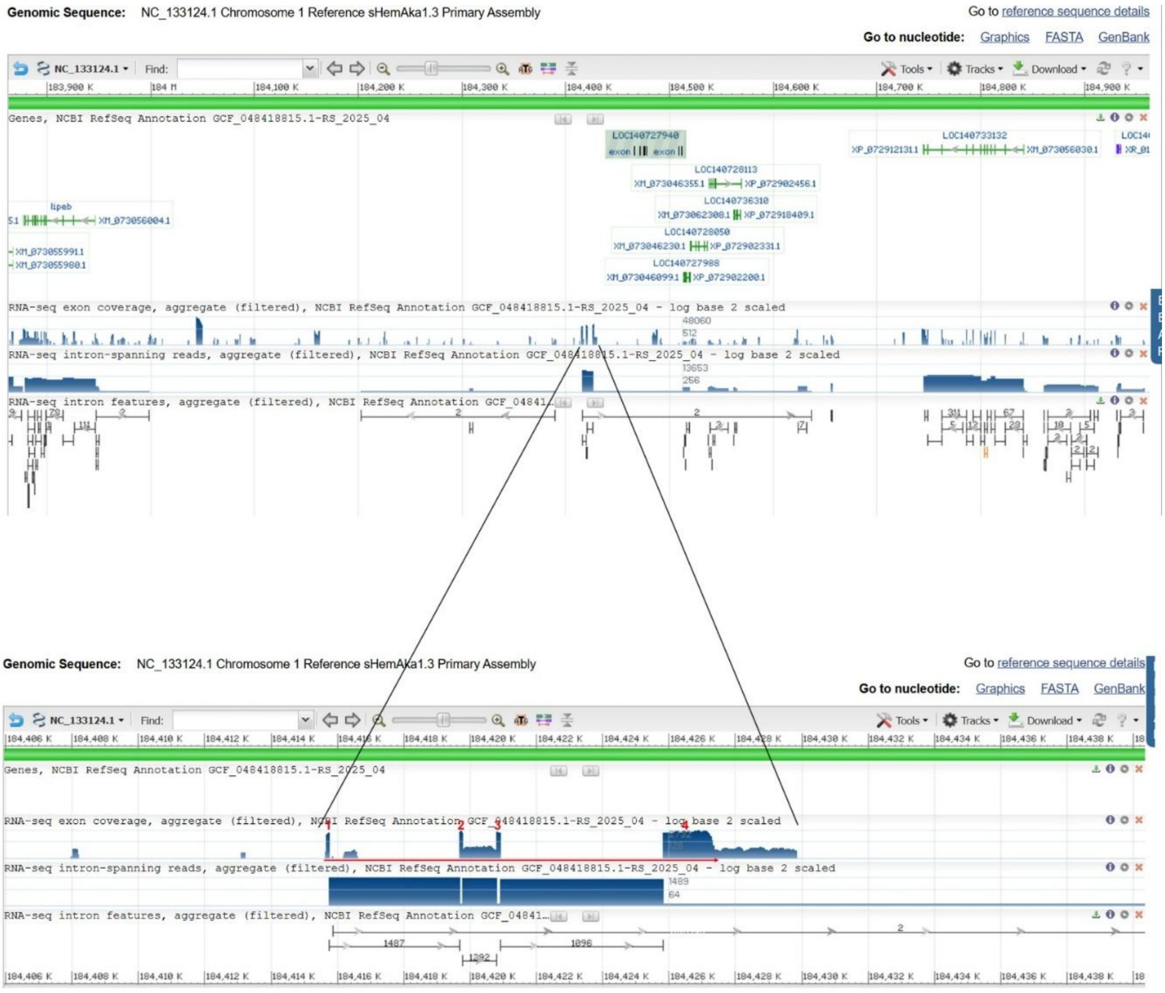


### B: Sequence of genomic DNA (exons in red and highlighted in yellow, coding region underlined)

cccacacaactcagttcaagtctgagcaggaataaaactgtcagggatacgcaagacagcacagaggagggatataagtcccacattcagagatctgggttcagctcatgatcacagcacctccgaagatgcagttttctacctggctgttcgtctttgccttagtcataattt
ctgcatttccaggaggtaagaccaccaaacatttgcaatttttttatttaatcccatgcttagaaaaaaaattgtatttattcagcagcattgctttaaaagtgaattttatattcacctgtgaggggtgaacattgttcaaaatattatttctccttccttctgtgtatcggaa
gggcttacttagccccaacagaaaaccacctcatgtgcaaatctggacaaagtgggcttggtaatagatcgtcccttaaagtcatttataaatcagttagctgctcttttgggaattcatcagccacaggaccatcttctatgacttcattattttaactgctaggctcagtaa
aatgaatttaatttaccaattagatagatagatagatagatagatactttattcatccccatggggaaattcaactttttttttccaatgtcccatacacttgttgtagcaaaactaattacatacaatacttaactcagtaaaaaatatgatatgcatctaaatcactatctc
aaaaagcattaataatagcttttaaaaagttcttaagtcctggcggttgaattgtaaagcctaatggcattggggagtattgacctcttcatcctgtctgaggagcattgcatcgatagtaacctgtcgctgaaactgcttctctgtctctggatgttgctatgtagaggatgt
tcagagttttccataattgaccgtagcctactcagcgcccttcgctcagctaccgatgttaaactctccagtactttgcccacgacagagcccgccttccttaccagcttattaagacgtgaggcgtccctcttcttaatgcttcctccccaacatgccaccacaaagaagagg
gcgctctccacaactgacctatagaacatcttcagcatctcactacagacattgaatgacgccaaccttctaaggaagtacagtcgactctgtgccttcctgcacaaggcatctgtgttggcagtccagtctagcttctcgtctaactgtactcccagatactattggaggagt
tagattcaacttctccaagttactcactcagtgatgtcaaactagtgaggaagatttatatttgtttagtgattttcatgagatcagaacatcacgactaattaattatcttttgtaaaaagacagtgaagctactgtatgcaggcagcagtttctataagaggtaacgtgcac
tgcggaacaaatcttaaacaaaactttgggaaggattcccctgctcttctttcagataatgtctctgaggtctatcatgactacccaagggcaaatggggcttcttaatgttatctgagagtgacattccctcagtatggcactgtctgatagcctgtcctagtccaactatgt
agatgccatgagtaagaaagtacaccagcacctctacctcctctggaggctaaataaatttaacatgtctcatgaaccctcaccagtttttgtagatgcaccatagcaagcctcccatctaaataagatggaaactgctttgcctgagactgcaggaaactgcagagcattgtg
aatgcttctcagcacataattcaacctctgcttcatgaactctgcctgcagttcatgctgcttttggaaagctaaatcttcccaccttagacactctctattctccctcccatcaggcagaagatacagaatttcaaagcacttactaccaggctcaaggacagcttctacccc
actgttataagactactgaatagaccccttgtatgataagatggactcttgacctcacatctacttcatcatcagcttgtacctcattgtcttcatgcattgcatgctctctgtaactgtaatactttattctgcactgttatttattttaccatgaactagctcaatgtactt
ataagatgaaataatctgtaagaaaaaaaatctgcggatgctggaaatcaaagcaatacacacaactaactcagataagtgtgaggtggctcattttggtaggtcaaatatgatggcagaatatagtattaatggtaagactcttggcagtgtggaggatcagagggatcttgg
gttccaagtccataagacgctcaaagcagctgcgcaggttgactctgtggttaagaaggcgtatagtgtgttggtcttcattaattgtggaattaaatttaggagccaagaggtaatgatgcagctgtataggaccctgggcagacaccacttgggagtactgtgctcagttat
ggtcgcctcactacaggaaggatgtggaaaccatagaaagggtgcagaggaaatttacaaggatgttgcctggattggggagcatgccttatgataacagcctcggccttttctccttggagcgacagtggatgagaggtgacctgacagaggtgtagaagatgatgagaggca
ttgattgtgtggatagtcagaggctttttcccagggctgaaatggttgccacaagaggacacagatttaagatgctgggaggtaagtacagaggagatgtcaggggtaacagagcgctgaatgcatggaatgggctgccagcaacggtggtggaggcagatatgatagggaatg
taacacttccctagcgttgtttttattaatactgcaccactcctttagtataccactgccacacaacacttcctctacactccctctgaaagtactgtgagtacttgtgtttctgatcatggctttttccacacagctcctgcttgtgcatgactactcactttaccctgtggc
actctctctgttctgcatctcccaagtcccttagtactacagtcccaacagtgcagtgttgtttggtggtggaagcaagtgccaaatttaagaagtatagtatagaaggttagagataaaattctgatcaataggaattgtatacatgggtactcaatgaccagcattgacaca
gtgatctgaaaggcttgattccatggtgtatgataatgattctatttcctttcaactgatttttgacagcttggcactacctcaaggcaacactgaaatattcctttgacatctgtagctaaattggagccacatccaaaacatttgatttgggaaccctggctaactgcccag
acactgtcatctcgccaccttgaaaatttgatttgaaaattcaagcgcattgcaggaaacaagagtatactgatgagttagggaatcatgcttatttattacaaccttcattcctgtcataaggagttttgtgcaccaaagctgggctgtgtgttgaacaggtactattaaggg
aagacttagtcaggcagttaatgaaaggctttgacaaaatagaagaggcaaacctgaatcagtggcaggtgaaaaccaaaaccacattggtttgcaataagtagcagcagttccagaggaaatgatttttaaacaacaagctaatttccaccctgcaataaacagagtacaatc
aaagggaattagcataaacagggtaaggaaaagcttgtaagactaaggaggaagcccattcaaacaatcaacataagcaaaaaaaaagccaataactttgagtacaatacgacccttaaaggtaaagtttcaaacaactgatgtgattttcttacaggcctaagaatcctctag
ataaaggactacaaactcaatagcttcctgggttaaacctaacactctgggcactccgagatctctcagcttatgacaatgagttccaaagtccctcctagaattgaacaaagaattgctaaatgaaaccatgaagactggggaaattcttagttccacatcacaaggatacat
ttctaccaaaccctttcaacctcccctccttcaactcttttatagatttacaactccatcacagacatatgagcattgtaaaaaacagatttttaacttttacaaataaatcttatatttcagatgccacagaactggaggaaaagaagaaagaaaccaataagaaggagtcca
ggaatgatgaaaacaactgtgcctgtaaaggtaaatctgtaacttaatatcacagactcagtgatacagcgacaaagaaaagttcaagcagttaatccagagatagtcaaagcagaggaaagagaaaccaatgcatctgctggaaaaaactcagcagtttaagcaaatgtcaac
gaattgggtgaaaaagaaaaacatgcattacgactgatcatcaggcttctaatgcagggtccaacgtaaaacatcgacaatttcttttctccctcatttatcagttgatccactgagttgattccagcagatagtttgttgttgcagatttcagcatctgcagtctcttgtgtc
tccagtatctgcagatgctggttacacatggaaactgtcaaaataaagtcaaacaggagaattgtcaaaccagggatcggacaaatgtggagaatatgcatgtcgaccaccaatttcaaggcaacccataatcacattttattttcccacttccacaaacttctggcagattct
cacttagacaagtctaaaactagagggaataggtttaagatgataggggaaaggaaggcaaggttttacacatggaaggtagtaggtatatggaacaagcagttagagctcaggtaggaattttggctagcatggaagtgtcagaatgaatggcctgctttaggacaatttgta
atagccagttagtctaccaatccatgtttttgaatatgtggaaaaagctaaagaaccgggaagaaacccatgaggtcacagagagaacatgcaaactccacataatcaggatggtcagaatgaatgcaggctgctggatgggattgagccaacaagggagcagtgaaaccaggt
gcaagtaaacaaaagacaggtcaaaacaggtattggataaacaagacaaatgtagaaatggaaagtacagatacagaaccaacagaagaagaaacacgacaggtacgtcttccactgcagggcagcttctgccttcactacttcaatatttctatcttcactgcatcatttatt
aatggaaacaatctggttccatacagacactcactctgaccccggattgagcatgcagcgactccaagatggaattgagaaaaagtgtaactgtcaggaggttaaaggagtcagcaagaagaaatcgagtgcaggtaatattagaaagtcacaaacactgtgtatacatcaatg
gagacacaacaaagaagagtgcgtggaatttcagattttgtacttaaagttttcagagtttactatctctgaagtagcttcaccatgattggtattggtttattattgtcacatgtacctacatacaaatgtgctcccataacattgttaaattcaagaagtccaatgtataaa
aagcagaaccattgatttcattcagaggcatagggcaatacagtacaatacaggccgtttggcccacgatgccatgctgacctttaaatatatttcaaatttaatctaacccttccctaccagatagcccttcacttttctttcatccatgtgacttctttcatccaagagcct
cttaagtatccttaatatcacctctgcatcacgttccaagcattcacacttccctaagaaaacctacctaacaacccccctacacgttcctccaaacatcttaagattgtgttccctaatagttattaacttcagagggcataacttcttcagaatggggaggaaactggcattc
ctgctgaaaatctttatgacaatggacatgctggcagaagaattggggagggggggtaggttaccggtagaggttgaagggaatgctgtgtaatcttctatctgttgtgctctctgaaatggtcaacaaaatgaggtcatttccatcaattcctcaggggggcataaaccagggat
tgagagtatcctcatcaatttctaccatgtgctgagactacatacctacagccactggagagagtaacaagaatagattaggtataagaccactgtcatttaattacaaatataaatgggacaaagaataataaaatcatatcagttgacttagtacttgtcatgggaaactcg
ctctaagccattgcaaaatatactctcaggacatacagtatgaagcaattaactgaaattaatcaggcaaagttagtaaggtgttgtgaaaagaagatcatttgccaaccccccccaaattacacccatgtgaccagttaagctattaacctgttaacctgcacctctttggaat
atgtggagaaaccggatcaccctgaggaatcccaggtagtcacgggcaaaacatacaaattccttacagacattagcgagaattaaatcaagattgttggtgctaccatagcattatactgaccactacacaactatgctgctcctaccctgctgagacactcctcactacata

cttacatgttgcaaaaactaatgttggaaatgctgagtagtcaaaggtcttcgtatatattggtaccaatgacatggttaggaaaagtgaggaggtcctgaaagagagattttagagagctgggtggaaagctgaaaggcaggtcctccatgttagtaatctttggattgttaa
ctgtaccatgtgccagtgacagaaagaataggagcatttggcagagaaatgcatggctaaggaactggcacagagggaaggggttcagatttatcagccattggtatctcttccggggtaactacgacctgcagaaaggtgatggattgtgcctaaccccaagggggggaacca
atgttcttgtgagatggttagctagagctgtttgggagaatatgaactaattggcaaggtgggtaataaccagaatgatagggctgaggatggaacagctggtttataaacagagacagtgtgagactgctagcaaggatagaccgatattagggcaaaattgcagtcaactag
acgatctgcaatgtagcaatgtaagagagggacaaaatcgaaatggctgatgaatacagaactgaaggtgttatatggaatttatgcagtttacagaataaagtacatgaacttgtcacacagttagaaattggcacatatgatgttccaggcatcactgaatcgtgaatgaaa
ttcaaggatacatattgtatcgcaaggacaagcagggaggaaaacacccttctgataaaaaatgaaatcaaagccttagaaagaggtgacatagtagaattgttgtggatggagctaagaaactgcaagggttaaaagtccctagttatatacagacctccaaacagtaagaaa
gatatgggctatgaattacaatgtgcaatagaaaagacatctcgaaatgttatggtaggcacgggggatttcaaaatgcaggtagattgggaaaatcaggttggtgctggatcccaagagggggattttgtaaaaggccaacatatagcttttaaaagcagcttgtggttgagc
ccagcaggtgatcagctattgtggattgtgtgtggtgcaataaaacagaattgattagaaagtttaagttaaagaagcccttaaaggacagtaatatggcagaatttaccctgcaattggaaagaaatatgctaaagtagaggtatcagtattacagtggagtaaagggaatta
cagaggcataagagatgagctggccaaagttgattggaaggggacactagcaggatggcggcagagcaggctggcatttctgggagcaattcagaaagtgtaggatagataaatcccaaagatgaagaagtattataaaggcaggatgatgccattgttgctgacatgggaagt
caaacccaacataaaagccaaagagagggcatgcaaaagagcaaaaattagtggcaagttagagttagttccatcattatgccttacttatttgccttctgttggtttgcagaagacttctcagagatatttggataagcacatagatgcaagaaagttggagggatctggaca
ttgtgtatctaggagggattagtgtttgagtgtttttgatttgcttttagctggttggcacaacattataggttgaacattataggttcctgtgctgtattgttgtatataagctaagtaataatattaaagagggtaccgaatgtttcttcagatatataaagtgtaaaaat
agaggtgagaatagatttaggactgctggaaaatgatgctggagaggtagtcatggaggcaagaaagtgggggacaaactgtccatcattttcatcagtccatggtattacaggaaagataattgtatagatagagcattggctgactggcaggaggcaaagagtgggcacaaa
ggaaaccttttatggttggctgctggtaactagtgatgttccatggggatgtgtgttgggaccgtttctttttatgttatatgtcagtgatttggatgatggaattgatggctttgtggctaagtttgggacaatatgaatataggcagaggtgcagctggtgttgaagaaaca
gggctgtggaaagatttagattaggggaacagtcaaagaagtggcagatggaatacagagttgggaagtatatggtcatgcacttcagtaaaagaaataaagttgtagattattttctaaatggaggaaaaattaaaaatctaagatgcaaagtgacttgacagccctcgagta
agattccttaaaggttagtttgcaggttgagtctgtggtgaagaaggcaaatgcaatgttagcattcagttcaagaggactagaatataaaagcaagcatgtaatgctgaggctttataaggcactgctgaggtattacttggagcattgtgagcagttttgggctccttatct
aagaatggatgtgctgacgttggagagggctcaaaggaggttcacaaaaatgattctgggattgaaaggcttgtcataagaggagcgtttgatggctctggtcctgtactcactggaatttggaagaatgatggggtgggggggggggatctcattgaaaacaataatgttgtt
ttaaaggttaaaaggttaaaggttaatttgcatgttgagtctgtggtgaaggcaaatgcaatgttagcattcatttcaagaggactagaatataaaagcaagcatgtaatgttgaggctttataaggcactgctgaggtattacttggagcatttggaagaatggtggtgtggg
ggggaggatctcattgaaaacaataatgttgaaaagccttgaaagagtggatgtggagaatgtgttttctttggtgggtgaatctaagaccacagaacacagcctcagaatagagggatatacgtacagaatggagatgaggaggagtctctttagcccgagggtggtgaatct
gtggaatttgtttccacaggcaactgtggaggcacaaatcactcggtatatttaaggtaaaggttgatagattcttgattaatctggtcatgaagggttgtggagagaaggctgaagtggggcttagagggaaatggatcagccatgatgaagtggcagagcagactcgatggg
ccaaatggcctattgctcctatatcttatggtcatactattcatacaattcactgcattgaggcataatgaggtaaagcagtaaaaatgcaaaaataagtctaagagctagagtactgggtttcacagaaaacggtacataaatgactttttttccgatataatgagtggtgtat
cagtaggattgttttggggcttcagttttacatgaatccaggacagataggaaagtagattgtgaagaggctacaaaatatataactaggtcaactgtgtgggcaaagccttgtaaatcaagtatgatgttctgcagagttctgaacaacagggaatttgggttccttatgtgt
cacattagactaatacgcagtagaagcaaatagttaggaaagctaacagaatactattaatataattactaactatgtggggtttcaatttgagaacgatcggtgccatttcctgcattttactctttcactgagtacttacactagctttctttattacagctgctgctggga
agaagagacagaagccccattacttacagagggcacgtcactgtctgataccaggaagggcccaaagaggagggataagacgctactactgcatacaaacaaaactctacctcaagaaatgcaaaccctgcaaacctgacaaatttatcccaaccccactttaaaaggaattga
tgcttcacactcctctctcggtcaaagaatttaatagggagcttcagtcattgcatgaattatccctcttgatagcattaattttgcaggaaagaaatcctacttgcccccaggaaagtcctcctggtatcagccatgcaccaacccagaaagtcaagggaccaaggctctgct
ccagtaccaaactgccctaatccaaatgcactgtgagaatgtgctaacccaccaaatgcacttagttttgcataagacaattaactgaggtcctgtttgcccccaaagatgacatgagtattccatagccatagtttgaagaaaaataaaattgtccttagtttcacaaagtaa
cattcatcttcttaacaaagacactaaaacaaaaacagaaaatgttgcaagaactcaaccagtcgaacagtatctgtggaatgagaaattagttaagtttttgggtcagggacacttccactttacacaggagctgctcgactagttggtgctcttccagcattttctgtttgt
atttcagattccgccacctgcagttttatttgttcaacattgaaagagagcttatcacaccagctgttgttctgctacttgtagcatcttaactcacgtagtttaactgagatgctatctatgagccagttgttgcagagaactacaccacagttctagcagagtaaccaacat
cctcttgtattatccttttctttctgtaggagcaggtgatagaatcctttaagtctgctgatcgcatggctgatcttccacctcaagaaccatatcccatgatcatctaacatttacaaaatttaacatctaatgtcaacacaacttgaataatcacaattcaatttctacatc
acttcaaggtagaaatctttctccacttaaagcttgagcagcccatcccttactctcaaactgtgccttttggtccaagactcttcaatcaggggtaatatcttctctgagtctagcctaatcatgttctttaagaatctggcaacttttaatgagatctcctctcatacttct
aaaacaactgttcagagggagacatgaagagggcaacagatggggccagatgtatcacctaggcaaaaaatattgactagacattacaaagagacgaaaataattctgagctcctatcagtgacttcctttccatttatatagtccacttcagactcaaaagatcatccctatt
tatctgcaagatggtgctgatgagacaatatcttaacttgcttaaggcttcctctaacaatgtacccttggaattaaatgcatattccttgtttctttgtgcttaaaaattgattattgttattattaattaaaagatattttgattaataagttgaaactgaaatggaatatg
gattttttcatcaaattgatagcataatggatttgatgcattgggtccatgataattgaaaaatggct

### C: Sequence of cDNA (coding region in red and highlighted in yellow)

gggttcagctcatgatcacagcacctccgaagatgcagttttctacctggctgttcgtctttgccttagtcataatttctgcatttccaggagatgccacagaactggaggaaaagaagaaagaaaccaataagaaggagtccaggaatgatgaaaacaactgtgcctgtaaag
acactcactctgaccccggattgagcatgcagcgactccaagatggaattgagaaaaagtgtaactgtcaggaggttaaaggagtcagcaagaagaaatcgagtgcagctgctgctgggaagaagagacagaagccccattacttacagagggcacgtcactgtctgataccag
gaagggcccaaagaggagggataagacgctactactgcatacaaacaaaactctacctcaagaaatgcaaaccctgcaaacctgacaaatttatcccaaccccactttaaaaggaattgatgcttcacactcctctctcggtcaaagaatttaatagggagcttcagtcattgc
atgaattatccctcttgatagcattaattttgcaggaaagaaatcctacttgcccccaggaaagtcctcctggtatcagccatgcaccaacccagaaagtcaagggaccaaggctctgctccagtaccaaactgccctaatccaaatgcactgtgagaatgtgctaacccacca
aatgcacttagttttgcataagacaattaactgaggtcctgtttgcccccaaagatgacatgagtattccatagccatagtttgaagaaaaataaaattgtccttagtttcacaaagtaacattcatcttcttaacaaagacactaaaacaaaaacagaaaatgttgcaagaac
tcaaccagtcgaacagtatctgtggaatgagaaattagttaagtttttgggtcagggacacttccactttacacaggagctgctcgactagttggtgctcttccagcattttctgtttgtatttcagattccgccacctgcagttttatttgttcaacattgaaagagagctta
tcacaccagctgttgttctgctacttgtagcatcttaactcacgtagtttaactgagatgctatctatgagccagttgttgcagagaactacaccacagttctagcagagtaaccaacatcctcttgtattatccttttctttctgtaggagcaggtgatagaatcctttaagt
ctgctgatcgcatggctgatcttccacctcaagaaccatatcccatgatcatctaacatttacaaaatttaacatctaatgtcaacacaacttgaataatcacaattcaatttctacatcacttcaaggtagaaatctttctccacttaaagcttgagcagcccatcccttact
ctcaaactgtgccttttggtccaagactcttcaatcaggggtaatatcttctctgagtctagcctaatcatgttctttaagaatctggcaacttttaatgagatctcctctcatacttctaaaacaactgttcagagggagacatgaagagggcaacagatggggccagatgta
tcacctaggcaaaaaatattgactagacattacaaagagacgaaaataattctgagctcctatcagtgacttcctttccatttatatagtccacttcagactcaaaagatcatccctatttatctgcaagatggtgctgatgagacaatatcttaacttgcttaaggcttcctc
taacaatgtacccttggaattaaatgcatattccttgtttctttgtgcttaaaaattgattattgttattattaattaaaagatattttgattaataagttgaaactgaaatggaatat

### D: Amino acid sequence (signal peptide shaded)

MITAPPKMQFSTWLFVFALVIISAFPGDATELEEKKKETNKKESRNDENNCACKDTHSDPGLSMQRLQDGIEKKCNCQEVKGVSKKKSSAAAAGKKRQK
PHYLQRARHCLIPGRAQRGGIRRYYCIQTKLYLKKCKPCKPDKFIPTPL

**Fig. S9.** Gene position and architecture (**A**), genomic DNA sequence (**B**), cDNA sequence (**C**), and encoded protein sequence (**D**) of the unannotated *cxcl17* gene in the NCBI reference genome of *Hemitrygon akajei* (red stingray). The information was downloaded from the NCBI gene database (https://www.ncbi.nlm.nih.gov/gene).

### A: Gene position and architecture

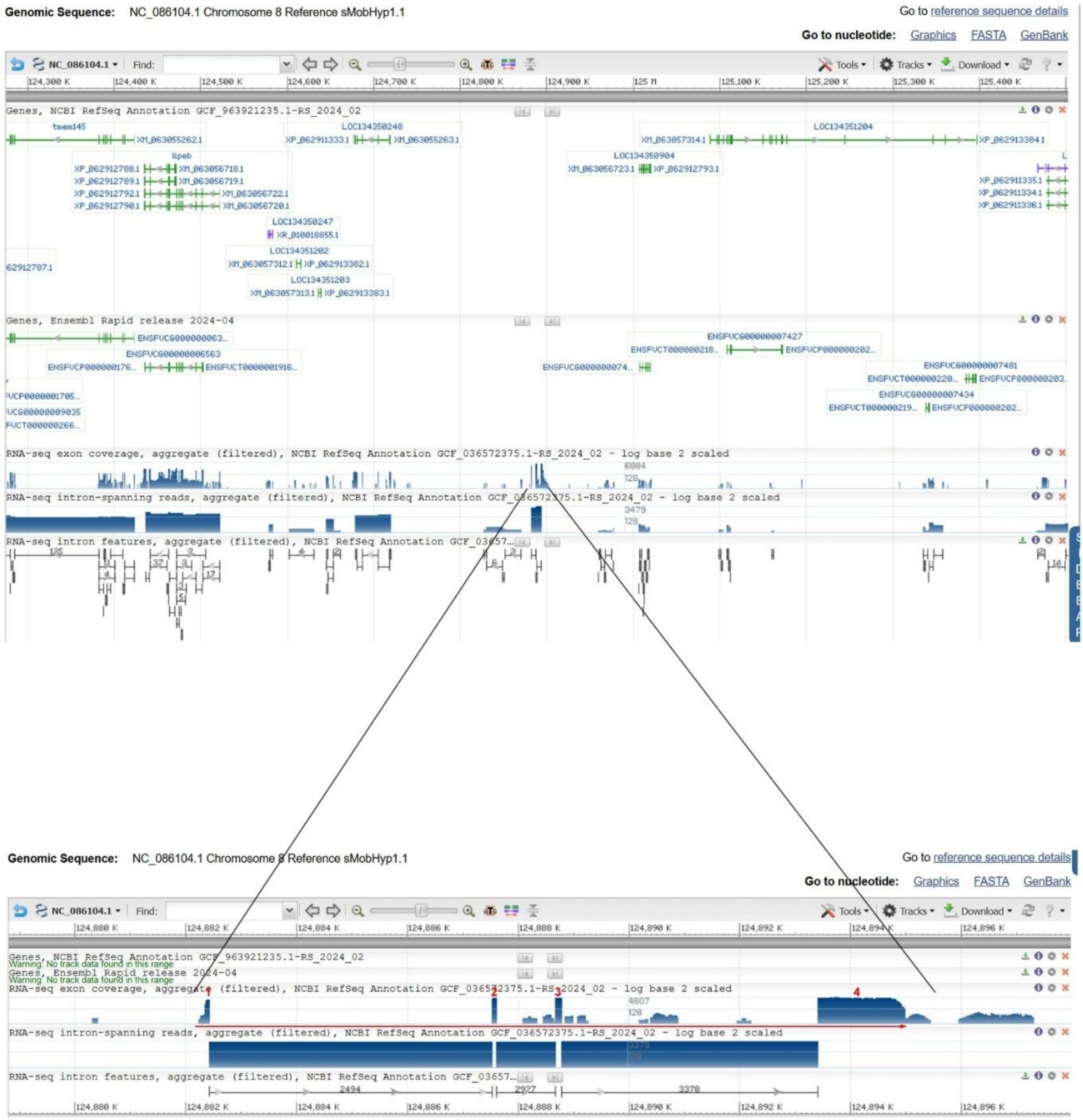


### B: Sequence of genomic DNA (exons in red and highlighted in yellow, coding region underlined)

atccctcttcgacctcatcacggaaactctgtgatggaatgagctctatcttaagccaagtcagagggtatttccactaaactcagcagatttcaggacttgttgttccttgctctcagtggctaggggatctcacagacacctcagttcaagtctgaacaggaataaaactgt
cagggatacgcaagacagcacagaggagggataaaagtcccacattcagagatctgggttcagctcatgatcacaacacctctgaagatgcagtttttacctggctgttcgtctttgccttaatcataatttctgcatttccaggaggtaagaccaccaaccttttgtgattt
ttttgtttaatcccatgcttagaaaaaagatgtatttattaagcagcagtgctttaaaaatgaattttatatgcacctgtgaggggtgagcattgttcaaaatatcagaaatgatctctccttccttctgagtatcggaagggcttagttaatcccaacagaaaaccatctcat
gtgcagatctggacaaagtgggcttggtagcagatcgtcccttaaggtcatttataaatcatttagctgctcttttgggaattcgtcagccacaggatcatcttctatgacttcattagtttaggctcagtaaactgaatttaatttaccaatttgctattgcaggagttagat
tcaacttctccaaattactcactcagtgatgccaaactaatgaggaagatttgtatttgtttagtgattttcatgagatcagaacatcatgaccaattaattagcttttgtaaaccaaaacagtgaagctatgcgggcagcagttcctataagtgcaaagaactgacactgagg
aacaaaccttaaacgaaattttgggaaagattcctccggtcttctttccaataatgactctgagctctatcatgactacccaagggcaaatggggcctcttaatgtaatctgagagtggcattcctcagtatggcactgtttgatagcctgtcctcgtccaactatgtagatgc
catgactaagaaagtacaccagcacctctacctcctcaggaggctaaataaatttaacatgtcccgttaaccctcaccagtttttgtagatgcaccatagcaagcctcccatctaagtaagatggcaactgctctgcctgagactgcaggaaactgcggagagttgtgggtgct
tctcaccactgaaattcaacctctgtttcatggactctgcctgcacttcaagctgctttggtaaaggtaattcctcccaccttagatactctctcttctccctcccatcaggcagaagatacagaatttgaaagcacttactaccaggctcaaggacagcttctatcccgctgt
tataggactattgaatagaccccttgtatgataagatagactcgacctcacatctacttcgtcatgagcttgtacctcattgtcttcatgcattgtacgttctctgtaactgtaatactttattctgcactctatttccctatgaactagctcaatgtactgatatgatgaaat
aatcggtaagaaaaaaaaatctgcagatgctggaaatcaaagcaatacgcacaacatgctagaggaactcagcaggccaggcagcatctatggaaaagagcaattttgtcaacatttcgggctgagagcccgaagtgaactgattactctttcccacagatgctgcctggcctg
ctgagttcctccagcattttgtgtgatgaaatgatctgtaagaagggtatgtaaaataaagattttcactgtaccttagtccatatgacaataaaccaattaactaatttaccctcagattcacccctcagaccacacagcacctcctaattattttccttctggctgtacagt
gttccttcaatattcccccctaaacaatgtaacactctcccagtgttgtttttattgatactgtaccaatcctttagtataccactgccatgcaacagttcctctatacattctgtctgaaagtactgtgagcacttgttttgttttgatggtggcttctcagcacagctcctgc
ttgtgaatgattcctcactttactctgtggtattctctctgctctgaatcccttagtactgcagccccaacagtgcggctttgtttgatggtggaagcaggtgccacatttaagaagcatatgtacatgcacgtgaattaccatagtatagaaggttagagataaaattctgat
caataggaattgtatacatgggtactcaatggccagcatcaacacttagagctgaaagacctgattctatggtgtatgataatgattctatttcctttcaactgattttgacagcttggcactatctcaaggcaacactgaaatattcctttgatttctgtagctagattgga
gccacatccaagacatttgatttggaaacactggctaactgcccagacactgtcatctggtcaccttgaaattttgttttgaaaattcacctgcatcgcaggaagcaagagtatagtgatgagttagtgaatcctgcttaattataacaacattcattcctgtcataaggagtt
ttgtggaccaaaactgggctgtgtgttgaacaggtactgttaagaggatacttagtcaggcagttaatgaagggctttgagaaaatagaggaggcaaacctgaatcaatggcaggtgaaaaccaaaaccacattgatttgcgataagtggcagaagttccagaagaacaaacta
aacaacaagctaattcccaccctgcaatgctcaaagttgctgtgtaaacagagtacaatcaaagggattgctaccactgctaccacccttatagttcccacaccccgcacttcccgtttctacctcctacccaagatccacaaacctgcctgtcctggcagacctattgtctca
gcttgctcctgccccaccaaactcatttctgcatacctcgacactgttttatcaccccttgttcaatcccttccgacctatgttcgtgacacttctcacgctctgaaacttttcgttgattttaagttccctggcccccactgctttattttcaccatggatgtccagtcccta
tatacttccatcccccatcaggaaggtctcaaagctctccgcttctttttggattccagacccaatcagttccccgctaccaccactctgctccatctagcggaattagtccttactcttaataatttctcctttggctcctcccacttcctccaaactaaaggtgtagctatg
ggcacccgtatgggtctgagctatgcctgctttttttgttggctttgtggaacaatctatgttccaaacctattctggtatctgtcccccactttgccttcgctacatcgacaactgcattggtgctgcttcctgcacgcatgctgagctcgttgacttcattaactttgcctcc
aactttcaccctgccctcaagtttacctggtccatttccgacacctccctcccctttctaggtctttctatctctatctctggagacagcttatctactgatgtctactataagcctactgactctcacagttatctggactattcctcttctcaccctgcctcttgcaaaaat
gccatccccttcttgcaattcctccgtctcaaccgcatctgctctcaggatgaggcttttcattccaggacgagggagatgtcctcctttttttaaaaaaaggggcttcccttcctccaccatcaactctgctctcaaacgcttctcccccatttcacgcacatctgctgtcact
ccatcctcccgccaccccactaggaatagggttcccctcatcctcacctaccaccccacctgcctccgggtccaacatattattctccgtaacttccaccacctccaacgggatcccaccactaagcacatctttccctcccctgcccctttctgctttcctcagggattgct
ccctacgcaattcccttgaccattcgttccccgcccccaccccttcccaccgatctccctcctggcacttatccttgtaagccgaacaagtgctacacatgcccttacacttcctcccttaccaccattcagggccccaaacagtccttccaagtaaggcaacacttcacctgt
gagtcgactggggtgatatactgcgtccggtgctcccgatgcagccttctatgtattggcgagacccgatgcagactgggagaccgctttgctgaacatctacactctgtccgccagagaaaagcaggatctcccagtggccacacattttaattccacgtcccattcccgttc
tggtatgtctatccacagcctcctctactgtaaagatgaagccacactcaggctggagggaacaacaccttatattccgtctgggtagcctccaacctgatggcatgaacattgacttctcaaacttctgctaatgccccacctcccccctcatacccatccattatttattta

cacacattctttttctccctgtcctttttctccctccgtccctctcactataccccttgcccatcctctgggttttccccttccccttttctttctccctgggcttcctgtcccatgatccgctcatatcccctttgccaatcacctgtccagctcttggctccatccctcc
cccttctgtcttctccaatcattttggatctcccccctcccctcccactttcaaatctcttactagctcttctttcagttagtcctgacgaagggtctcagcccaaaacgtcgactgtacctcttcctggagatgctgcctggcctgctgcattcacgagcaacttttatgtgt
gttgcttgaaattccagcacctgcagatttcctcgtgtgtgtgtttttacaatcaagcagaattagtttataaacagggtaagggaaagcttgtgagactaagaaggaaacccgctcaaactatcagcataagtcaaaagagccaaataactttaaatacaacacaaccccttaa
aggctaacctagtgtaaagtttcaaacaactgatgtgattttcttacaggcctaagagtcacctagacaaaggactacaaactcaataccttcctaggttaaacgtgacactctgggctctccaagatctctcagtttgacaatgagttccaaggtccctcctagaattgaaca
aagaattgctaaacaaaaccatgaagattggggaaattcttagttccacatccctaaacaatgacacagggatgcatttctactaaaccctttcaacttcccctccctcaactcttttatgttttgacaactccatcacagacatatcagtattgtaaaaaccagatttttaac
ttttacaaataaatctcacatttcag**atgccacagaacaggaggaaaacaagaaagaaaccaatagggatgagtccaggaatgatgaaaacagctgtgtctgtaaag**gtaagtctgtaacttagtatcacagactcaatgatacagagacaaagaaaagtggaagcagttaaat
cagagatgttcgaagcagaggaaagagaaaccaatgcatctactggaaaaactcagcagtttaagcaactgtcaacgaattgggtcaaaagaaaaacacgtataacgactgatcatcaggcttctaatgcagggtttcaacgtaaaacaatgacaaattcttttctccctcaga
tatcatttgatccactaagttaattccagcagatagcttgttgcagatttcagcatctgcagtctcttgtgtctccagtatctgcagatgctggttacacataaaactgtcaaaataaagtcaaattggggaactgtcaaaccagggatcaaataaacgtggagaatacacatg
ttgaccaccattttcatgtaaccgtcaaatacccatgatcacatttttatttccccacttccacaaacttcctgcagattctcacttagacaagtctaaaactagagggaatgggtttatgatgatagggaaaaggaaggcaaggttttacacacagaagatagctggtatatgg
gacaagcagttagagctcaggtaggaattttggctagcatggaagtgtcagaatgaatggcctgctttaggacaatttgtaatagccagttagtctactaatccatgtttttgaatgtgtggaaaaagctgaagaaccagaaagaaacccatgtggtcacagagggaatgtgca
aactccacgtagtccggatggtcagaatgaatgcaggctgctggatttataaggcaactgctatatgagggattaggccaacatgggagcagtaaaaccaggtgcaagtaaacaaaagacaggtcaaaacaggtattggataaacaagtcaaagtagaaatggaaagtacagat
acaactaaaccaatggaagaagaaacacaacaggtaggtatcccactacaaggcaggttctgccttcactacttcaatatttcaatcttccctgcatcatttattaatggcaataatctggttccatacag**acactcactctcggcttggattgaagatgcagcaactccaaga**
**tggaattgggaaaaagtgtaactgtcagaaggctaaaggagtcagcaagaaattgagtgcag**gtaatattagaaagtcacaatcaccatgtacaaatcaatggagacacaacaaagaaggatgtgtggaatttcagactttggacttaaagttttcacagtttcctatctctga
aattatcttcaccatgattggtgttggtttattattgtcacatgtacctacatacaaaggtgctcccataacgttgtcaaattcaaaaaatccaatgtataaaaagcagaaccattttgatttcattgcagaggcatagagcaatacagtacagtataggccttttggcccacg
atgccgtgctgaccttttaatctatttcatattcaatctaacccttccctcccaggtagtccttcagttttctttcatctatgtgactatctaagagcctcttaaatgtccttaatatatcggcctctgcattacgttccaagcattcacaatttcctgtgtaagaaaacctag
cagacaaccccatacatgttcctccaaacatcttaagattatgttccctaatagttattaacttcagagggcataacttctttaggatggggaggatactggcattcctgctgtaaatctttatgacaatggacacgctggcagaagaattagggtgggggtaggtcactggaa
gaggttgaagggaacgaggaaaagtgggattggtgtagattggcgtgccattcatgctgtacaatcctgtatctgttgtgctctctgaaatggtcaacaaaatgaggtcatttccatcaatttctcagggggcattgaataaggattgagagcaccctcatcagtttctaccat
gtcctgagactgcatacctacagccattggagagagtaacactaatagattaggtataggacaactgccatttaattacaaatataaatgggacaaacaaataataaaatcatatcggttgacttaatatttgtcatgggaaactcactctaagccattgcgaaagcgactctt
gggacatacaatatgaagcaataactgaaattaatcaggcaaagttagtaaggtgttgtgaaaagaagatcatttgccattcacaacccctcccccccaaattacacccacgtaactaattaagctactaacctgttaacctgcacctctttggaatatgtggagaaaccagat
ccctgaggaatcccagacagtcacgggcaaaacatacaaattccttacagacattagcgagaattgaatcaagattgttggtgctacaatagtattatactaaccactacattactatgctgctcctaccctgctgggacacgcctaactactataattacatgttgcaaaaac
taaagctagaaatggttttgagagagggtgagtagtcaaaagccttggtatatattggtaccaatgacatggttaggaaaggtgagaaggtcctgaaagagagattttagggagctaggtggaaagctgaaaggcaggacctccatgttagtactctttggattgttgcctgta
ccatgtgccagtgagagtaggaatagaagcatttggtagataaatgcatggctaaggctaaggaactggtgatcataatatggcagaatttatcctacaatttgaaagagagatgctaaagtaaaggtatcagtattacagtggagtaaagggaattacagaggcacaagagat
gagctggccaaagttgattggaaggggacattagcagggatggcggcagagcaggctggcatttctgggagcaattcagaaagtgcaggatagataaatcccaaagatgaagtattataaaagcaggatgatgccattgttgctgacaagggaagtcaaacccgacataaaagc
caaagagagggcatacaaaagagcaaaaattagtggaaagttagagttagttccatcattatgacttacttagttaccttctgttggtttttaaaagacttttcagagatgtttggataagcacatagatgcaagaaagttggagggatatggatattgtgtatcccaggaggg
attagtgtttgggtgtttgggtgtttttgatttgctttttagctggttggcacaacattataggtttaattgggttataggtttaattgtcctgacgaagggtctcggcctgaaacgtcgactgcacctcttcctagagatgctgcctggcctgccgcgttcaaaagcaacttt
tatgtgtgttgattataggtttaatggcctgtttctgtgctgtattgttctatataagctaagtaacaatattaaagagggtaccaaatgtttcttcagatatataaagtgtaaaagagaggtgagagtagatttgggaactgctgaaaaatgatgctggagaagtagtcatgg
gggcaagaaagtaggggacaaactgtccatcattttcatcagtccatggtattacaggaaagatacttgcatagatagagcattggctgactggcaggaggcaaagagtggacataaaggaaaccatttatggttggctgctgataactagtgatgttccatggggatgtgtgt
tgggaccgtttcctttttatgttatatgtcaatgatttggataatgaaactgatggctttgtggttaagtttgggacaatatgaacataggcagaggtgcaggtggtgttgaagaaacagagggcagcagaaggatttagattaggagaacggtcaaagaagtggcagatggaat
atggagttaggaagtgtatggtcatgcacttcagtaaaggaaataaaatcgtagattattttataaaaggaggaaaaattaaaaatctgagatgcaaagtgacttggcagacctggagtaagattctttaaaggttaattttcaggttgagtctgtggtgaggaaggcaaatgc
aatattagcattcagttcaagaggactagaatataaaagcaaggatgtaatgttcaaactttataaagcactggtgaggcattacgtggagcattgtgagccattttggactcctggactcgttggggagggttcaaaggaggttcacaaaaatgattctgggattgaaaggct
tgtcataaggggagcgtttgatggctctgggcctgtactcactggaatttggaagaatgatgggtggggtgggggagctcattgaaaccaataatgttgaaaagccttgatagagtggatgtggagaatatgtttcctctggtgggtgaatctaagaccacagaacacagactc
agaatagagggatgtacatttagaaaggagatgaggaggagtttctttagccagagggtgatgaatctgtggaatttgttgccacaggcgactgtggaggaaaaaaaatcactcggtatatttaaggcaaaggttgatagattcttaattaatcaggtcatgaagggatgagga
gagaaggctgatgtggggcttagagggaaatggatcagccatgatgaacagcagactcgatgggacaaatggcctattgctcctttatcttatggtcgtaccattcatacaattcactgcattgaggcataatgaggtaaagcagtaaaaatgcaaaaataagtctaagagcta
gaatactgggtttgacagaaaacatggtacataaacgtctttttttctggtataatgaatgatgtgtcagtaggattgttttggggttttagctttacacaaattcatgacagataggaaagtagattgtgaagaagctacaaaacatataattaggtcaactgtatggacaaag
ccttgcaaatcaagcatgatgttgacaatgtgaacatccattttgctcaaaataatacaaaaagcaaaaatcgaattgtgaatgcttgcagagttctgaacaacagaggaatttgggttccttatgtgtcacattagactagtatgcagtagaagcaaatagttaggaaagcta
acagaatagtatcttttgagcaacccacacaaaatgctggagaaatttagcaaggactggcccgatgaaggctctcaactcaaaatgtcgatcatttattccccttcaattatattgcccaacttcccgagttcatccagcattttgtatgtgttgctcaaaatttccagcatc
tgcagaattttgtgagagaaaggggagggatattcctgttcataatttatgtgaagagtttttcctttcatactgtgcttttctcctctataactcatcctaaagtatttgatgttgctttttgctgtaatgcaaaaagtttaattttcacacagcaagatcctcgaaacagca
agtaaaaaaaaacagataatttatttatgatataaactaagagtcaaatattgccctggttatcagggagaaatttcccactttcctttgaaccactggggagtaattttttaccttagacaaatcctcagggaaatataaaggatgccttctgatagcacaccagttggcatt
taatgagagtgggctggcagaggtcagggagatggagagatctagatttgtggaggtcaacaattcaagatataattactaattatgtggggtttaaatctgagaatgaacagtgccatttgctgcattttactctttcactgagtacttacactagcttcctttattacag**ct**
**gctgctgggaagaagagaaagaagccccattaccttcagagtccacgttactgtctgataccagggaaggccaaaaggggcggggcaagacgcttctactgcatacaaacgagactctacctcaggaaatgcaaaccctgcaaacctgtcatatctatcccaacgccactttaa**
**aaggaattgatgcccccatcttcacactcctctctcttggtcaaagaatttaaaagggagcttcagtcattgcataaattatccctcttgatagcattaattttgctggaaatacattctacttgcccccaggaaagtcctcgatattagccatgctccgacacagaaagtcaa**
**aggtccaagactctgctccagtctcaaaccaccctaatccaaatgcactgtgagagtgtgctaacccaccaaatgcacttggttttgcatgagacaattaactgaggtcctgtttgccccctgcccccaaagatgacatgagtattccatagccatagcttgaagaaaaataaa**
**gttgtccatagtttcacaaagtaacatccatctttttaacaaagacacacaaacaaaaacaaaaaatcttgcaagaactcagccagtcaaacagtatctgtggaatgagaaattagttaagtttctgggttagggacacttccactttccacaggagctgtttgactagttggt**
**gttcttccagcattttctgtctgtatttcagattctaccactcgcagttttattcgttcaacattattgagagagagcttatcacaccagttattgttctgctacttgtagcatcttaactcacatagtttaactgagatgctatctatgagccagttctctcagagaactgca**
**ccacagttctctcagagtaaccaacatcctcttgtattatccttccctttctgtaggagcaggtgacacagtcccttaagtctgctgatggcatgactgatcatccacctcaagaaccatatcccatgatcatttaacatttataaaatttaacatctaatgttaacacaattt**
**aaataatcacaattcagtttctacagcactccaaggtagaaatctttctccacttaaggcttaaatagcccatcccttactctcaaactgtgacttttggtccaagactcttcaatcaggggaatatcttctctgagtctagcctaatcatgctctttaaaaatctggcaatt**
**tttaatgaggtctcctgtcatacttctaaaactaccattcagagggaggcatgaagagtgcaacagatgcagccagatgtatcacctaagcacaaaatattgactagacattacaaagagatgaaaataggtcttagctcctatcgctgacttcctttccatttatatagtcca**
**tttcaccctatttatagccctatttatctgcaagatggtgatgagacaatatcttaacatgcataaggcttcctctaacaatatatccctggaactaaatgcatattccttgtttctttgtgcttaaaaattgatttatgttattattaattaaaagatattttgattaagttg**
**caaactg**aaatggaatatggattg

### C: Sequence of cDNA (coding region in red and highlighted in yellow)

agacacctcagttcaagtctgaacaggaataaaactgtcagggatacgcaagacagcacagaggagggataaaagtcccacattcagagatctgggttcagctc**atgatcacaacacctctgaagatgcagtttttttacctggctgttcgtctttgccttaatcataatttctg**
**catttccaggagatgccacagaacaggaggaaaacaagaaagaaaccaatagggatgagtccaggaatgatgaaaacagctgtgtctgtaaagacactcactctcggcttggattgaagatgcagcaactccaagatggaattgggaaaaagtgtaactgtcagaaggctaaag**
**gagtcagcaagaaattgagtgcagctgctgctgggaagaagagaaagaagccccattaccttcagagtccacgttactgtctgataccagggaaggccaaaaggggcggggcaagacgcttctactgcatacaaacgagactctacctcaggaaatgcaaaccctgcaaacctg**
**tcatatctatcccaacgccactttaa**aaggaattgatgcccccatcttcacactcctctctcttggtcaaagaatttaaaagggagcttcagtcattgcataaattatccctcttgatagcattaattttgctggaaatacattctacttgcccccaggaaagtcctcgatatt
agccatgctccgacacagaaagtcaaaggtccaagactctgctccagtctcaaaccaccctaatccaaatgcactgtgagagtgtgctaacccaccaaatgcacttggttttgcatgagacaattaactgaggtcctgtttgccccctgcccccaaagatgacatgagtattcc
atagccatagcttgaagaaaaataaagttgtccatagtttcacaaagtaacatccatctttttaacaaagacacacaaacaaaaacaaaaaatcttgcaagaactcagccagtcaaacagtatctgtggaatgagaaattagttaagtttctgggttagggacacttccacttt
ccacaggagctgtttgactagttggtgttcttccagcattttctgtctgtatttcagattctaccactcgcagttttattcgttcaacattattgagagagagcttatcacaccagttattgttctgctacttgtagcatcttaactcacatagtttaactgagatgctatcta
tgagccagttctctcagagaactgcaccacagttctctcagagtaaccaacatcctcttgtattatccttccctttctgtaggagcaggtgacacagtcccttaagtctgctgatggcatgactgatcatccacctcaagaaccatatcccatgatcatttaacatttataaaa
tttaacatctaatgttaacacaatttaaataatcacaattcagtttctacagcactccaaggtagaaatctttctccacttaaggcttaaatagcccatcccttactctcaaactgtgacttttggtccaagactcttcaatcaggggaatatcttctctgagtctagcctaa
tcatgctctttaaaaatctggcaatttttaatgaggtctcctgtcatacttctaaaactaccattcagagggaggcatgaagagtgcaacagatgcagccagatgtatcacctaagcacaaaatattgactagacattacaaagagatgaaaataggtcttagctcctatcgct
gacttcctttccatttatatagtccatttcaccctatttatagccctatttatctgcaagatggtgatgagacaatatcttaacatgcataaggcttcctctaacaatatatccctggaactaaatgcatattccttgtttctttgtgcttaaaaattgatttatgttattatt
aattaaaagatattttgattaagttgcaaactg

### D: Amino acid sequence (signal peptide shaded)

MITTPLKMQFFTWLFVFALIIISAFPGDATEQEENKKETNRDESRNDENSCVCKDTHSRLGLKMQQLQDGIGKKCNCQKAKGVSKKLSAAAAGKKRKKP
HYLQSPRYCLIPGKAKRGGARRFYCIQTRLYLRKCKPCKPVISIPTPL

**Fig. S10.** Gene position and architecture (**A**), genomic DNA sequence (**B**), cDNA sequence (**C**), and encoded protein sequence (**D**) of the unannotated *cxcl17* gene in the NCBI reference genome of *Mobula hypostoma* (lesser devil ray). The information was downloaded from the NCBI gene database (https://www.ncbi.nlm.nih.gov/gene).
